# Simulations probe the role of space in the interplay between drug-sensitive and drug-resistant cancer cells

**DOI:** 10.1101/2024.04.29.591633

**Authors:** Kira Pugh, Rhys DO Jones, Gibin Powathil, Sara Hamis

## Abstract

The interplay between drug-sensitive and drug-resistant cancer cells has been observed to impact cell-to-cell interactions in experimental settings. However, the role that space plays in these interactions remains unclear. In this study, we develop mathematical models to investigate how spatial factors affect cell-to-cell competition between drug-sensitive and drug-resistant cancer cells in silico. We develop two baseline models to study cells from the epithelial FaDu cell line subjected to two drugs, specifically the ATR inhibitor ceralasertib and the PARP inhibitor olaparib, that target DNA damage response pathways. Our baseline models are: (1) a temporally resolved ordinary differential equation (ODE) model, and (2) a spatio-temporally resolved agent-based model (ABM). The models simulate cells in well-mixed and spatially structured cell systems, respectively. The ODE model is calibrated against in vitro data and is thereafter mapped onto the baseline ABM which, in turn, is extended to enable a simulation-based investigation on how spatial factors impact cell-to-cell competition. Simulation results from the extended ABMs demonstrate that the in silico treatment responses are simultaneously affected by: (i) the initial spatial cell configurations, (ii) the initial fraction of drug-resistant cells, (iii) the drugs to which cells express resistance, (iv) drug combinations, (v) drug doses, and (vi) the doubling time of drug-resistant cells compared to the doubling time of drug-sensitive cells. These results reveal that spatial structures of the simulated cancer cells affect both cell-to-cell interactions, and the impact that these interactions have on the ensuing population dynamics. This leads us to suggest that the role that space plays in cell-to-cell interactions should be further investigated and quantified in experimental settings.

## 1 Introduction

The interplay between drug-sensitive and drug-resistant cancer cells has been observed to impact treatment responses in experimental settings (Emond et al., 2023; Kaznatcheev et al., 2019; Obenauf et al., 2015; Bacevic et al., 2017). This interplay is driven by cell-to-cell interactions, which are mediated by e.g., cells’ ability to send and receive signalling molecules (Armingol et al., 2021; Brady-Kalnay, 2012), and compete for resources (Tsuboi et al., 2018). In this work, we are specifically interested in interactions between drug-sensitive and drug-resistant cancer cells that compete for one such resource: space. Cancer cells can express drug resistance in several ways. Here, we focus on two broad categories of drug resistance, namely intrinsic and inherited resistance. Intrinsic drug resistance refers to resistance that exists in cancer cells or tumours before they are subjected to treatments (Wang et al., 2019). Inherited resistance occurs when drug-resistant parental cells pass on their drug-resistant traits to their daughter cells (Friedman, 2016). Cancer cells can be resistant to different types of therapies, including chemotherapy (Halacli et al., 2013), radiotherapy (Alamilla-Presuel et al., 2022), and targeted therapies, such as DNA damage response (DDR) inhibitor drugs (Baxter et al., 2022) which are the focus of this article. Cancer cells that are resistant to multiple drugs simultaneously can be referred to as multi-drug-resistant (Emran et al., 2022).

In clinics, cancer drug treatments are commonly administered at maximum tolerated doses (MTDs). However, MTDs have been associated with rapid onsets of drug resistance (Kouyos et al., 2014). As alternative treatment approaches to MTDs, drugs can be administered at low doses (i.e., lower than MTD doses) as part of single-drug or combination therapies. In fact, experimental results suggest that low-dose drug treatments can out-perform MTD strategies in certain in vitro experiments (Lloyd et al., 2020; Bacevic et al., 2017; Vallés-Martí et al., 2023). Beyond drug combinations and doses, another factor that may influence treatment responses is the spatial structure of cancer cells (Gatenby et al., 2020; Karimi et al., 2023; Sorin et al., 2023). Notably, Noble et al. 2022 recently combined mathematical modelling and spatially resolved tumour sequencing data to show that the spatial structure of a tumour can determine its evolutionary mode, or behaviour, as regulated by cell-to-cell interactions and cell dispersal. Indeed, with recent technological advances, there now exist methods that enable us to observe spatial organisations of different cell types within biopsied tumours. These methods include spatial transcriptomics (Hunter et al., 2021) and mass spectrometry imaging (Zhang et al., 2023). Although these methods reveal spatial, intratumoural cell structures, they do not explicitly provide dynamic information about e.g., the interplay between different intratumoural cell types.

One way to study the dynamics of spatially structured cancer cell populations is through simulations and mathematical modelling. Indeed, several mathematical models have been developed for studying interactions between drug-sensitive and drug-resistant cancer cells. These include two-dimensional (2-D) agent-based models (ABMs) in which each agent typically represents one cancer cell (Wang et al., 2015). For instance, Strobl et al. (2022) used such a model to study the competition between drug-sensitive and drug-resistant cancer cell subpopulations during adaptive therapy treatments. The authors showed that not only the inter-subpopulation competition (between drug-sensitive and drug-resistant cells), but also the intra-population competitions impact treatment responses in silico. Hamis et al. (2018) also used a 2-D ABM to study the eco-evolution of cancer cell populations comprising drug-sensitive and drug-resistant cells. By including different types of drug-resistant cells in their model (e.g., cells that are drug-resistant before treatments start, and cells that become drug-resistant once they are exposed to drugs) the authors showed that the mechanisms by which cells are drug-resistant affect treatment responses and optimal treatment strategies. In a third 2-D ABM, Bacevic et al. (2017) simulated a cross-section through tumour spheroids with drug-sensitive and drug-resistant cells. They showed that drug-resistant cells on the periphery of the spheroid had a proliferate advantage over drug-resistant cells in the middle of the spheroid, which they attributed to a surplus of space and oxygen resources on the spheroid’s periphery. A shared finding between the three aforementioned ABM studies is that the spatial composition of drug-sensitive and drug-resistant cells impacts cell population dynamics and treatment responses.

Inspired by these previous studies we, in this work, use a 2-D ABM to simulate the interplay between drug-sensitive and drug-resistant cancer cells in response to different drug combinations and doses. By including spatio-temporal dynamics in our mathematical model, we aim to understand how the spatial structure of cells, and the competition for space between drug-sensitive and drug-resistant cancer cells, impact treatment responses to drugs that inhibit cancer cells’ ability to repair DNA damage. We especially investigate how this resource competition affects the dynamic ratios of subpopulation sizes. Our baseline model is calibrated by data from in vitro experiments in which cells from the FaDu cell line, an epithelial morphology cell line derived from a human with squamous cell carcinoma, were subjected to two DNA damage repair-inhibitor drugs, namely ceralasertib and olaparib. Ceralasertib is an ataxia-telangiectasia mutated and Rad3-related (ATR) inhibitor that has been shown to inhibit cell cycle checkpoint activation and repair of DNA replication damage (Lloyd et al., 2020; O’Connor, 2015). Currently, ceralasertib is used in clinical phase 1 and 2 trials for treating advanced solid tumours, advanced solid malignancies, and melanomas that are resistant to PD-(L)1 inhibitors (Wang, 2022; AstraZeneca, 2022b,a). Olaparib is a poly (ADP-ribose) polymerase (PARP) inhibitor drug that has been shown to inhibit the repair of DNA single-strand breaks (Lloyd et al., 2020; Li et al., 2023). If not repaired before DNA replication, DNA single-strand breaks can lead to DNA replication stress, which in turn may cause cell death. PARP has also been shown to contribute to DNA replication fork reversal and stability, and thus PARP inhibition can lead to increased genomic instability (Chaudhuri and Nussenzweig, 2017). Olaparib is FDA-approved for the treatment of ovarian, breast, pancreatic, and prostate cancer (Food and Drug Administration, 2017, 2018, 2019, 2023). Combining ceralasertib and olaparib has demonstrated synergistic activity both in vitro and in vivo (Lloyd et al., 2020), and is being used in phase 2 clinical trials (Janeway, 2022; Aggarwal, 2019). Experimental studies have shown that cancer cells can be resistant to both ATR and PARP inhibitor drugs (O’Leary et al., 2022; Cahuzac et al., 2022; Baxter et al., 2022). Population-level implications of cell-level resistance to the ATR inhibitor (ATRi) ceralasertib and the PARP inhibitor (PARPi) olaparib are mathematically and computationally studied in this work.

## 2 Model and method

The aim of this study is to develop mathematical models to study the competition for space between drug-sensitive and drug-resistant cancer cells that are subjected to DNA damage response inhibitor drugs. As a first step towards achieving this aim, we develop a temporal compartment model to simulate how cell populations that are (a) not spatially structured, and (b) fully drug-sensitive respond to drug treatments. The compartment model is formulated by a system of ordinary differential equations (ODEs). In the compartment model, cell division and drug responses are, respectively, modelled by a cell cycle model and a drug response model (Section 2.1). As a second step, we map the compartment model onto a baseline, 2-D ABM (Section 2.2) in which all cells are fully drug-sensitive, and the total growth of the cell population size is not limited by spatial structures. The compartment model and baseline ABM are calibrated by, and evaluated against, in vitro data (Section 2.3). To investigate how spatial cell structures, specifically cell crowding effects, impact dynamics of heterogeneous cell populations in silico, we extend the baseline ABM to include spatial heterogeneity and drug-resistant cells. In this cell crowding limited ABM, drug-sensitive and drug-resistant cells are placed on the lattice in a variation of initial structures (Section 2.4). To ensure that our ABM-generated in silico results are based on enough simulations to mitigate uncertainty originating from intrinsic model stochasticity, we perform consistency analyses in the Supplementary Material (S4).

### 2.1 Temporal cell population dynamics are modelled by a compartment model

#### 2.1.1 Cells progress through the cell cycle according to a cell cycle model

We develop a compartment model that describes how cell populations that are (a) not spatially structured, and (b) fully drug-sensitive change over time in response to treatments with one or two DNA damage response inhibitor drugs. One of these drugs (drug 1) is an ATRi, and the other drug (drug 2) is a PARPi. The compartment model used in this study (Fig. 1**a**) is based on previous modelling work (Checkley et al., 2015; Hamis et al., 2021b; Pugh et al., 2023). The model is described by a system of ODEs, in which each dependent variable [*y*] describes the concentration of cells in compartment *y*. The drugs that we consider in this study target cells that are in specific phases of the cell cycle and, accordingly, each compartment represents a cell cycle phase state. We include three undamaged and proliferative cell cycle states in the model, specifically the gap 1 (G1) state, the synthesis (S) state, and the combined gap 2/mitosis (G2/M) state (Fig. 1**a**). This model is an abstraction of the biological cell cycle, where cells in the gap 1 phase grow and prepare for DNA replication, which occurs in the synthesis phase, and cells in the subsequent gap 2 phase prepare for cell division, which occurs in the mitosis phase (Israels and Israels, 2000). We also include a damaged S state (SD), representing cells that have replication stress-induced DNA damage, a type of DNA damage that results from faulty DNA replication (O’Connor, 2015; Saxena and Zou, 2022). FaDu ATM-KO cells, which are used to calibrate our model, are prone to such replication stress (Lloyd et al., 2020). A non-cycling (NC) state is also included in the model. Cells in the NC state have irreparable DNA damage and are thus unable to proliferate. Cells that enter the NC state remain there throughout the entire simulation and are not removed from the lattice. This simplifying modelling assumption is based on experimental observations of the modelled system in which damaged cells do not detach from the plates within the time frame of the experiment.

**Fig. 1:**
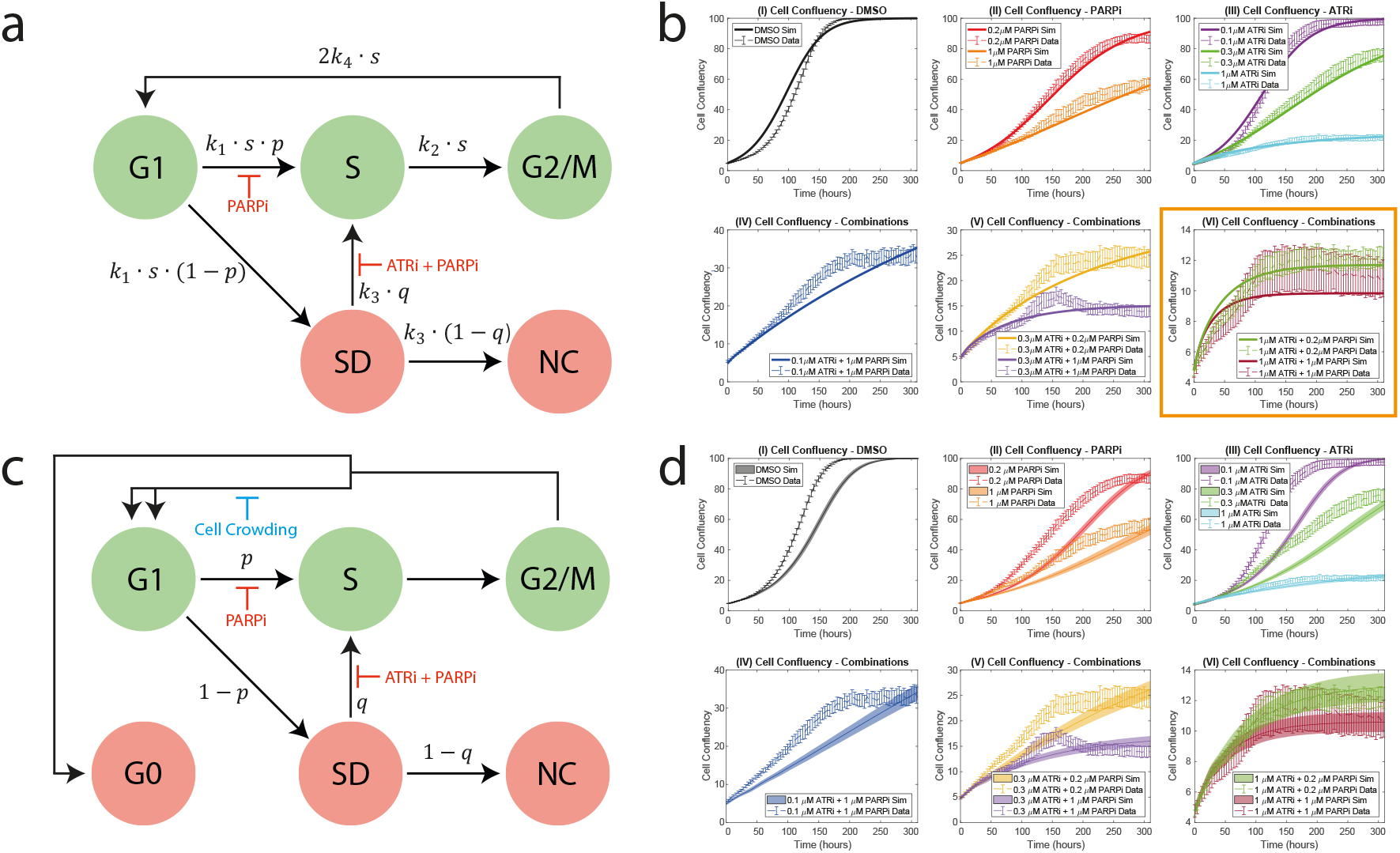
The compartment model (a) is parameterised and evaluated against in vitro data, and is thereafter mapped to an ABM (c). Undamaged and cycling cell states are shown in green nodes. Damaged and non-cycling cell states are shown in red nodes. **a** In the compartment model, cells can progress through the undamaged states G1, S, and G2/M and the damaged states SD, NC. Cells in SD experience DNA replication stress, and cells in NC have irreparable DNA damage. The paths show the transitions between states where *k*_*i*_(*U* (*t*)), *i* = 1, 2, 3, 4 are rate constants and *s* is a scaling parameter. *p* and *q* represent drug-dependent probabilities that a particular path will be chosen at a fork. The PARPi inhibits cells from progressing from G1 to S. Both the ATRi and the PARPi inhibit cells from transiting from SD to S. **b** The plots show simulated and experimental cell confluency over 310 hours for various dose combinations of the ATRi and PARPi. Training data are used to estimate model parameters (plots **I-V**) and test data are used to evaluate the model (plot **VI**, in the orange box). The solid lines show cell confluency calculated by the compartment model. The dashed lines show the mean in vitro data, with standard errors for three experiments indicated with error bars. **c** In the ABM, cells progress through the same states as in **a**. In addition, cell crowding may cause cells to enter the G0 state that represents quiescent cells. Each cell in the ABM has an individual cell cycle clock that tracks its progression through the cell cycle. **d** The plots show simulated and experimental cell confluency over 310 hours for various dose combinations of the ATRi and PARPi. The solid lines represent the mean cell confluency from the ABM with standard deviations from 100 simulation runs in opaque bands. The time and drug dependencies of the parameters in **a** and **c** have been omitted for ease of presentation.

In the model, cells progress through the cell cycle states via unidirectional paths that are associated with rates (Fig. 1**a**). These rates depend on drug concentrations, as explained in more detail in Section 2.1.2, and the total cell concentration *U* (*t*). The latter dependency is included in the model to achieve logistic growth rates of the cell populations, as are observed in the in vitro data that we use to parameterise and evaluate our model (Section 2.3). Note that after the G1 state, cells enter the S state at a rate that is proportional to a weighting factor *p*, and cells enter the damaged SD state at a rate that is proportional to 1 − *p*. The factor *p* decreases with increasing concentrations of the PARPi (drug 2), which captures the drug’s replication stress-inducing drug effect. Note also that cells in the SD state repair their DNA damage at a rate that is proportional to the weighting factor *q*, which is a function of both the PARPi and the ATRi (drug 1). As such, *q* describes the fraction of cells leaving SD that enter state S. The remaining fraction of cells that leave the SD state enter the NC state. The factor *q* decreases with increasing drug concentrations, which captures the replication repair-inhibiting drug effects of the ATRi and PARPi. Cells that successfully progress to, and leave, the G2/M state re-renter the G1 state and produce a daughter cell that is initiated in state G1 (Fig. 1**a**). The compartment model is described with the following system of ODEs

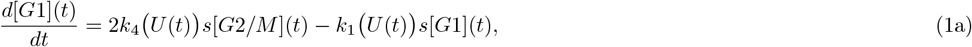

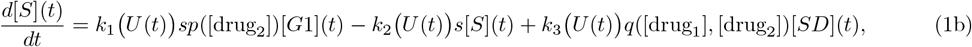

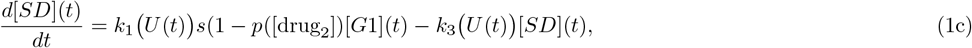

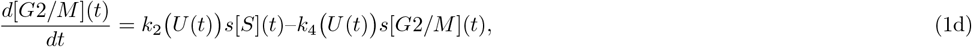

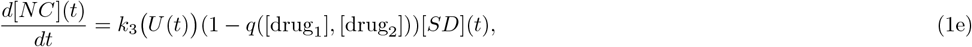

with the initial conditions

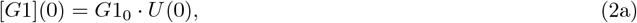

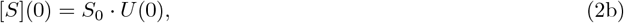

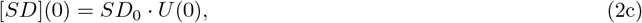

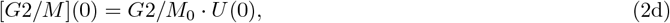

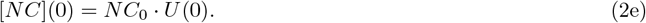

In Eqs. 1 and 2, *U* (*t*) is the total concentration of the cell population,

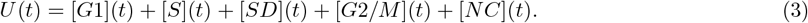

The drug effects *p*([drug_2_]) and *q*([drug_1_], [drug_2_]) are described in Section 2.1.2. To represent cell cycle arrest in our model, we assume that cell cycle progression is halted when cells are in the SD state. Therefore, to fit the model to cell doubling time data, we multiply the rate parameters *k*_1_ (*U* (*t*)), *k*_2_ (*U* (*t*)), and *k*_4_ (*U* (*t*)) with the scaling parameter *s* in Eqs. 1a-e. This scaling increases the cell cycle doubling time to fit data whilst accounting for the time spent in state SD. To achieve logistic growth of *U* (*t*), all rates *k*_1_ (*U* (*t*)) to *k*_4_ (*U* (*t*)) depend on *U* (*t*) and the carrying capacity, *C*_CM_, such that,

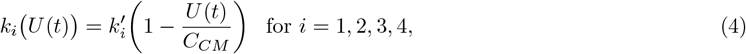

where the values of 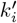 are estimated from in vitro data and are given in Table 2.

#### 2.1.2 Drugs induce DNA damage and inhibit DNA damage repair

The effects of the ATRi (drug 1) and the PARPi (drug 2) are implicitly modelled by inhibition of the rates at which cells progress in the cell cycle model where paths are weighted by factors *p*([drug_2_]) and *q*([drug_1_], [drug_2_]) (Fig. 1**a**). The drug effects are modelled by

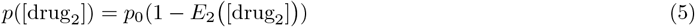

and

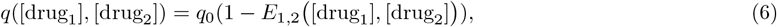

where *p*_0_ and *q*_0_ are the baseline weighting factors that represent the system in the absence of drugs, and [drug_*i*_] denotes the concentration of drug *i*. The functions *E*_1_ and *E*_2_ are calculated using the sigmoid Emax model (Holford, 2017; Hamis et al., 2021b) and are introduced to achieve drug effects that match in vitro data (Section 2.3). We thus set

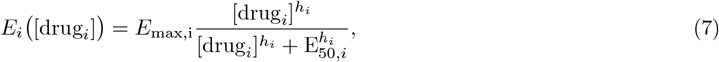

where *i* = 1, 2. *E*_max,i_ is the maximal drug effect, *h*_*i*_ is the Hill coefficient, and E_50,*i*_ is the concentration of the drug that results in half the maximal drug effect. To model the combined effect of the two drugs, we use the Bliss independence synergy model (Koizumi and Iwami, 2014) so that

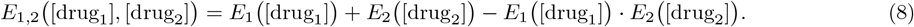

### 2.2 The compartment model is mapped to a baseline agent-based model

As a first step towards studying spatial competition amongst cells (Section 2.4), we here map the compartment model (Fig. 1**a**) onto a baseline 2-D ABM (Fig. 1**c**) following a modelling approach made by e.g., Jenner et al. 2022. In the baseline ABM, cells progress through a drug-dependent cell cycle model (Sections 2.2.1 and 2.2.2), and exist on a 2-D lattice in which each lattice site can either be occupied by a drug-sensitive cancer cell or be empty (Section 2.2.3).

#### 2.2.1 Cells stochastically progress through the cell cycle according to a cell cycle model

Each agent (i.e., cell) in the ABM progresses through the cell cycle via an agent-intrinsic cell cycle model that consists of a set of nodes, that represent cell cycle states, and edges, that represent paths between states (Fig. 1**c**). In the ABM, we include the cell cycle states in the compartment model (Fig. 1**a**) and an additional quiescent state, G0 (Fig. 1**c**). The G0 state is introduced to achieve growth inhibition by spatial limitations. Cells in the ABM are equipped with a cell cycle clock that monitors their progression through the cell cycle and determines when they leave compartments according to allocation times that are estimated from in vitro data (Section 2.3). For each cell, cell_*j*_, in the system, the clock starts when the cell enters the G1 state and terminates when the cell reaches its individual doubling time, *T*_*j*_. The doubling time is chosen from a normal distribution with mean value *τ* and standard deviation *σ*. Here, *τ* and *σ* are estimated from experimental data, where *σ* is chosen to achieve asynchronously dividing cell populations, as observed in the regarded in vitro experiments (Section 2.3). Note that if *σ* = 0, then every dividing cell on the lattice will produce daughter cells at the same time. When a focal cell’s internal cell cycle clock terminates and the cell leaves the G2/M state, it attempts to produce a daughter cell (Section 2.2.3). If there are no available lattice sites on which to place the daughter cell, the cell division fails and the focal cell enters the G0 state. A cell that enters G0 will remain there for the remainder of the simulation. To represent cell cycle arrest, a cell’s internal cell cycle clock is paused when a cell is in the SD state.

#### 2.2.2 Drugs stochastically induce DNA damage and inhibit DNA damage repair

As in the compartment model, the edges in the ABM are assigned weights that are functions of drug doses (Section 2.1.2). The weights denote probabilities with which paths in the cell cycle model are taken. To decide which path is taken at forks, random numbers between zero and one are generated from uniform distributions. If a generated random number at the post-G1 fork is smaller than *p*([drug_2_]) (Eq. 5), cells evade the effect of drug 2 and enter state S. Similarly, if the random number generated at the post-SD fork is smaller than *q*([drug_1_], [drug_2_]) (Eq. 6), cells evade drug effects and enter state S. Drugs are administered at the start of the in silico experiments and are modelled to be constant throughout time and space in the simulations. This mimics an in vitro experiment without drug elimination.

#### 2.2.3 Cells co-exist on a lattice

In the in vitro experiments, cells grow in a monolayer which we abstract to a 2-D space in the ABM. As such, the ABM comprises 100 × 100 lattice sites that can be occupied by one cell or be empty. The baseline ABM has no-flux boundary conditions, which are chosen to represent a physical boundary in an in vitro experiment. At the start of the simulations, *αP*_0_ drug-sensitive cells are seeded on the lattice, where *P*_0_ is the total number of seeded cells on the lattice (chosen to match initial in vitro cell confluency data) and *α* takes the value 0 (in the baseline case), or 0.1 or 0.3 (in the spatial in vitro experiments described in subsection 2.4). The cells are randomly scattered across the lattice in single-cell clusters (Fig. 2**a**) and are initiated in different cycling cell cycle states (i.e., G1, S, SD, G2/M), according to a cell cycle distribution frequency that is estimated from in vitro data (Section 2.3) and correspond to the factors *G*1_0_, *S*_0_, *SD*_0_, *G*2*/M*_0_ in Eqs. 2a-e. When the cell population, *P* (*t*), has reached a critical size *P* *, the drugs are applied to the system. *P* * is stochastic and estimated from in vitro data (Section 2.3). To approximate circular growth of cell clusters, each daughter cell is randomly assigned to be placed in a free lattice site that belongs up to either the *v*_*B*_th order Moore neighbourhood, or the *v*_*B*_th order Von Neumann neighbourhood, of its parental cell (Supplementary Material S1). We estimate *v*_*B*_ from in vitro data (Section 2.3).

**Fig. 2:**
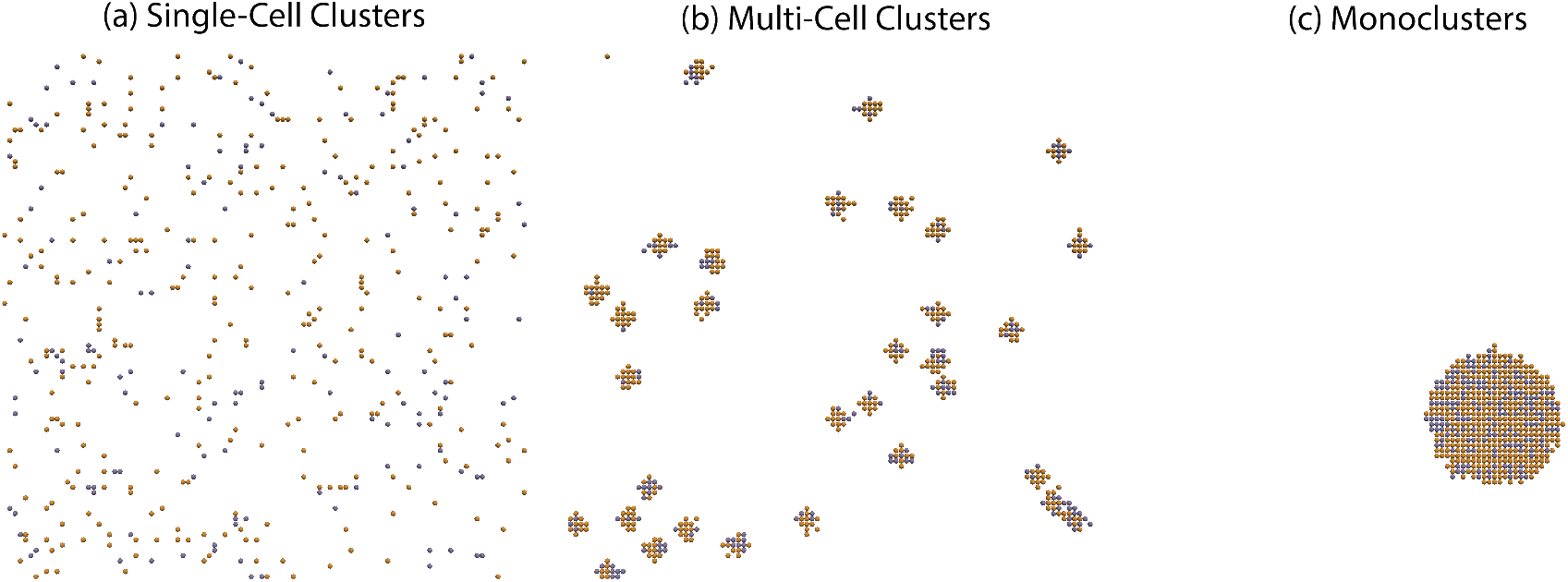
In the cell crowding limited ABM, cells are initiated in (a) single-cell clusters, (b) multi-cell clusters, or (c) monoclusters. In (**a,b,c**), we seed a total number of *P*_0_ cells on the lattice, where 0.7*P*_0_ cells are drug-sensitive (orange) and 0.3*P*_0_ cells are drug-resistant (purple). The example snapshots in (**a,b,c**) are taken when the total cell counts have reached the critical value *P* * which is chosen stochastically and determines when drugs should be applied.

### 2.3 The models are parameterised and evaluated against in vitro data

The compartment model and the ABM parameters are estimated from an in vitro experiment in which FaDu ATM-KO cells were subjected to one or two DNA damage response inhibiting drugs (Lloyd et al., 2020). Cell confluency and cell death data were reported for 0-310 hours, and pulse-chase data were reported for 0-24 hours. To parameterise the compartment model, we first directly read some model parameters (*U* (0), *C*_CM_, *G*1_0_, *S*_0_, *SD*_0_, *G*2*/M*_0_,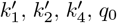) from the in vitro data. Note that the parameter value of *C*_CM_ represents 100% cell confluency in the in vitro experiments. The initial concentration of cells in the non-cycling compartment (*NC*_0_) is assumed to be 0 cells/A, where A denotes the area of the spatial domain in the in vitro experiment. We thereafter estimate the remaining parameters (*p*_0_, 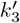, *s, E*_max,1_, *E*_50,1_, *h*_1_, *E*_max,2_, *E*_50,2_, *h*_2_) with a global optimiser in MATLAB that minimises the sum of squares of the residuals between the model output and the in vitro data (Ugray et al., 2007). Following the 80-20 rule of thumb, which recommends using 80% of available data to train a model, and 20% of available data to test a model (Joseph, 2022), we use data from 9 time series experiments to parameterise (or train) the model (Fig 1**b** I-V), and 2 time series experiments to evaluate (or test) the model (Fig 1**b** VI). A data-driven model selection that motivates the choice of the model equations (Eqs. 1a-2e) is included in the Supplementary Material (S3).

After the model evaluation and selection, the compartment model is mapped to the baseline ABM. First, some ABM model parameters (*C*_ABM_, *µ*_*P*_ *, *q*_0_, *p*_0_, *E*_max,1_, *E*_50,1_, *h*_1_, *E*_max,2_, *E*_50,2_, *h*_2_, *τ*, *τ*_*G*1_, *τ*_*S*_, *τ*_*SD*_, and *τ*_*G*2*/M*_) are carried over from the compartment model. Thereafter, some ABM-specific parameters (*σ*_*P*_ * and *σ*) are directly read from the in vitro data. Lastly, the parameter *v*_*B*_ is estimated by matching simulation outputs to in vitro data.

Parameter values related to the initial conditions are listed in Table 1 and all other parameters are listed in Table 2. A thorough description of the model parameter estimation is available in the Supplementary Material (S2). Since both the compartment model (Fig. 1**a**) and the ABM (Fig. 1**c**) are able to satisfactorily predict unseen time series data, we argue that the models are appropriate for our current study which aims at modelling cell population dynamics in response to 0.1-1 *µ*M ceralasertib doses and 0.2-1 *µ*M olaparib doses. While our models do not replicate the bumps observed in the in vitro data (e.g., the dashed purple curve at 150 hours in Fig. 1**b** V), we choose not to over-parameterise the models to fit noisy data and thus make a trade-off between model fit and complexity. We stress that the motivation for moving the compartment model to the ABM is *not* to better capture the in vitro system in Fig. 1. Instead, the ABM is introduced in order to next allow us to simulate theoretical scenarios in which FaDu ATM-KO cells are spatially structured (and thus not well mixed). This will enable us to study the role of space in relevant treatment dynamics. Indeed, the compartment model captures the training data better than the ABM, which suggests that the modelled system is well-mixed. For the chosen split of training and testing data, the baseline ABM does, however, capture the test data better than the compartment model. For a quantitative model comparison, see Supplementary Table S1 with root mean square errors between model outputs and in vitro data.

**Table 1:**
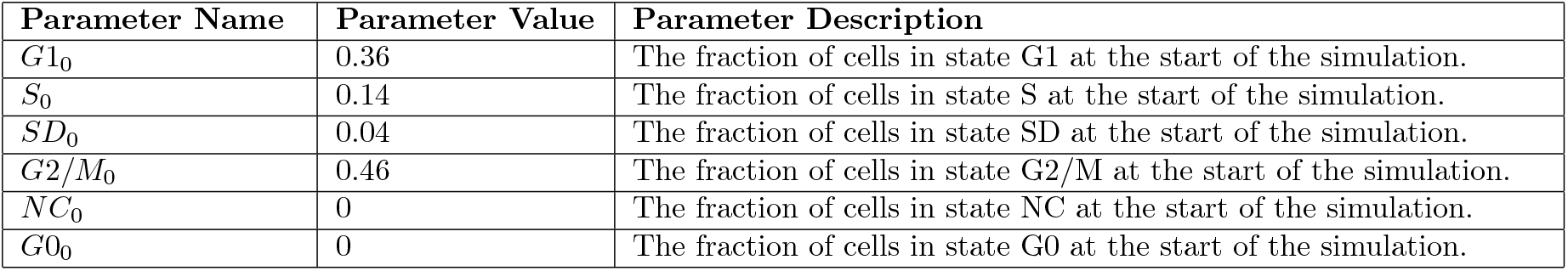
Initial conditions are estimated from in vitro data. The initial fraction of cells in states G1, S, SD, and G2/M are estimated from pulse-chase data. At the start of the simulations, no cells are in the non-cycling states NC and G0. Note that, state G0 is included in the agent-based model but not the compartment model.

**Table 2:**
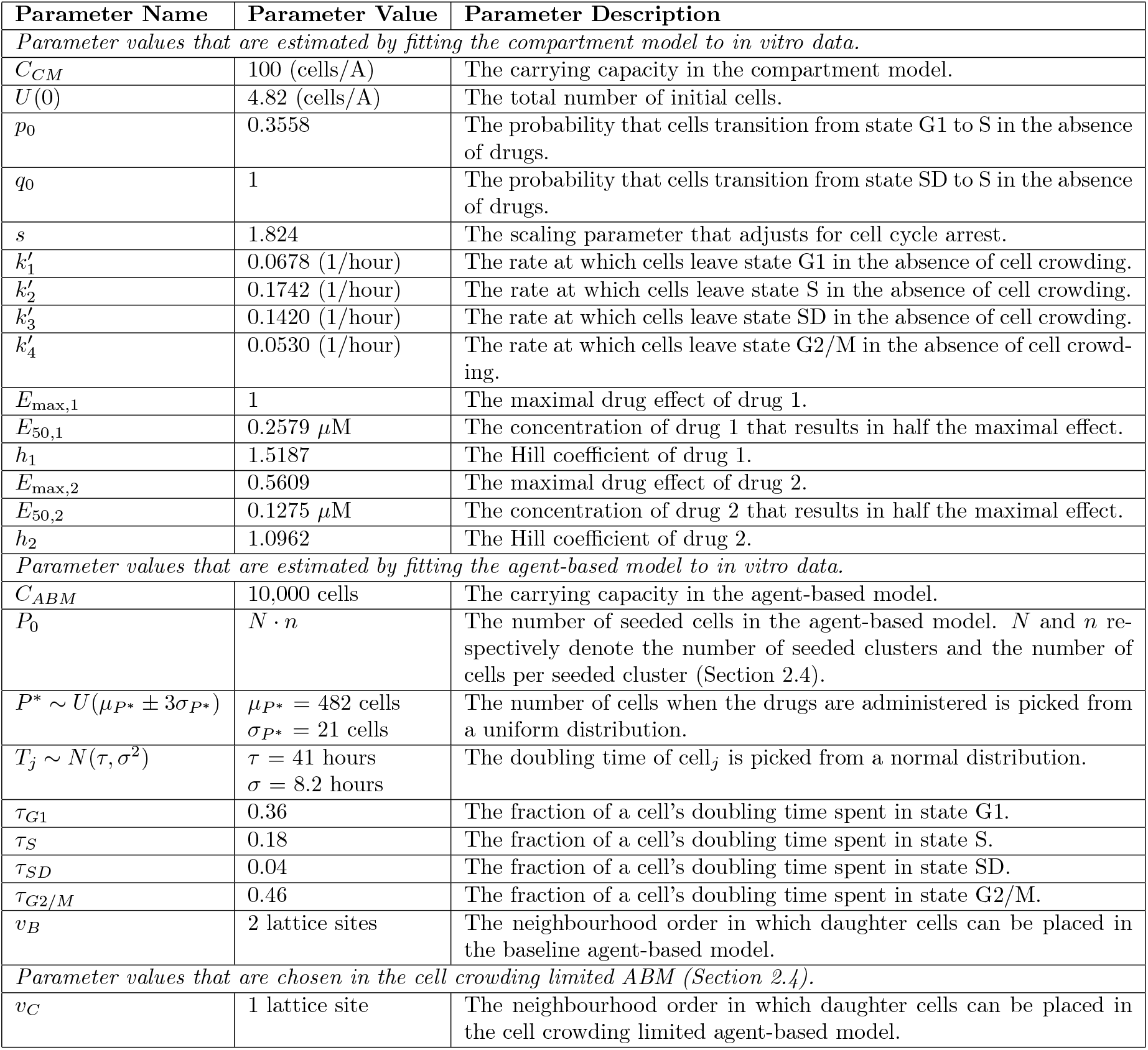
Parameter values are estimated from in vitro data. The table displays the parameter values that are used in the compartment model and the ABMs.

| Parameter Name | Parameter Value | Parameter Description |
| --- | --- | --- |
| *Parameter values that are estimated by fitting the compartment model to in vitro data.* |  |  |
| $C_{CM}$ | 100 (cells/A) | The carrying capacity in the compartment model. |
| $U(0)$ | 4.82 (cells/A) | The total number of initial cells. |
| $p_0$ | 0.3558 | The probability that cells transition from state G1 to S in the absence of drugs. |
| $q_0$ | 1 | The probability that cells transition from state SD to S in the absence of drugs. |
| $s$ | 1.824 | The scaling parameter that adjusts for cell cycle arrest. |
| $k'_1$ | 0.0678 (1/hour) | The rate at which cells leave state G1 in the absence of cell crowding. |
| $k'_2$ | 0.1742 (1/hour) | The rate at which cells leave state S in the absence of cell crowding. |
| $k'_3$ | 0.1420 (1/hour) | The rate at which cells leave state SD in the absence of cell crowding. |
| $k'_4$ | 0.0530 (1/hour) | The rate at which cells leave state G2/M in the absence of cell crowding. |
| $E_{\max,1}$ | 1 | The maximal drug effect of drug 1. |
| $E_{50,1}$ | 0.2579 μM | The concentration of drug 1 that results in half the maximal effect. |
| $h_1$ | 1.5187 | The Hill coefficient of drug 1. |
| $E_{\max,2}$ | 0.5609 | The maximal drug effect of drug 2. |
| $E_{50,2}$ | 0.1275 μM | The concentration of drug 2 that results in half the maximal effect. |
| $h_2$ | 1.0962 | The Hill coefficient of drug 2. |
| *Parameter values that are estimated by fitting the agent-based model to in vitro data.* |  |  |
| $C_{ABM}$ | 10,000 cells | The carrying capacity in the agent-based model. |
| $P_0$ | $N \cdot n$ | The number of seeded cells in the agent-based model. $N$ and $n$ respectively denote the number of seeded clusters and the number of cells per seeded cluster (Section 2.4). |
| $P^* \sim U(\mu_{P^*} \pm 3\sigma_{P^*})$ | $\mu_{P^*} = 482$ cells<br>$\sigma_{P^*} = 21$ cells | The number of cells when the drugs are administered is picked from a uniform distribution. |
| $T_j \sim N(\tau, \sigma^2)$ | $\tau = 41$ hours<br>$\sigma = 8.2$ hours | The doubling time of cell<sub>$j$</sub> is picked from a normal distribution. |
| $\tau_{G1}$ | 0.36 | The fraction of a cell's doubling time spent in state G1. |
| $\tau_S$ | 0.18 | The fraction of a cell's doubling time spent in state S. |
| $\tau_{SD}$ | 0.04 | The fraction of a cell's doubling time spent in state SD. |
| $\tau_{G2/M}$ | 0.46 | The fraction of a cell's doubling time spent in state G2/M. |
| $v_B$ | 2 lattice sites | The neighbourhood order in which daughter cells can be placed in the baseline agent-based model. |
| *Parameter values that are chosen in the cell crowding limited ABM (Section 2.4).* |  |  |
| $v_C$ | 1 lattice site | The neighbourhood order in which daughter cells can be placed in the cell crowding limited agent-based model. |

### 2.4 Competition for space amongst cells is modelled by a cell crowding limited agent-based model

To investigate the role of space in the competition between drug-sensitive and drug-resistant cancer cells, we extend the baseline ABM (Section 2.2) to (1) include both drug-sensitive and drug-resistant cells, and (2) include spatial limitations by cell crowding. In the model, drug-resistant cells can be intrinsically resistant to drug 1 (ATRi), drug 2 (PARPi), or to both drugs 1 and 2. The latter case simulates multi-drug resistance. Furthermore, we incorporate inherited resistance into the model so that drug-resistant parental cells pass on their drug-resistant traits to their daughter cells. In the cell crowding limited ABM, parental cells may only place daughter cells in their immediate neighbourhoods, i.e., in first-order neighbourhoods, such that *v*_*C*_ = 1. To ensure that this spatial competition is an effect of cell crowding and not the boundary, we use periodic boundary conditions in this part of the study.

To investigate how various cell-level and population-level properties impact the competition for space amongst cancer cells, we vary six factors in the initial cell configurations of the ABM, as listed in 1-6 below.

1. The cells are seeded in different spatial configurations on the lattice (Fig. 2).
2. The initial fraction of drug-resistant cells is either 0.1 or 0.3.
3. The drug-resistant cells are resistant to drug 1, drug 2, or both drugs 1 and 2.
4. Cells are treated with either drug 1 monotherapy, drug 2 monotherapy, or drug 1-2 combination therapy.
5. Drugs are given at higher doses ([drug_1_] = 1 *µ*M, [drug_2_] = 1 *µ*M), or lower doses ([drug_1_] = 0.3 *µ*M, [drug_2_] = 0.2 *µ*M).
6. The mean doubling time of drug-resistant cells is varied between *τ* and 2*τ*, where *τ* is the mean doubling time of drug-sensitive cells.

List 1: **To investigate the spatial competition between drug-sensitive and drug-resistant cells in response to drug treatments, we vary items 1-6 in simulation experiments**.

To achieve different spatial configurations (Item 1, List 1), we seed the cells in different-sized clusters. We let *N* denote the total number of initial clusters on the lattice and *n* denote the total number of cells in each seeded cluster. Examples of initial cell configurations with *N* = 419, 32, and 1 cluster(s) are shown in Fig. 2, where the fraction of drug-resistant cells is 0.3.

Before proceeding to the result-generation, we note that simulation stochasticity arises from multiple sources in the ABMs: the initial cell configurations (Sections 2.2.3 and 2.4), the placement of daughter cells (Section 2.2.3), the paths taken in the cell cycle model and the cell doubling times (Section 2.2.1), and drug responses (Section 2.2.2). Moreover, we want to report our result with summary statistics that are averaged over multiple simulations. This begs the question: How many simulations do we need to run in order to report meaningful results that mitigate uncertainty originating from intrinsic model stochasticity? One way to answer this question is through performing a consistency analysis. The process is described in detail by Hamis et al. (2021a) and builds on previous statistical work (Vargha and Delaney, 2000; Alden et al., 2013). From the consistency analysis, we conclude that 100 simulation runs suffice to form the basis for our reported results, as is outlined in the Supplementary Material (S4).

## 3 Results and discussion

We perform a series of in silico experiments that are designed to investigate the dynamics of simulated cell populations in which drug-sensitive and drug-resistant cells co-exist and compete for space. Specifically, we investigate the implications of varying items 1-6 in List 1. We simulate cell population dynamics between 0 and 310 hours to be consistent with the in vitro experiments that we simulate in Fig. 1. Our simulation results are summarised in five result figures (all figures in Section 3) produced with the model parameters listed in Table 2 such that non-cycling cells remain on the lattice and cells can divide in their first order neighbourhoods (*v*_*C*_ = 1). In the Supplementary Material (S6), we reproduce the result figures with the difference that non-cycling cells are removed from the lattice after a time period that is equivalent to their cell cycle length. This cell removal enables cells in state G0 to re-enter the cell cycle, should free space in a neighbouring lattice site become available. Our simulation results show that including such cell removal in the model favours drug-resistant cells which get more spatial resources on which to proliferate. We also reproduce in the result figures in Supplementary Material S7, there with the variation that daughter cells can be placed in up to 2nd-order (*v*_*C*_ = 2) and 3rd-order (*v*_*C*_ = 3) neighbourhoods, as opposed to first-order neighbourhoods only. Our simulation-based conclusions are robust to changes in *v*_*C*_.

### 3.1 Spatial cell structures impact dynamic treatment responses

In the first in silico experiment, we aim to elucidate how the competition between drug-sensitive cells and cells that are resistant to both drugs 1 and 2 is impacted by spatial cell configurations. To do this, we initiate the silico experiments by seeding *P*_0_ cells on the lattice in three different configurations: single-cell clusters, multi-cell clusters, and monoclusters (Item 1, List 1; Fig. 2). For each of these configurations, we vary the initial fractions of drug-resistant cells (Item 2, List 1), drug treatments (Item 4, List 1), and drug doses (Item 5, List 1). The results from these experiments are summarised in Fig. 3, where we plot the fraction of drug-sensitive and drug-resistant cells over the simulation time course, 310 hours. Visual maps of end time cell populations for all investigated initial configurations and dose-combinations are available in the Supplementary Material (S5).

**Fig. 3:**
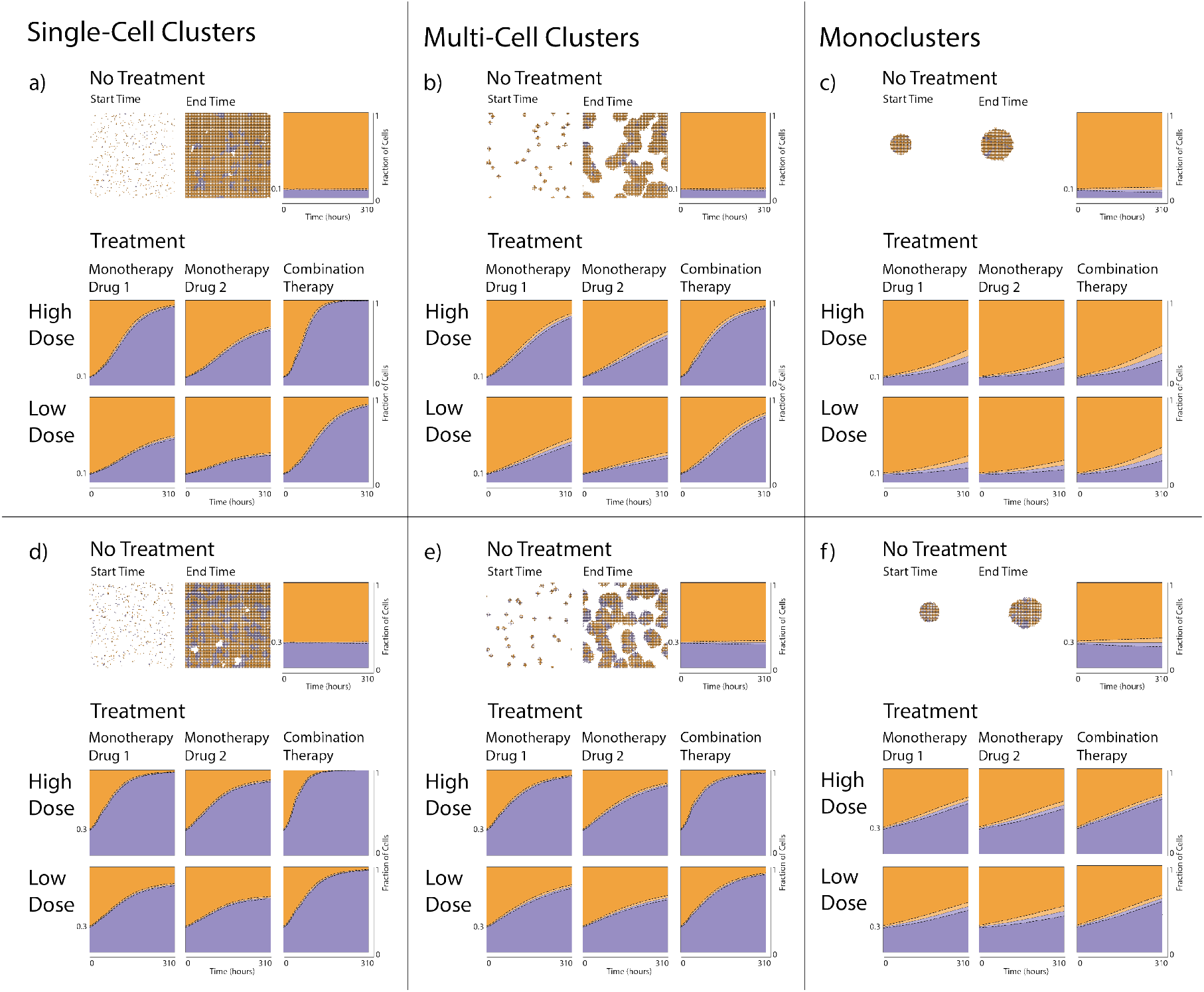
Cells that are resistant to both an ATRi (drug 1) and a PARPi (drug 2) compete for spatial resources with drug-sensitive cells. A total number of *P*_0_ cells are randomly seeded on the lattice in single-cell clusters (**a,d**), multi-cell clusters (**b,e**), or monoclusters (**c,f**), with either 0.1*P*_0_ (**a,b,c**) or 0.3*P*_0_ (**d,e,f**) drug-resistant cells. In each panel **a-f**, results are shown from simulations with no drugs (top row), drug 1 monotherapy (left column in each treatment panel), drug 2 monotherapy (middle column in each treatment panel), and combination therapy (right column in each treatment panel) for high (middle row) and low (bottom row) doses. Examples of initial and final simulation snapshots are shown in each panel. Drug-sensitive and drug-resistant cells are, respectively, orange and purple. The panels also include plots of the dynamic mean fraction of drug-sensitive (orange) and drug-resistant (purple) cells, and standard deviations (dashed lines) from 100 simulation runs.

By comparing the panels in Fig. 3, we can see how the different seeded cell populations lead to different population dynamics. From the results in Fig. 3 (top rows of **a-f**), we see that, in the absence of drugs, the fractions of drug-sensitive and drug-resistant cells do not notably vary over the simulation time course for any seeded cell configuration. This is because there are no drugs to target the drug-sensitive or drug-resistant cells, which means both populations grow at the same rate. However, when drugs are applied in the in silico experiments, the fraction of the drug-resistant populations increase over time due to the drugs targeting the drug-sensitive populations only. This result is particularly significant when the cells are seeded in smaller-sized clusters. For instance, when cells are seeded in single-cell clusters and given high-dose combination treatments, the fraction of drug-resistant cells increases to the extent of dominating the population by the end of the simulations (Fig. 3**a,d**). The partial result that the initial cell configurations impact the dynamic composition of drug-sensitive to drug-resistant cells can be explained by the fact that cells can only proliferate if they have space to do so, i.e., are on the periphery of clusters. As such, cell crowding impedes the growth of both cell populations, whereas drugs impede the growth of drug-sensitive cell populations only. By decreasing the seeded cell cluster sizes, the space for undamaged and dividing cells to proliferate increases. This effect is exasperated when: (i) the initial fraction of drug-resistant cells is high at the start of the simulation, as can be observed by comparing panels (**a,b,c**) to (**d,e,f**) in Fig. 3, (ii) drugs are given at high doses, as can be observed by comparing the middle to bottom rows in panels **a-f** in Fig. 3, (iii) drugs are administered in combination, as can be observed by comparing the columns in treatment panels **a-f** in Fig. 3.

In Figs. 4 and 5, we have repeated the in silico experiments that are summarised in Fig. 3, for cases where the drug-resistant cells are resistant to drugs 1 or 2 only. The results show that ATRi-treatments on populations with ATRi resistant cells yield higher drug-resistant fractions than PARPi-treatment on populations with PARPi resistant cells. This difference is exasperated by small cluster sizes and follows from the model calibration to the experimental data which shows that ATRi monotherapies are more effective at inhibiting the repair of DNA-damaged cells than PARPi monotherapies. Rather than being a comparatively effective monotherapy, the PARPi makes the cells more susceptible to the ATRi in our simulations. This follows from the shape of the biologically-informed pathway model (Fig. 1**c**) in which the PARPi pushes cells to the SD state, making them susceptible to the ATRi.

**Fig. 4:**
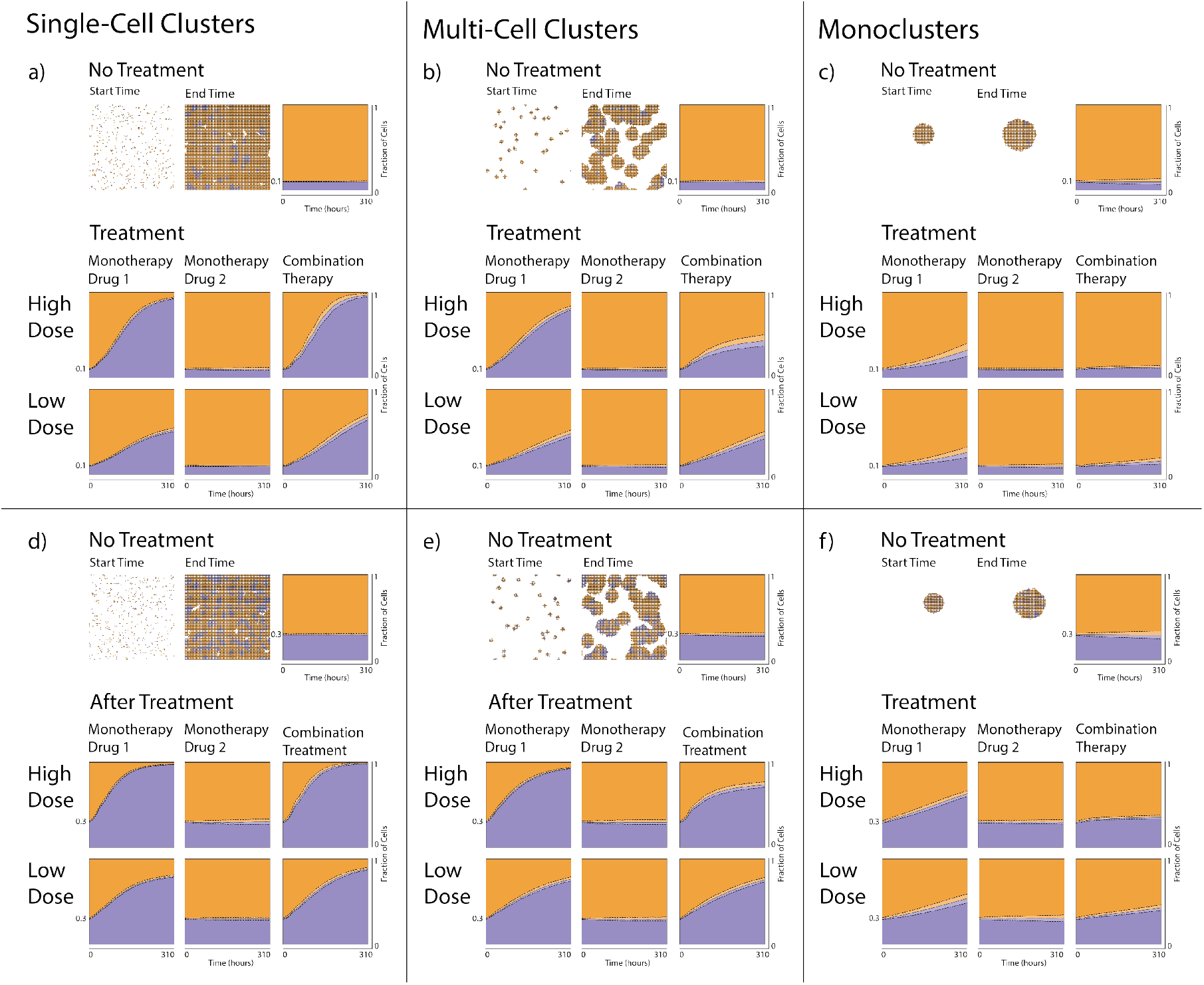
Cells that are resistant to an ATRi (drug 1) compete for spatial resources with drug-sensitive cells. A total number of *P*_0_ cells are randomly seeded on the lattice in single-cell clusters (**a,d**), multi-cell clusters (**b,e**), or monoclusters (**c,f**), with either 0.1*P*_0_ (**a,b,c**) or 0.3*P*_0_ (**d,e,f**) drug-resistant cells. In each panel **a-f**, results are shown from simulations with no drugs (top row), drug 1 monotherapy (left column in each treatment panel), drug 2 monotherapy (middle column in each treatment panel), and combination therapy (right column in each treatment panel) for high (middle row) and low (bottom row) doses. Examples of initial and final simulation snapshots are shown in each panel. Drug-sensitive and drug-resistant cells are, respectively, orange and purple. The panels also include plots of the dynamic mean fraction of drug-sensitive (orange) and drug-resistant (purple) cells, and standard deviations (dashed lines) from 100 simulation runs.

**Fig. 5:**
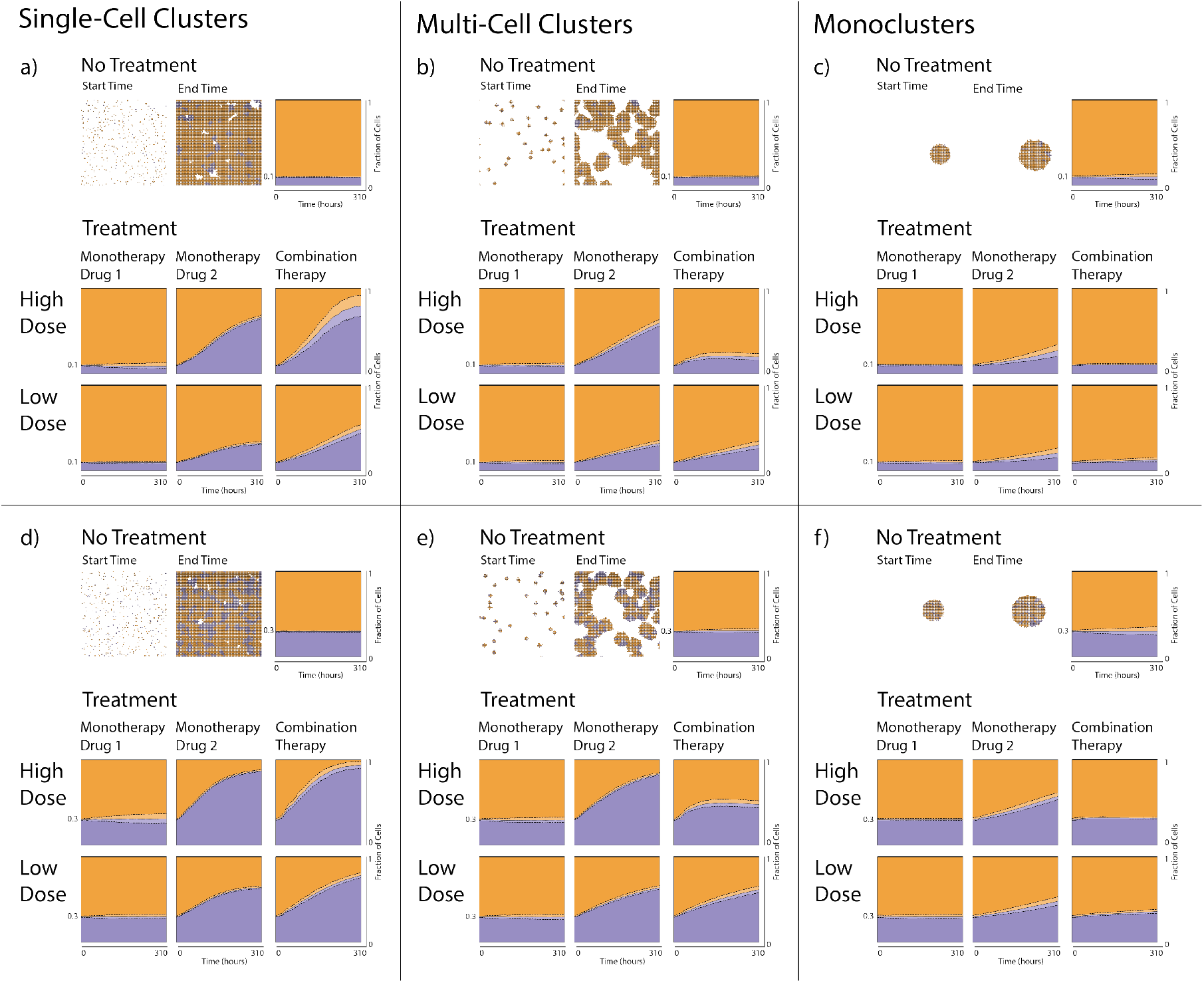
Cells that are resistant to a PARPi (drug 2) compete for spatial resources with drug-sensitive cells. A total number of *P*_0_ cells are randomly seeded on the lattice in single-cell clusters (**a,d**), multi-cell clusters (**b,e**), or monoclusters (**c,f**), with either 0.1*P*_0_ (**a,b,c**) or 0.3*P*_0_ (**d,e,f**) drug-resistant cells. In each panel **a-f**, results are shown from simulations with no drugs (top row), drug 1 monotherapy (left column in each treatment panel), drug 2 monotherapy (middle column in each treatment panel), and combination therapy (right column in each treatment panel) for high (middle row) and low (bottom row) doses. Examples of initial and final simulation snapshots are shown in each panel. Drug-sensitive and drug-resistant cells are, respectively, orange and purple. The panels also include plots of the dynamic mean fraction of drug-sensitive (orange) and drug-resistant (purple) cells, and standard deviations (dashed lines) from 100 simulation runs.

Interestingly, when cells are resistant to only one drug and are seeded in multi-cell clusters or monoclusters, low-dose combination treatments lead to higher fractions of drug-resistant cells compared to high-dose combination treatments at the end of the simulations (Figs. 4 and 5 **b,c,e,f**). This is because even though one of the drugs suppresses both populations, the other drug suppresses the drug-sensitive population only. Therefore, compared to the high-dose combination treatment, the low-dose combination treatment marginally favours the drug-resistant populations in Figs. 4 and 5 **b,c,e,f**. This statement is based on comparing end time fractions of drug-resistant cells only, but does not hold for all simulation time points as is depicted more clearly in Fig. S23 in the Supplementary Material (S8).

In summary, our simulation results demonstrate that there is a complex interplay between in silico cell competition and seeded cell configurations, initial fraction of seeded drug-resistant cells, mode of drug resistance, drug treatments, and drug doses (Items 1-5, List 1).

### 3.2 Spatial cell structures impact the feasibility of treatment objectives

To study how the spatial competition between cells impact not only drug-sensitive and drug-resistant subpopulation sizes, but also total cell counts, we here expand on the experiments described in Section 3.1. To do this, we vary two in silico inputs: the initial spatial cell configurations (Item 1, List 1) and drug doses (Item 5, List 1). In the experiments, we seed *P*_0_ cells on the lattice in different number of clusters, where 0.3*P*_0_ cells are resistant to both drugs 1 and 2. Importantly, the number of seeded cells is the same for all in silico experiments, but the number of clusters in which the cells are seeded ranges between single-cell clusters to monoclusters. We also vary the combination drug dose between 0 and 1 *µ*M. Simulation results are summarised in heatmaps in Fig. 6, where the first heatmap (**a**) shows the total cell count, and the second heatmap (**b**) shows the fraction of drug-resistant cells at the end of the simulation for different drug doses (vertical axes) and number of seeded clusters (horizontal axes).

**Fig. 6:**
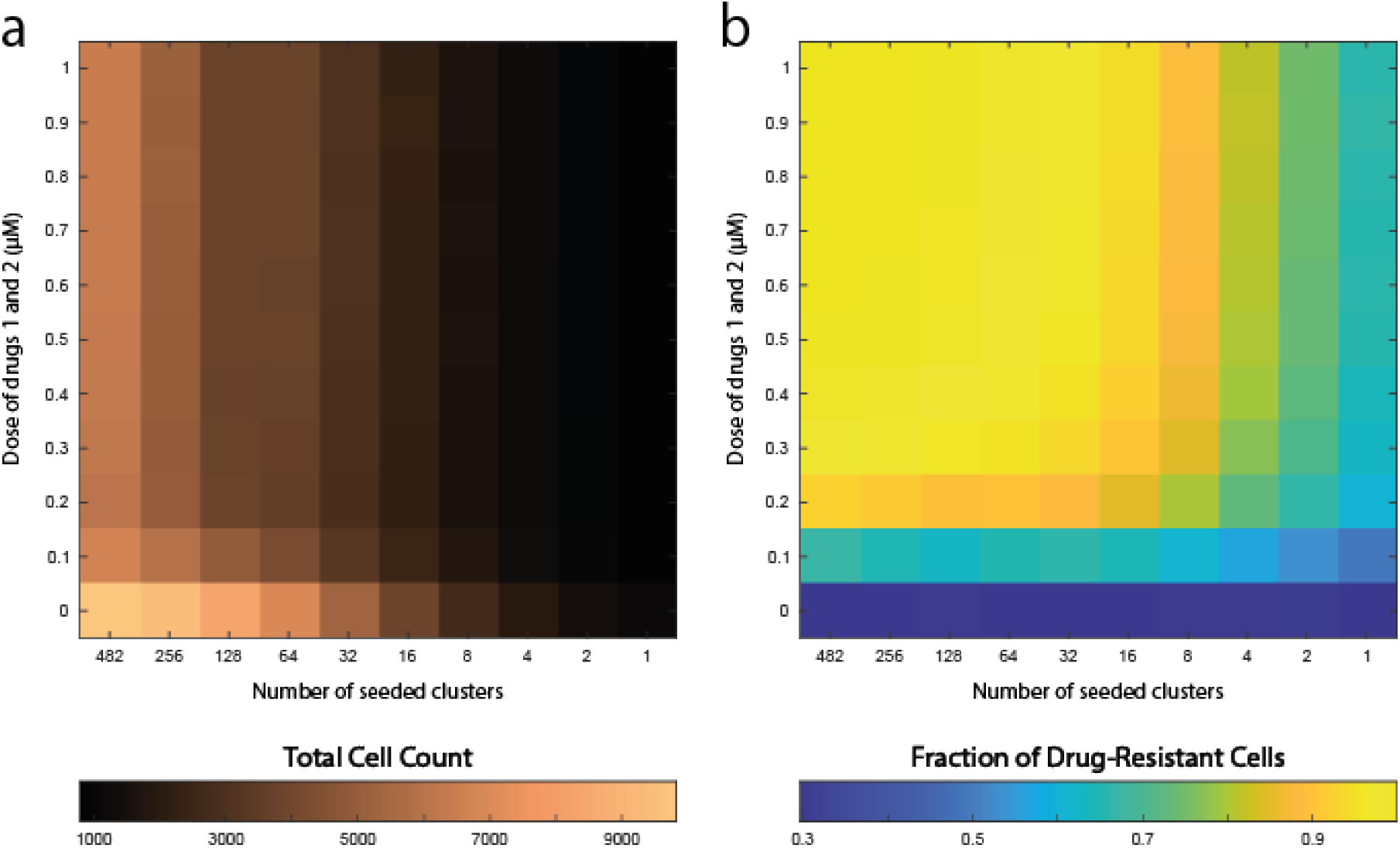
Drug doses and spatial cell configurations impact total cell counts and the composition between drug-sensitive and drug-resistant cells. The heatmaps show results from in silico experiments in which drug-sensitive and drug-resistant cells coexist. At the start of the experiments, a total number of *P*_0_ cells are seeded on the lattice, where 0.3*P*_0_ cells are resistant to both drugs 1 and 2. Two inputs are varied in the simulations: (1) the number of clusters in which cells are seeded (indicated by the horizontal heatmap axes), and (2) the combination treatment drug doses (indicated by the vertical heatmap axes). The results show (**a**) the total cell count and (**b**) the fraction of drug-resistant cells at the end of the simulations. Each heatmap bin shows the mean value of 100 simulation runs.

Our simulation results show that the end time cell counts increase with the number of clusters and decrease with increasing drug doses (Fig. 6**a**). Drug-resistant population fractions also increase with the number of clusters but increase with increasing drug doses (Fig. 6**b**). From an in silico treatment perspective, this means that the treatment objective should influence the applied drug doses. Indeed, biologically relevant treatment objectives include (i) suppressing the total cell count, and (ii) suppressing the drug-resistant fraction, and might be best achieved by different doses. We exemplify this argument by considering a scenario in which the initial cell configuration (i.e., the number of seeded clusters) is fixed, but the applied drug dose is human-controllable. In this example, we start with 64 clusters and our treatment objective is to keep the end time cell count below 5000 cells. We can use Fig. 6**a** to see that the drug dose needs to be in the range 0.1-1 *µ*M. From Fig. 6**b**, we also see that dose 1 *µ*M yields an end time drug-resistant fraction of 0.99, whereas dose 0.1 *µ*M yields an end time drug-resistant fraction of 0.66. Hence, one could argue that the latter drug dose is more favourable than the former drug dose from a treatment perspective.

Overall, our results suggest that different treatment objectives (e.g., suppressing cell counts, or suppressing drug-resistant cell populations) should be leveraged when designing treatment strategies (here, drug doses) since the objectives might be best achieved with different approaches.

### 3.3 Spatial cell structures impact doubling time associated treatment responses

In the previous in silico experiments (Sections 3.1 and 3.2), the doubling times of drug-sensitive and drug-resistant cells are picked from the same normal distribution with mean doubling time, *τ*, and standard deviation *σ*. Now we investigate how population dynamics are affected when the mean doubling time *τ*_*R*_ of the drug-resistant cells are increased. As done in previous experiments, we seed *P*_0_ cells on the lattice in single-cell clusters, multi-cell clusters, or monoclusters (Item 1, List 1) where 0.3*P*_0_ cells are resistant to both drugs 1 and 2. We also, simultaneously, vary the drug 1 and 2 doses (Item 5, List 1; Fig. 7 vertical axes), and the mean doubling time of drug-resistant cells (Item 6, List 1; Fig. 7 horizontal axes). The results are summarised in heatmaps in Fig. 7, where we measure both the total cell count (top panel) and the fraction of drug-resistant cells (bottom panel) at the end of the simulations for different drug doses and drug-resistant doubling times for each seeded cell configurations.

**Fig. 7:**
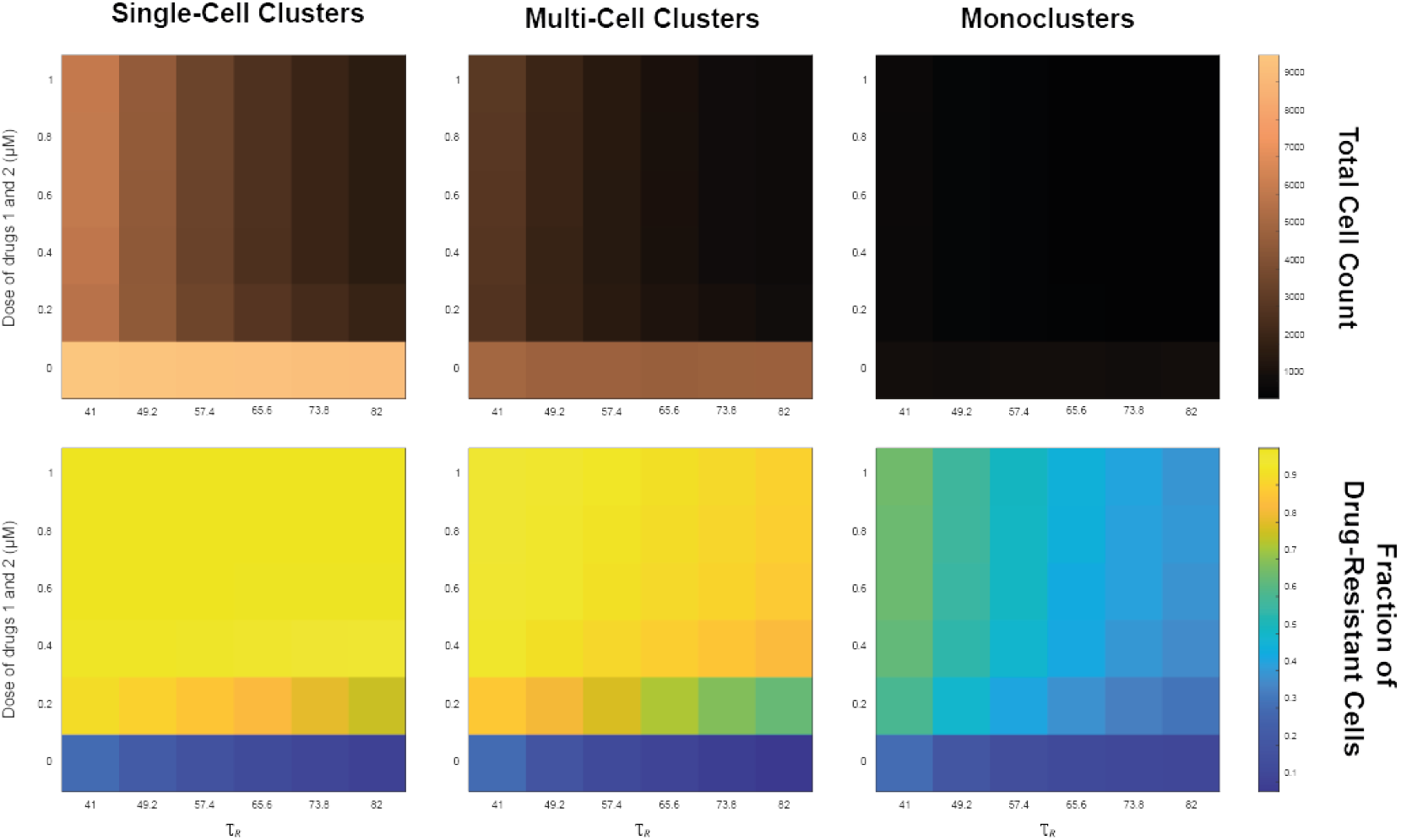
Spatial cell configurations, drug doses, and cell doubling times impact total cell counts and the composition between drug-sensitive and drug-resistant cells. At the start of the experiments, a total number of *P*_0_ cells are seeded on the lattice, where 0.3*P*_0_ cells are resistant to both drugs 1 and 2. Three inputs are varied in the simulations: (1) the seeded cluster configurations (left, middle, right panel), (2) the combination treatment drug doses (indicated by the vertical heatmap axes), and (3) the mean doubling time of the drug-resistant cells (indicated by the horizontal heatmap axes). The heatmaps show the total cell count (top panel) and the fraction of drug-resistant cells (bottom panel) at the end of the simulation. Each heatmap bin shows the mean value of 100 simulation runs.

As expected, our simulation results show that both cell counts and drug-resistant cell fractions decrease with increasing values of *τ*_*R*_, due to fewer cell-divisions of drug-resistant cells. What is interesting to note, however, is that cell populations that are seeded in monoclusters are less sensitive to changes in *τ*_*R*_, than populations that are seeded in single-cell and multi-cell clusters (Fig. 7). This can be observed by comparing the spread in cell counts in the top heatmaps in Fig. 7, and can be explained by the spatial limitations that follow from increased cluster sizes. On the other hand, the relationship between (1) cell configurations, (2) drug doses, and (3) doubling times is more complicated to delineate. For instance, for the no drug treatment, the end time drug-resistant fraction varied between 0.3003-0.1029, 0.3001-0.0775, 0.3034-0.1274 when *τ*_*R*_ is increased from 41 to 82 hours for single-cell, multicell, and monoclusters, respectively. However, for the 1 uM drug dose, these fractions respectively vary between 0.9990-0.9979, 0.9691-0.9014, and 0.6661-0.3950 (Fig. 7, bottom rows). Together, these results indicate that spatial factors may significantly impact simulated cell populations’ sensitivities to both cell-extrinsic factors (e.g., drug doses) and cell-intrinsic factors (e.g., variations in doubling times).

## 4 Conclusion

A growing body of mathematical modelling work highlight that spatio-temporal factors may impact cell-to-cell interactions and the ensuing dynamics in cancer cell populations. Recent contributions to this body of work includes studies on interactions between drug-sensitive and drug-resistant cells (Strobl et al., 2022), growth-factor producing and non-producing cells (Hamis et al., 2023), and cancer cells and immune response cells (Retzlaff et al., 2023; Ruiz-Martinez et al., 2022; van Genderen et al., 2024). In line with these studies we, here, use an on-lattice agent-based modelling approach to simulate spatially structured cell populations comprising drug-sensitive and drug-resistant cells that compete for space on the lattice. The cell-to-cell competition for space is implemented by a volume exclusion modelling rule which ensures that parental cells only divide if free lattice sites are available in their immediate neighborhood. This rule is simple by design, in order to directly probe the role that space plays in the interplay between drug-sensitive and drug-resistant cells in silico. The simulated cell populations are subjected to, up to two, drugs and our results demonstrate that the treatment responses are simultaneously affected by: (i) the initial spatial cell configurations, (ii) the initial fraction of drug-resistant cells, (iii) the drugs to which cells express resistance, (iv) drug combinations, (v) drug doses, and (vi) the doubling time of drug-resistant cells compared to the doubling time of drug-sensitive cells. Adding to the understanding gained from previous ABMs that have been used to study interactions between drug-sensitive and drug-resistant cancer cells (Wang et al., 2015; Strobl et al., 2022; Bacevic et al., 2017), our study contributes simulation-based evidence that factors (i-vi) impact in silico treatment responses both independently (through variation of one factor) and collectively (through simultaneous variation of multiple factors). We iterate that our in silico simulations represent in vitro scenarios, but that our model could be extended to study spatial implications of cell-to-cell interactions in vivo. Such a model extension could include a move from 2-D to 3-D space, the inclusion of healthy cells and other cells of the tumour microenvironment, and plasticity in how individual cells express drug resistance.

In summary, our results demonstrate that spatial factors affect not only cell-to-cell interactions, but also the impact that the cell-to-cell interactions have on the macroscopic population dynamics. From this, we suggest that the role that space plays in cell-to-cell competition should be further investigated and quantified in experimental settings, for instance through analysis of time-lapse microscopy data of cells in varying seeding densities and subclone compositions. We argue that it would be especially meaningful to develop studies that determine when spatial factors impact cell-to-cell interactions and the resulting population dynamics, and when they do not. These studies would be best implemented by multidisciplinary research teams that integrate experiments with mathematical modelling.

## Acknowledgements

KP was supported by EPSRC DTP via Swansea University [Grant EP/T517987/1]. SH was funded by Wenner-Gren Stiftelserna/the Wenner-Gren Foundations (WGF2022-0044), the Tampere Institute for Advanced Study (2021-2023), and the Kjell och Märta Beijer Foundation. We thank the AstraZeneca research and development team for discussions.

## Authors’ contributions

KP customised problem-specific agent-based model (ABM) codes and implemented the ABM simulations, consistency analysis (CA), and parameter estimation. SH provided ABM, CA, and global parameter estimation codes. KP and SH drafted the manuscript. All authors revised the manuscript, have approved of the manuscript, and have agreed to be accountable for all aspects of the work presented in the manuscript.

## Code availability

The code is available on GitHub at https://github.com/KiraPugh/DDR_inhibitor_resistance and extends the code base from https://github.com/SJHamis/DDRinhibitors. Detailed instructions on how to run and modify the codes are provided in the README file in the formerly mentioned repository.

## References

R. Aggarwal. Ceralasertib (AZD6738) alone and in combination with olaparib or durvalumab in patients with solid tumors. Identifier NCT03682289, 2019. URL https://clinicaltrials.gov/study/NCT03682289?cond=ceralasertib%20and%20olaparib&rank=2#study-plan.

J. C. Alamilla-Presuel, A. M. Burgos-Molina, A. González-Vidal, F. Sendra-Portero, and M.J. Ruiz-Gómez. Factors and molecular mechanisms of radiation resistance in cancer cells. Int J Radiat Biol, 98(8):1301–1315, 2022. doi:10.1080/09553002.2022.2047825.

K. Alden, M. Read, J. Timmis, P. S. Andrews, H. Veiga-Fernandes, and M. Coles. Spartan: A comprehensive tool for understanding uncertainty in simulations of biological systems. PLoS Comput Biol, 9(2):e1002916, 2013. doi:10.1371/journal.pcbi.1002916.

E. Armingol, A. Officer, O. Harismendy, and N. E. Lewis. Deciphering cell-cell interactions and communication from gene expression. Nat Rev Genet, 22(2):71–88, 2021. doi:10.1038/s41576-020-00292-x.

AstraZeneca. A study of ceralasertib monotherapy and ceralasertib plus durvalumab in patients with melanoma and resistance to PD-(L)1 inhibition (monette). Identifier NCT05061134, 2022a. URL https://clinicaltrials.gov/study/NCT05061134?cond=NCT05061134&rank=1.

AstraZeneca. An open-label phase 1 study of ceralasertib in Japanese patients with advanced solid malignancies. Identifier NCT05469919, 2022b. URL https://clinicaltrials.gov/study/NCT05469919?cond=NCT05469919&rank=1.

K. Bacevic, R. Noble, A. Soffar, O. W. Ammar, B. Boszonyik, S. Prieto, C. Vincent, M. E. Hochberg, L. Krasinska, and D. Fisher. Spatial competition constrains resistance to targeted cancer therapy. Nat Commun, 8(1):1995, 2017. doi:10.1038/s41467-017-01516-1.

J. S. Baxter, D. Zatreanu, S. J. Pettitt, and C. J. Lord. Resistance to DNA repair inhibitors in cancer. Mol Oncol, 16(21):3811–3827, 2022. doi:10.1002/1878-0261.13224.

S. M. Brady-Kalnay. Molecular mechanisms of cancer cell-cell interactions: Cell-cell adhesion-dependent signaling in the tumor microenvironment. Cell Adh Migr, 6(4):344–345, 2012. doi:10.4161/cam.21489.

M. Cahuzac, B. Péant, A. Mes-Masson, and F. Saad. Development of olaparib-resistance prostate cancer cell lines to identify mechanisms associated with acquired resistance. Cancers (Basel), 14(16):3877, 2022. doi:10.3390/cancers14163877.

A. R. Chaudhuri and A. Nussenzweig. The multifaceted roles of PARP1 in DNA repair and chromatin remodelling. Nat Rev Mol Cell Biol, 18(10):610–621, 2017. doi:10.1038/nrm.2017.53.

S. Checkley, L. MacCallum, J. Yates, P. Jasper, H. Luo, J. Tolsma, and C. Bendtsen. Bridging the gap between in vitro and in vivo: Dose and schedule predictions for the ATR inhibitor AZD6738. Sci Rep, 5:13545, 2015. doi:10.1038/srep13545.

R. Emond, J. I. Griffiths, V. K. Grolmusz, A. Nath, J. Chen, E. F. Medina, R. S. Sousa, T. Synold, F. R. Adler, and A. H. Bild. Cell facilitation promotes growth and survival under drug pressure in breast cancer. Nat Commun, 14(1):3851, 2023. doi:10.1038/s41467-023-39242-6.

T. B. Emran, A. Shahriar, A. R. Mahmud, T. Rahman, M. H. Abir, M. F. Siddiquee, H. Ahmed, N. Rahman, F. Nainu, E. Wahyudin, S. Mitra, K. Dhama, M. M. Habiballah, S. Haque, A. Islam, and M. M. Hassan. Multidrug resistance in cancer: Understanding molecular mechanisms, immunoprevention and therapeutic approaches. Front Oncol, 12:891652, 2022. doi:10.3389/fonc.2022.891652.

Food and Drug Administration. FDA approves olaparib tablets for maintenance treatment in ovarian cancer, 2017. URL https://www.fda.gov/drugs/resources-information-approved-drugs/fda-approves-olaparib-tablets-maintenance-treatment-ovarian-cancer.

Food and Drug Administration. FDA approves olaparib for germline BRCA-mutated metastatic breast cancer, 2018. URL https://www.fda.gov/drugs/resources-information-approved-drugs/fda-approves-olaparib-germline-brca-mutated-metastatic-breast-cancer.

Food and Drug Administration. FDA approves olaparib for gBRCAm metastatic pancreatic adenocarcinoma, 2019. URL https://www.fda.gov/drugs/resources-information-approved-drugs/fda-approves-olaparib-gbram-metastatic-pancreatic-adenocarcinomac.

Food and Drug Administration. FDA approves olaparib with abiraterone and prednisone (or prednisolone) for BRCA-mutated metastatic castration-resistant prostate cancer, 2023. URL https://www.fda.gov/drugs/drug-approvals-and-databases/fda-approves-olaparib-abiraterone-and-prednisone-or-prednisolone-brca-mutated-metastatic-castration.

R. Friedman. Drug resistance in cancer: Molecular evolution and compensatory proliferation. Oncotarget, 7(11):11746–11755, 2016. doi:10.18632/oncotarget.7459.

R. A. Gatenby, Y. Artzy-Randrup, T. Epstein, D. R. Reed, and J. S. Brown. Eradicating metastatic cancer and the eco-evolutionary dynamics of anthropocene extinctions. Cancer Res, 80(3):613–623, 2020. doi:10.1158/0008-5472.CAN-19-1941.

S. O. Halacli, B. Halacli, and K. Altundag. The significance of heat shock proteins in breast cancer therapy. Med Oncol, 30(2):575, 2013. doi:10.1007/s12032-013-0575-y.

S. Hamis, P. Nithiarasu, and G. G. Powathil. What does not kill a tumour may make it stronger: In silico insights into chemotherapeutic drug resistance. J Theor Biol, 454:253–267, 2018. doi:10.1016/j.jtbi.2018.06.014.

S. Hamis, S. Stratiev, and G. G. Powathil. Uncertainty and sensitivity analyses methods for agent-based mathematical models: An introductory review. In The Physics of Cancer, chapter 1, pages 1–37. World Scientific, 2021a. doi:10.1142/97898112234950001.

S. Hamis, J. Yates, M. A. J. Chaplain, and G. G. Powathil. Targeting cellular DNA damage responses in cancer: An in vitro-calibrated agent-based model simulating monolayer and spheroid treatment responses to ATR-inhibiting drugs. Bull Math Biol, 83(10):103, 2021b. doi:10.1007/s11538-021-00935-y.

S. Hamis, P. Somervuo, J.A. Ågren, D. S. Tadele, J. Kesseli, J. G. Scott, M. Nykter, P. Gerlee, D. Finkelshtein, and O. Ovaskainen. Spatial cumulant models enable spatially informed treatment strategies and analysis of local interactions in cancer systems. J Math Biol, 86(5):68, 2023. doi:10.1007/s00285-023-01903-x.

N. Holford. Pharmacodynamic principles and the time course of immediate drug effects. Transl Clin Pharmacol, 25(4):157–161, 2017. doi:10.12793/tcp.2017.25.4.157.

M. V. Hunter, R. Moncada, J. M. Weiss, I. Yanai, and R. M. White. Spatially resolved transcriptomics reveals the architecture of the tumor-microenvironment interface. Nat Commun, 12(1):6278, 2021. doi:10.1038/s41467-021-26614-z.

E. D. Israels and L. G. Israels. The cell cycle. Oncologist, 5(6):510–513, 2000. doi:10.1634/theoncologist.5-6-510.

K. Janeway. Olaparib with ceralasertib in recurrent osteosarcoma. Identifier NCT04417062, 2022. URL https://clinicaltrials.gov/study/NCT04417062?cond=ceralasertib%20and%20olaparib&rank=1.

A. L. Jenner, M. Smalley, D. Goldman, W. F. Goins, C. S. Cobbs, R. B. Puchalski, E. A. Chiocca, S. Lawler, P. Macklin, A. Goldman, and M. Craig. Agent-based computational modeling of glioblastoma predicts that stromal density is central to oncolytic virus efficacy. iScience, 25(6), 2022. doi:10.1016/j.isci.2022.104395.

V. R. Joseph. Optimal ratio for data splitting. Stat Anal Data Min, 15(4):531–538, 2022. doi:10.1002/sam.11583.

E. Karimi, M. W. Yu, S. M. Maritan, L. J. M. Perus, M. Rezanejad, M. Sorin, M. Dankner, P. Fallah, S. Doré, D. Zuo, B. Fiset, D. J. Kloosterman, L. Ramsay, Y. Wei, S. Lam, R. Alsajjan, I. R. Watson, G. R. Urgoiti, M. Park, D. Brandsma, D. L. Senger, J. A. Chan, L. Akkari, K. Petrecca, M. Guiot, P. M. Siegel, D. F. Quail, and L. A. Walsh. Single-cell spatial immune landscapes of primary and metastatic brain tumours. Nature, 614 (7948):555–563, 2023. doi:10.1038/s41586-022-05680-3.

A. Kaznatcheev, J. Peacock, D. Basanta, A. Marusyk, and J. G. Scott. Fibroblasts and alectinib switch the evolutionary games played by non-small cell lung cancer. Nat Ecol Evol, 3(3):450–456, 2019. doi:10.1038/s41559-018-0768-z.

Y. Koizumi and S. Iwami. Mathematical modeling of multi-drugs therapy: A challenge for determining the optimal combinations of antiviral drugs. Theor Biol Med Model, 11:41, 2014. doi:10.1186/1742-4682-11-41.

R. D. Kouyos, C. J. E. Metcalf, R. Birger, E. Y. Klein, P. Abel zur Wiesch, P. Ankomah, N. Arinaminpathy, T. L. Bogich, S. Bonhoeffer, C. Brower, G. Chi-Johnston, T. Cohen, T. Day, B. Greenhouse, S. Huijben, J. Metlay, N. Mideo, L. C. Pollitt, A. F. Read, D. L. Smith, C. Standley, N. Wale, and B. Grenfell. The path of least resistance: Aggressive or moderate treatment? Proc Biol Sci, 281(1794):20140566, 2014. doi:10.1098/rspb.2014.0566.

W. Li, F. Wang, G. Song, Q. Yu, R. Du, and P. Xu. PARP-1: A critical regulator in radioprotection and radiotherapy-mechanisms, challenges, and therapeutic opportunities. Front Pharmacol, 14:1198948, 2023. doi:10.3389/fphar.2023.1198948.

R. L. Lloyd, P. W. G. Wijnhoven, A. Ramos-Montoya, Z. Wilson, G. Illuzzi, K. Falenta, G. N. Jones, N. James, C. D. Chabbert, J. Stott, E. Dean, A. Lau, and L. A. Young. Combined PARP and ATR inhibition potentiates genome instability and cell death in ATM-deficient cancer cells. Oncogene, 39(25):4869–4883, 2020. doi:10.1038/s41388-020-1328-y.

R. Noble, D. Burri, C. Le Sueur, J. Lemant, Y. Viossat, J. N. Kather, and N. Beerenwinkel. Spatial structure governs the mode of tumour evolution. Nat Ecol Evol, 6(2):207–217, 2022. doi:10.1038/s41559-021-01615-9.

A. C. Obenauf, Y. Zou, A. L. Ji, S. Vanharanta, W. Shu, H. Shi, X. Kong, M. C. Bosenberg, T. Wiesner, N. Rosen, R. S. Lo, and J. Massagué. Therapy-induced tumour secretomes promote resistance and tumour progression. Nature, 520(7547):368–372, 2015. doi:10.1038/nature14336.

M.J. O’Connor. Targeting the DNA damage response in cancer. Mol Cell, 60(4):547–560, 2015. doi:10.1016/j.molcel.2015.10.040.

P. C. O’Leary, H. Chen, Y. U. Doruk, T. Williamson, B. Polacco, A. S. McNeal, T. Shenoy, N. Kale, J. Carnevale, E. Stevenson, D. A. Quigley, J. Chou, F. Y. Feng, D. L. Swaney, N. J. Krogan, M. Kim, M. E. Diolaiti, and A. Ashworth. Resistance to ATR inhibitors is mediated by loss of the nonsense-mediated decay factor UPF2. Cancer Res, 82(21):3950–3961, 2022. doi:10.1158/0008-5472.CAN-21-4335.

K. Pugh, M. Davies, and G. Powathil. A mathematical model to investigate the effects of ceralasertib and olaparib in targeting the cellular DNA damage response pathway. J Pharmacol Exp Ther, 387(1):55–65, 2023. doi:10.1124/jpet.122.001558.

J. Retzlaff, X. Lai, C. Berking, and J. Vera. Integration of transcriptomics data into agent-based models of solid tumor metastasis. Comput Struct Biotechnol J, 21:1930–1941, 2023. doi:10.1016/j.csbj.2023.02.014.

A. Ruiz-Martinez, C. Gong, H. Wang, R.J. Sové, H. Mi, H. Kimko, and A. S. Popel. Simulations of tumor growth and response to immunotherapy by coupling a spatial agent-based model with a whole-patient quantitative systems pharmacology model. PLoS Comput Biol, 18(7):e1010254, 2022. doi:10.1371/journal.pcbi.1010254.

S. Saxena and L. Zou. Hallmarks of DNA replication stress. Mol Cell, 82(12):2298–2314, 2022. doi:10.1016/j.molcel.2022.05.004.

M. Sorin, M. Rezanejad, E. Karimi, B. Fiset, L. Desharnais, L. J. M. Perus, S. Milette, M. W. Yu, S. M. Maritan, S. Doré, E. Pichette, W. Enlow, A. Gagné, Y. Wei, M. Orain, V. S. K. Manem, R. Rayes, P. M. Siegel, S. Camilleri-Bröet, P. O. Fiset, P. Desmeules, J. D. Spicer, D. F. Quail, P. Joubert, and L. A. Walsh. Single-cell spatial landscapes of the lung tumour immune microenvironment. Nature, 614(7948):548–554, 2023. doi:10.1038/s41586-022-05672-3.

M. A. R. Strobl, J. Gallaher, J. West, M. Robertson-Tessi, P. K. Maini, and A. R. A. Anderson. Spatial structure impacts adaptive therapy by shaping intra-tumoral competition. Commun Med (Lond), 2:46, 2022. doi:10.1038/s43856-022-00110-x.

A. Tsuboi, S. Ohsawa, D. Umetsu, Y. Sando, E. Kuranaga, T. Igaki, and K. Fujimoto. Competition for space is controlled by apoptosis-induced change of local epithelial topology. Curr Biol, 28(13):2115–2128, 2018. doi:10.1016/j.cub.2018.05.029.

Z. Ugray, L. Lasdon, J. Plummer, F. Glover, J. Kelly, and R. Martí. Scatter search and local NLP solvers: A multistart framework for global optimization. INFORMS J Comput, 19(3):328–340, 2007. doi:10.1287/ijoc.1060.0175.

A. Vallés-Martí, G. Mantini, P. Manoukian, C. Waasdorp, A. F. Sarasqueta, R. R. de Goeij-de Haas, A. A. Henneman, S. R. Piersma, T. V. Pham, J. C. Knol, E. Giovannetti, M. F. Bijlsma, and C.R. Jiménez. Phosphoproteomics guides effective low-dose drug combinations against pancreatic ductal adenocarcinoma. Cell Rep, 42(6):112581, 2023. doi:10.1016/j.celrep.2023.112581.

M. N. G. van Genderen, J. Kneppers, A. Zaalberg, E. M. Bekers, A. M. Bergman, W. Zwart, and F. Eduati. Agent-based modeling of the prostate tumor microenvironment uncovers spatial tumor growth constraints and immunomodulatory properties. NPJ Syst Biol Appl, 10(1):20, 2024. doi:10.1038/s41540-024-00344-6.

A. Vargha and H.D. Delaney. A critique and improvement of the “cl” common language effect size statistics of McGraw and Wong. J Educ Behav Stat, 25(2):101–132, 2000. doi:10.3102/10769986025002101.

J. Wang. A study to evaluate the safety and pharmacokinetics of ceralasertib in combination with durvalumab in Chinese patients with advanced solid tumours. Identifier NCT05514132, 2022. URL https://clinicaltrials.gov/study/NCT05514132?cond=NCT05514132&rank=1.

X. Wang, H. Zhang, and X. Chen. Drug resistance and combating drug resistance in cancer. Cancer Drug Resist, 2(2):141–160, 2019. doi:10.20517/cdr.2019.10.

Z. Wang, J. D. Butner, R. Kerketta, V. Cristini, and T. S. Deisboeck. Simulating cancer growth with multiscale agent-based modeling. Semin Cancer Biol, 30:70–78, 2015. doi:10.1016/j.semcancer.2014.04.001.

H. Zhang, D. G. Delafield, and L. Li. Mass spectrometry imaging: The rise of spatially resolved single-cell omics. Nat Methods, 20(3):327–330, 2023. doi:10.1038/s41592-023-01774-6.

