## Supplementary Material for "Simulations probe the role of space in the interplay between drug-sensitive and drug-resistant cancer cells"

**S1 Daughter cells are placed on the ABM lattice** **2**

**S2 Model parameters are estimated from in vitro data** **2**

**S8 Low versus high combination treatments are zoomed in on** **29**

#### S1 Daughter cells are placed on the ABM lattice

In the ABM, when a parental cell reaches the end of its cell cycle doubling time, it will attempt to divide. If a free lattice site is available in a permitted neighbourhood, the parental cell will produce a daughter cell. Daughter cells can be placed in von Neumann neighbourhoods (Fig. S1**a**) or Moore neighbourhoods (Fig. S1**b**) (Hamis et al., 2021b). To achieve circular-like growth of the cell populations, the choice of neighbourhood (von Neumann or Moore) is probabilistically selected by generating a random number that is either one or two, with equal probability (Fig. S1**c**) (Hamis et al., 2021b). Note that in the baseline ABM, cells can be placed in up to the $v_B$th order neighbourhood (O.N.), and lower-order neighbourhoods are filled with cells first. Neighbourhood order is represented by the numbers in Fig. S1**a** and **b**, which depict up to the 3rd order neighbourhoods. To simulate growth inhibition caused purely by a lack of space, if all lattice sites up to and including the $v_B$th O.N. are occupied by cells, the parental cell will go into the G0 state, where it will remain for the duration of the in silico simulation (Hamis et al., 2021b).

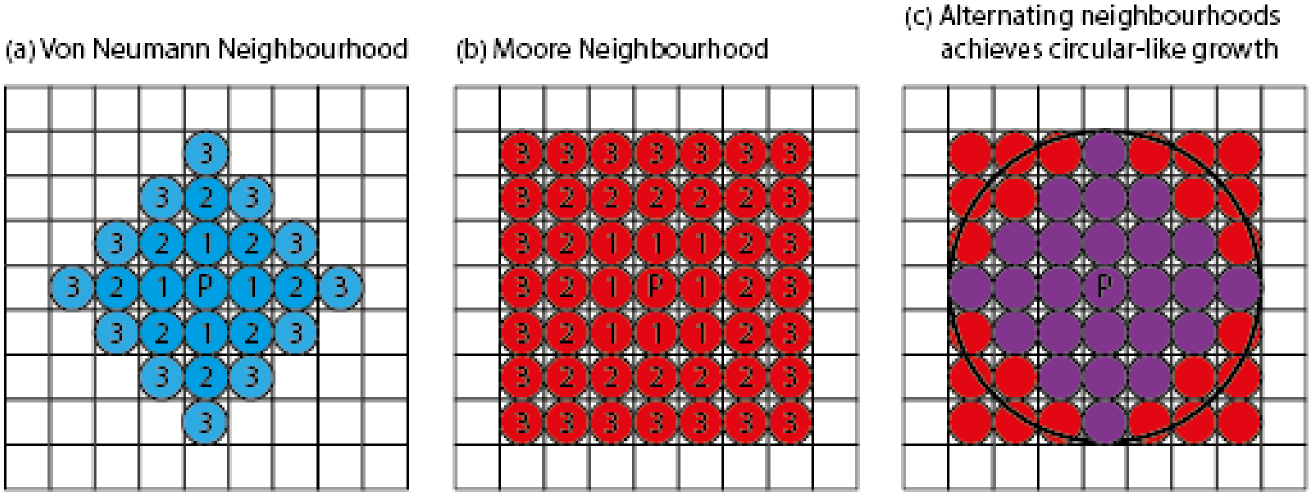

Fig. S1: **Daughter cells are probabilistically placed in von Neumann and Moore neighbourhoods to achieve circular-like growth of the cancer cells.** When a parental cell divides, a daughter cell gets placed on the lattice using either (**a**) the von Neumann neighbourhood or (**b**) the Moore neighbourhood. The numbers in the figure indicate the order of the neighbourhood and P indicates the parental cell. (**c**) Probabilistically placing cells in von Neumann and Moore neighbourhoods achieves circular-like growth.

#### S2 Model parameters are estimated from in vitro data

Table S1 contains the root mean square errors (RMSEs) between mean in vitro cell confluency data and simulated data for both the compartment model and the baseline agent-based model.

##### S2.1 Estimating the parameters in the compartment model

###### Parameters that are estimated directly from the data

The parameters $U(0)$, $C_{\mathrm{CM}}$, $G1_0$, $S_0$, $SD_0$, $G2/M_0$, $k'_1$, $k'_2$, $k'_4$, and $q_0$ are estimated directly from data outputs measured by Lloyd et al. (2020). The initial concentration of cells, $U(0) = 4.82$ (cells/A), and the carrying capacity, $C_{\mathrm{CM}} = 100$ (cells/A), are obtained from initial cell confluency data (Lloyd et al. (2020), Supplementary information Figure 8). These parameter values follow from the mean values of three experimental repeats in the absence of drugs, the initial mean cell confluency is 4.82% and maximum cell confluency is 100%. Note that the cell confluency data also includes standard errors from the experimental repeats (Lloyd et al. (2020), Supplementary information Figure 8).

The initial concentration of cells in each compartment ($G1_0$, $S_0$, $SD_0$, $G2/M_0$) is read from mean values obtained from pulse-chase experiments recorded two hours after the start of the experiments (Lloyd et al., 2020). These experiments measured the fraction of cells in the G1, S, and G2/M phases and cells that are $\gamma$H2AX positive, i.e., DNA damaged (Lloyd et al., 2020). In the mathematical model, cells in the SD state have DNA replication stress and are therefore $\gamma$H2AX positive. The fractions $G1_0$, $S_0$, $SD_0$, and $G2/M_0$ are given in Table 1 in the main manuscript and are multiplied by the initial concentration of cells, $U(0)$ (cells/A), to estimate the initial concentration of cells in compartments G1, S, SD, and G2/M respectively. We assume that all initial cells in the system are cycling (G1, S, or G2/M), or have the potential to become cycling (SD). Therefore, no cells are in the NC state at the beginning of the simulation.

As described in the main manuscript, the rates $k_i\big(U(t)\big)$, depend on $U(t)$ and $C_{\text{CM}}$, such that

$$k_i\big(U(t)\big) = k_i'\bigg(1 - \frac{U(t)}{C_{CM}}\bigg) \quad \text{for } i = 1, 2, 3, 4, \tag{1}$$

where $k_i'$ are the rates of cells leaving the corresponding state when they have the space to grow exponentially. We assume that the fraction of the cell cycle doubling time that cells spend in a state without any spatial restrictions is equal to the initial fraction of cells in a state. Since the values $k_i'$ are measured in units of 1/hour and in the regarded in vitro experiments, the doubling time of FaDu ATM-KO cells was estimated to be $\tau = 41$ hours, we set

$$k_1' = \frac{1}{\tau \cdot G1_0}, \tag{2a}$$

$$k_2' = \frac{1}{\tau \cdot S_0}, \tag{2b}$$

$$k_4' = \frac{1}{\tau \cdot G2/M_0}. \tag{2c}$$

The baseline weighting factor, $q_0$, represents the probability that cells transition from state SD to S in the absence of drugs (Eq. 6 in the main manuscript). In the model, we assume that in the absence of drugs, cells do not obtain irreparable DNA damage. Therefore, all cells repair from the SD state, which means the baseline probability $q_0 = 1$. This is based on observations from in vitro experiments that measured cell death activity using mean fluorescence levels of cytotox. The experiments showed that cell death activity does not increase over time in the absence of drugs.

###### Parameters that are estimated via a global optimiser on MATLAB

The parameters $p_0$, $k_3'$, $s$, $E_{\max,1}$, $E_{50,1}$, $h_1$, $E_{\max,2}$, $E_{50,2}$, and $h_2$ cannot be directly estimated from data and are instead obtained via an optimisation method. Specifically, we use the global optimiser GlobalSearch (Ugray et al., 2007) in MATLAB, that minimises the sum of squares of the residuals between the model output, i.e., the total number of cells in the system, and the in vitro mean cell confluency data which was measured approximately every 3 hours for 310 hours.

##### S2.2 Estimating the parameters in the ABM

###### Parameters that are carried over from the compartment model

The parameters values $C_{\text{ABM}}$, $\mu_{P^*}$, $q_0$, $p_0$, $E_{\max,1}$, $E_{50,1}$, $h_1$, $E_{\max,2}$, $E_{50,2}$, $h_2$, $\tau$, $\tau_{G1}$, $\tau_S$, $\tau_{SD}$, and $\tau_{G2/M}$ are carried over from the compartment model. The ABM-specific carrying capacity is $C_{\text{ABM}} = 10,000$ cells, which is the total number of lattice sites and corresponds to an in vitro cell plate with full cell coverage. The mean number of cells when drugs are administered in the in silico experiments, $\mu_{P^*} = 482$ cells, corresponds to the initial mean cell confluency of 4.82% cells in the in vitro experiments and is obtained by the calculation $\mu_{P^*} = 0.0482 \cdot C_{\text{ABM}}$ cells. Note that to create diversity in initial cell configurations, we ensure that the number of seeded cells $P_0 < \mu_{P^*}$, which allows for cell proliferation before drug administration.

As in the compartment model, we seed cells in different cell cycle states in the ABM, and all seeded cells are either cycling or can become cycling, therefore, none of the cells are seeded in the NC or G0 states. In the ABM, the time spent in state SD does not contribute to a cell's doubling time. Instead, a separate clock is used to track how long cells spend in the SD state. We assume that $G1_0$, $S_0 + SD_0$, $SD_0$, and $G2/M_0$ are the fraction of a cell's doubling time spent in the G1, S, SD, and G2/M state respectively. These fractions are multiplied by $P_0$, to obtain the initial number of cells in each compartment. Note that cells are not necessarily seeded at the beginning of their assigned cell cycle state. Instead, cells are randomly seeded within the range of the cell cycle clock for their assigned cell cycle state, chosen from a uniform distribution.

| Drug Treatment | Compartment Model | Baseline Agent-Based Model (mean) |
|---|---|---|
| DMSO | 6.0381 | 13.2945 |
| 0.2 $\mu$M PARPi | 2.8427 | 12.4894 |
| 1 $\mu$M PARPi | 3.8133 | 8.4309 |
| 0.1 $\mu$M ATRi | 4.2030 | 16.1981 |
| 0.3 $\mu$M ATRi | 3.3602 | 12.3478 |
| 1 $\mu$M ATRi | 1.1291 | 1.8281 |
| 0.1 $\mu$M ATRi + 1 $\mu$M PARPi | 3.2569 | 5.4690 |
| 0.3 $\mu$M ATRi + 0.2 $\mu$M PARPi | 1.7380 | 2.9527 |
| 0.3 $\mu$M ATRi + 1 $\mu$M PARPi | 1.2945 | 1.5630 |
| 1 $\mu$M ATRi + 0.2 $\mu$M PARPi | 0.6523 | 0.4005 |
| 1 $\mu$M ATRi + 1 $\mu$M PARPi | 1.2400 | 0.7739 |
| RSME training data | 3.0751 | 8.2859 |
| RSME test data | 0.9462 | 0.5872 |

Table S1: **A quantitative comparison of the goodness of fit between the in vitro data and the simulated data from the compartment model and the baseline agent-based model.** The table shows the RMSEs between model outputs (compartment model and the baseline agent-based model) and in vitro cell confluency, in units of cell confluency percentages. The table also shows the mean RMSE values for training data (the top 9 dose combinations) and test data (the bottom 2 dose combinations).

**Parameters that are estimated directly from the data**

The parameters $\sigma_{P^*}$ and $\sigma$ are read from data outputs measured by Lloyd et al. (2020). We choose the initial number of cells from a uniform distribution $P^* \sim U(\mu_{P^*} \pm 3\sigma_{P^*})$. The value $\sigma_{P^*} = 21$ cells follows from the standard deviation, 0.21289, of the in vitro initial cell confluency from three experimental repeats. In the ABM, a cell's doubling time is chosen from a normal distribution with mean $\tau = 41$ hours and standard deviation $\sigma = 8.2$ hours. We choose a standard deviation equal to 20% of the mean to achieve an asynchronous population as observed in the in vitro data.

**Parameters that are estimated via optimisation**

The number of neighbourhoods in which daughter cells can be placed in the baseline ABM, $v_B$, is estimated by matching simulation outputs, specifically the percentage of cell coverage on the lattice, to in vitro cell confluency data (Lloyd et al. (2020), Supplementary information Figure 8). The RMSEs between the mean in vitro data and the simulated data from the parameterised models are given in Table S2 for $v_B = 1$, 2, 3, 4, 5, and $\infty$. Since the RMSE test data value stays consistent and the RMSE training data value notably drops between $v_B = 1$ and $v_B = 2$, and we want to use the simplest possible model (i.e., the smallest value of $v_B$), we use $v_B = 2$. Note that the value of $v_B$ is only used to calibrate the baseline ABM, and that parental cells are only allowed to divide in one neighbourhood in the cell crowding limited ABM.

| Drug Treatment | $v_B = 1$ | $v_B = 2$ | $v_B = 3$ | $v_B = 4$ | $v_B = 5$ | $v_B = \infty$ |
|---|---|---|---|---|---|---|
| RSME training data | 12.0068 | 8.2859 | 7.6500 | 7.2334 | 7.2595 | 7.3461 |
| RSME test data | 0.5561 | 0.5872 | 0.5561 | 0.6208 | 0.6118 | 0.6078 |

Table S2: **A quantitative comparison of the goodness of fit between the in vitro data and the simulated data from the cell crowding limited ABM for different values of neighbourhood sizes $v_B$.** The table shows the mean RMSEs between (mean) model outputs and in vitro cell confluency, in units of cell confluency percentages, for training data (the top 9 dose combinations in Table S1) and test data (the bottom 2 dose combinations in Table S1).

### S3 A model selection is performed

We perform a data-driven model selection that motivates the choice of the model equations (Eqs. 1a-1e in the main manuscript). We compare three compartment models: (1) a low complexity model (Model LC, Fig. S3**a**), (2) a

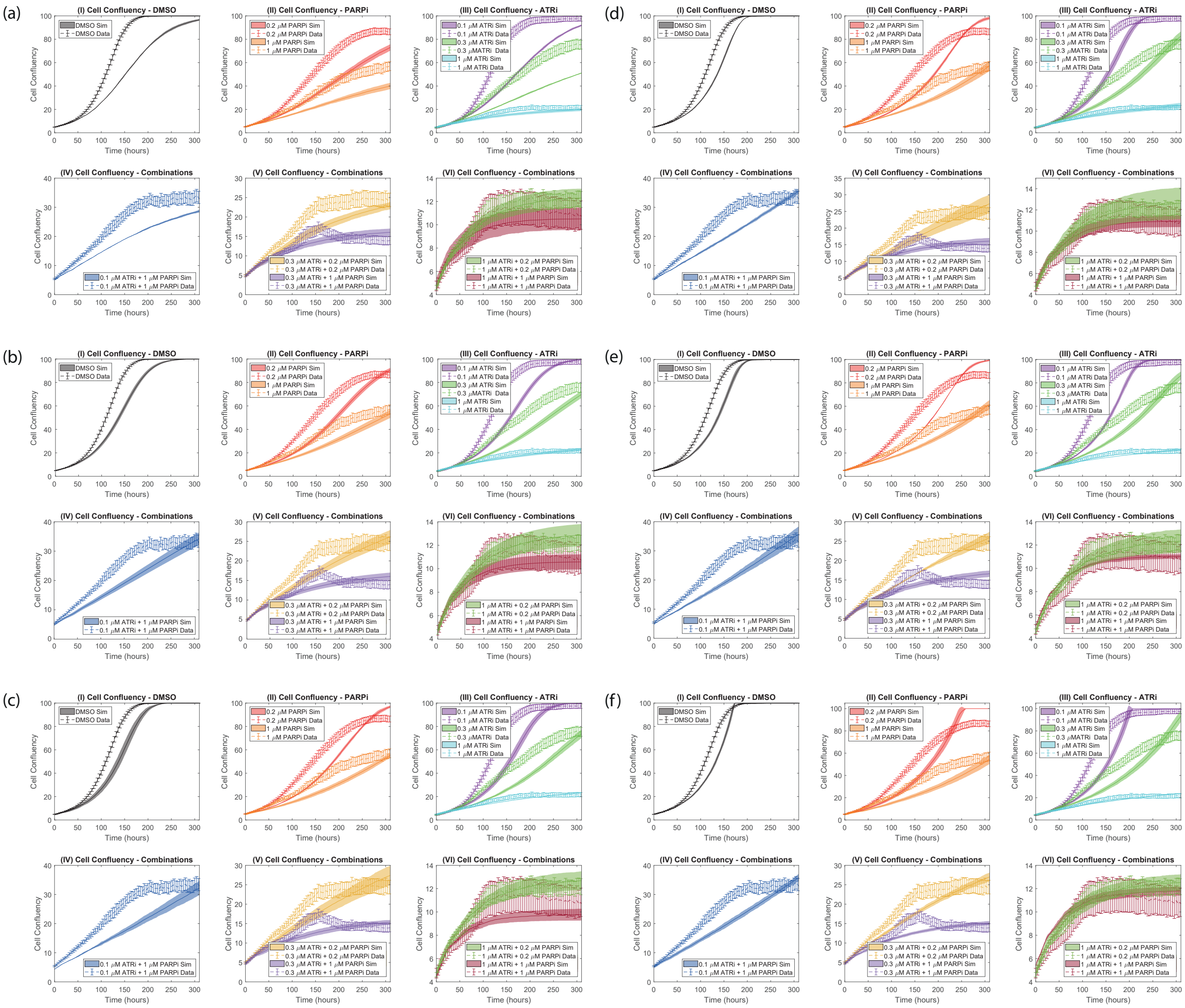

Fig. S2: **The parameter $v_B$, which denotes the neighbourhood order in which daughter cells can be placed in the baseline ABM, is calibrated from experimental data.** The plots show simulated and experimental cell confluency over 310 hours for various dose combinations of the ATRi (drug 1) and PARPi (drug 2). We test multiple values of $v_B$: (**a**) $v_B = 1$, (**b**) $v_B = 2$, (**c**) $v_B = 3$, (**d**) $v_B = 4$, (**e**) $v_B = 5$, and (**f**) $v_B = \infty$. The latter scenario represents the case where cells can divide anywhere within the lattice. The solid lines represent the mean cell confluency from the ABM with standard deviations from 100 simulation runs in opaque bands. The dashed lines represent the mean in vitro data, with standard errors for three experiments indicated with error bars.

medium complexity model (Model MC, Fig. S3**c**), and (3) a high complexity model (Model HC, Fig. S3**e**). All models are described by a system of ODEs, where each dependent variable $[y]$ describes the concentration of cells in compartment $y$.

#### Low Complexity Model

Model LC (Fig. S3**a**) is a modification of Hamis et al. (2021b). The compartments and parameters are the same as in Model MC, which is described in Section 2.1 in the main manuscript. In Model LC, we assume that both drugs act only on the path where cells transition from state SD to S. Therefore, the factor $q$, which represents the fraction of cells transitioning from state SD to S, decreases with increasing drug concentrations (see specifically Eq. 6 in Section 2.1.2 in the main manuscript for more details). To parameterise Model LC, we first directly read some model parameters ($U(0)$, $C_{\text{CM}}$, $G1_0$, $S_0$, $SD_0$, $G2/M_0$, $k_1'$, $k_2'$, $k_4'$, $q_0$) from the in vitro data (Lloyd et al., 2020).

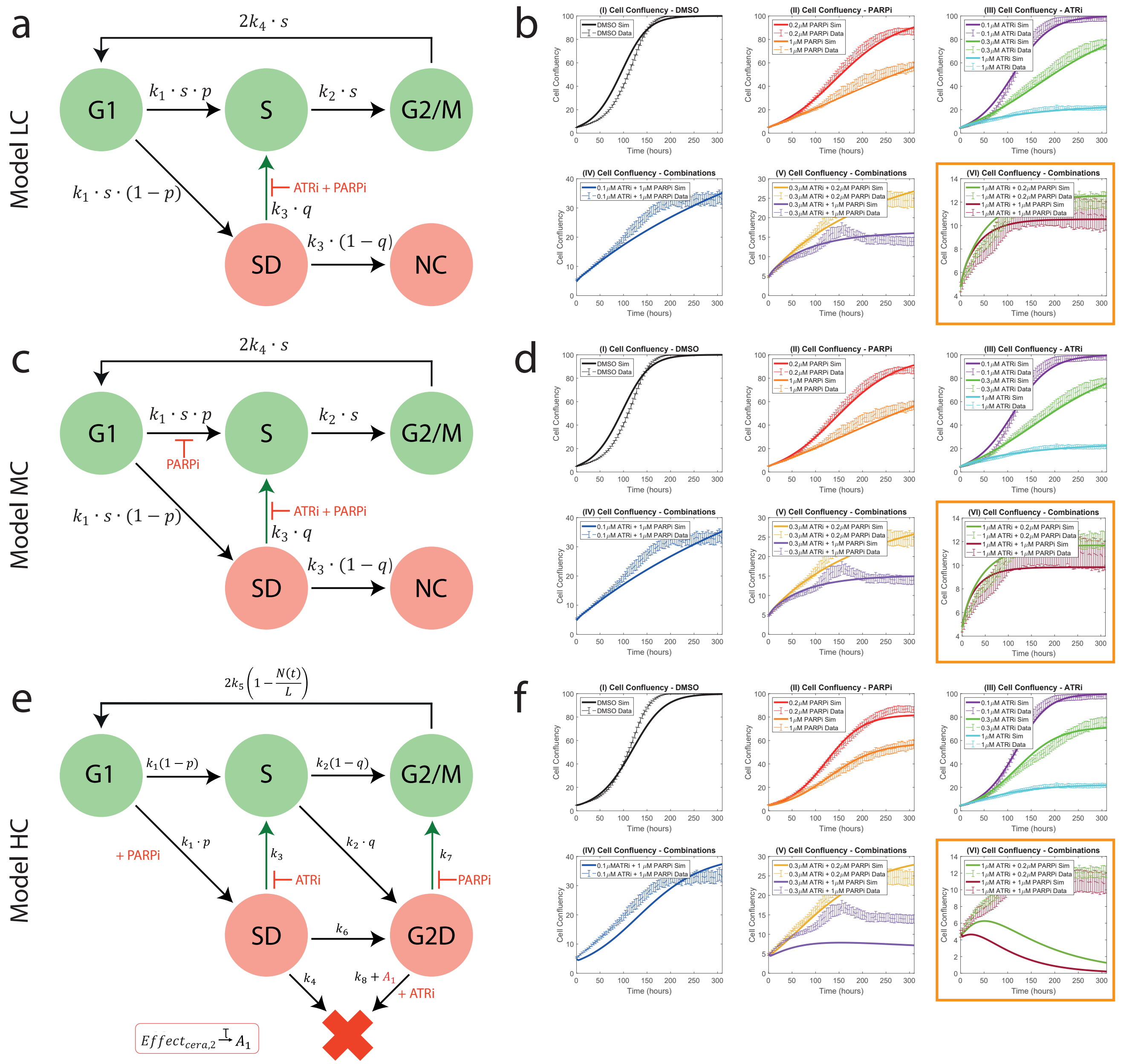

Fig. S3: **Three compartment models are parameterised and evaluated against in vitro data and are visually compared for the best model fit.** The plots show the schematic of the three compartment models: (**a**) Model LC, (**c**) Model MC, and (**e**) Model HC. The time and drug dependencies of the parameters in (**a**,**c**,**e**) have been omitted for ease of presentation. (**b**,**d**,**f**) The plots show simulated and experimental cell confluency over 310 hours for various dose combinations of the ATRi and PARPi. Training data is used to estimate model parameters (plots I-V) and test data is used to evaluate the model (plot VI with the orange borders). The solid lines represent the simulation results from the three compartment models ((**b**) Model LC, (**d**) Model MC, (**f**) Model HC). The dashed lines represent the mean in vitro data, with standard errors for three experiments indicated with error bars.

We thereafter estimate the remaining parameters ($p_0$, $k_3'$, $s$, $E_{\max,1}$, $E_{50,1}$, $h_1$, $E_{\max,2}$, $E_{50,2}$, $h_2$) with the global optimiser GlobalSearch in MATLAB that minimises the sum of squares of the residuals between the model output and the in vitro data (Ugray et al., 2007). Model LC is described with the following system of ODEs

$$\frac{d[G1](t)}{dt} = 2k_4\big(U(t)\big)s[G2/M](t) - k_1\big(U(t)\big)s[G1](t), \tag{3a}$$

$$\frac{d[S](t)}{dt} = k_1\big(U(t)\big)sp[G1](t) - k_2\big(U(t)\big)s[S](t) + k_3\big(U(t)\big)q([\text{drug}_1],[\text{drug}_2])[SD](t), \tag{3b}$$

$$\frac{d[SD](t)}{dt} = k_1\big(U(t)\big)s(1-p)[G1](t) - k_3\big(U(t)\big)[SD](t), \tag{3c}$$

$$\frac{d[G2/M](t)}{dt} = k_2\big(U(t)\big)s[S](t) - k_4\big(U(t)\big)s[G2/M](t), \tag{3d}$$

$$\frac{d[NC](t)}{dt} = k_3\big(U(t)\big)(1-q([\text{drug}_1],[\text{drug}_2]))[SD](t). \tag{3e}$$

In summary, including initial conditions, Model LC includes 10 parameters that are directly estimated from the data and 9 parameters that are obtained via a global optimiser in MATLAB.

#### Medium Complexity Model

Model MC (Fig. S3**c**), is an expansion of Model LC and is described in Section 2.1 in the main manuscript, see especially Eqs. 1a-1e. See also Section 2.3 in the main manuscript on how the model parameters are estimated. In summary, including initial conditions, Model MC includes 10 parameters that are directly estimated from the data and 9 parameters that are obtained via a global optimiser in MATLAB.

#### High Complexity Model

Model HC is described in detail by Pugh et al. (2023). In summary, Model HC includes three undamaged and proliferative cell cycle states (G1, S, and G2/M), two damaged states which represent replication stress-induced DNA damage (SD) and DNA damage in the G2/M phase (G2D), and a dead state which represents cell death via apoptosis. The simulations start with no dead cells, and cells in the dead state remain there for the duration of the in silico simulation and do not contribute to the cell confluency. However, we assume that dead cells take up space and therefore, contribute to the carrying capacity. Cells can enter state G2D directly from state SD, which represents unrepaired or wrongly repaired replication stress (O'Connor, 2015). Since DNA damage can occur at any point during the cell cycle, cells can also transition to state G2D directly from state S at a rate that is proportional to a weighting factor $q$.

In the model, some of the rates depend on drug concentrations (Fig. S3**e**). After the G1 state, cells enter the damaged SD state at a rate that is proportional to a weighting factor $p$, and cells enter the S state at a rate that is proportional to $1-p$. The factor $p$ increases with increasing concentrations of the PARPi (drug 2), which captures the drug's replication repair-inhibiting effects. Cells in the damaged states SD and G2D can repair their DNA damage with rates $k_3$ and $k_7$ respectively. These rates decrease with increasing drug concentrations of the ATRi (drug 1) and the PARPi respectively, which capture the drugs replication repair-inhibiting effects. Cells in state G2D can transition to the dead state by an ATRi-related cell death. This represents the in vitro scenario where the ATRi releases cells from G2 arrest, which can cause mitotic catastrophe (Lloyd et al., 2020). In this model, we assume that this drug-induced cell death occurs at a delayed rate. Therefore, to incorporate the delay observed in the drug-induced cell death, we introduce a transit compartment into the model to represent death delay (Miao et al., 2016a; Lobo and Balthasar, 2002; Miao et al., 2016b). $A_1$ represents a time-dependent rate at which cells leave state G2D which increases over time to represent delayed cell death and $\tau$ represents the time spent in the delay. Cells that successfully progress to, and leave, the G2/M state re-renter the G1 state and produce a daughter cell that is initiated in state G1 (Fig. S3**e**).

The initial concentration of cells and the initial number of cells in compartments G1, S, SD, G2/M, and G2D ($G1_0$, $S_0$, $SD_0$, $G2/M_0$, $G2D_0$) are directly estimated from in vitro data. The parameters $k_1$, $k_2$, $k_3$, $k_4$, $k_5$, $k_6$, $k_7$, $k_8$, $p$, $q$, $L$, $\tau$, $I_{\max,C}$, $EC_{50,C}$, $h_{C,1}$, $K_{\max,C}$, $KC_{50,C}$, $h_{C,2}$, $E_{\max,O}$, $EC_{50,O}$, $h_{O,1}$, $I_{\max,O}$, $EC_{50,O}$, $h_{O,2}$ in Model HC are estimated by using the local optimiser lsqnonlin in MATLAB that minimises the sum of squares of the residuals between the model output and the in vitro data (Levenberg, 1944). Note that the values $p$ and $q$ have different meanings in Model HC compared to Model MC and Model LC. Model HC is described with the following system of ODEs

$$\frac{d[G1](t)}{dt} = 2k_5\left(1-\frac{N(t)}{L}\right)[G2/M](t) - k_1[G1](t), \tag{4a}$$

$$\frac{d[S](t)}{dt} = k_1(1 - p(1 + \text{Effect}_{\text{Ola},1}))[G1](t) - k_2[S](t) + k_3(1 - \text{Effect}_{\text{Cera},1})[SD](t), \tag{4b}$$

$$\frac{d[G2/M](t)}{dt} = k_2(1 - q)[S](t) - k_5\left(1 - \frac{N(t)}{L}\right)[G2/M](t) + k_7(1 - \text{Effect}_{\text{Ola},2})[G2D](t), \tag{4c}$$

$$\frac{d[SD](t)}{dt} = k_1 p(1 + \text{Effect}_{\text{Ola},1})[G1](t) - k_3(1 - \text{Effect}_{\text{Cera},1})[SD](t) - k_6[SD](t) - k_4[SD](t), \tag{4d}$$

$$\frac{d[G2D](t)}{dt} = k_2 q[S](t) + k_6[SD](t) - k_7(1 - \text{Effect}_{\text{Ola},2})[G2D](t) - k_8[G2D](t) - [A_1](t)[G2D](t), \tag{4e}$$

$$\frac{d[D](t)}{dt} = k_4[SD](t) + k_8[G2D](t) + [A_1](t)[G2D](t), \tag{4f}$$

$$\frac{d[A_1](t)}{dt} = \frac{1}{\tau}(\text{Effect}_{\text{Cera},2} - [A_1](t)). \tag{4g}$$

Where the drug effects are modelled by

$$\text{Effect}_{\text{Cera},1} = I_{\max,\text{C}} \cdot \left(\frac{ATRi^{h_{C,1}}}{IC_{50,\text{C}}^{h_{C,1}} + ATRi^{h_{C,1}}}\right), \tag{5a}$$

$$\text{Effect}_{\text{Cera},2} = K_{\max,\text{C}} \cdot \left(\frac{ATRi^{h_{C,2}}}{KC_{50,\text{C}}^{h_{C,2}} + ATRi^{h_{C,2}}}\right), \tag{5b}$$

$$\text{Effect}_{\text{Ola},1} = E_{\max,\text{O}} \cdot \left(\frac{PARPi^{h_{O,1}}}{EC_{50,\text{O}}^{h_{O,1}} + PARPi^{h_{O,1}}}\right), \tag{5c}$$

$$\text{Effect}_{\text{Ola},2} = I_{\max,\text{O}} \cdot \left(\frac{PARPi^{h_{O,2}}}{IC_{50,\text{O}}^{h_{O,2}} + PARPi^{h_{O,2}}}\right). \tag{5d}$$

In summary, including initial conditions, Model HC includes 6 parameters that are directly estimated from the data and 24 parameters that are obtained via a local optimiser in MATLAB.

#### Model Comparison

By visually comparing the cell confluency plots in Fig. S3 **b**, **d**, and **f**, we observe that Model HC is overfitted to the experimental data due to the high number of parameters. Model HC is also unable to match the data for the combination $[\text{drug}_1]$=0.3 $\mu$M and $[\text{drug}_2]$=1 $\mu$M, most likely because a local optimiser is used to parameterise the model. We argue that both Model LC and Model MC are able to satisfactorily predict unseen time series data. The RMSE for training/test data of Model LC and Model MC are, in units of cell coverage percentages, 3.2076%/0.7201% and 3.0751%/0.9462% respectively. We also note that Model MC includes more biologically relevant drug effects of the PARPi compared to Model LC. This is because Model MC includes both the single-strand break and double-strand break repair-inhibiting effects of the PARPi, whereas Model LC only includes the latter effect. It is commonly believed that the main function of PARPi drugs is to inhibit the repair of single-strand breaks which cause double-strand breaks (Chen, 2011; Ashworth, 2008). Yet, the effect of PARP on double-strand break repair is heavily disputed. While some authors argue that PARP is involved in the repair of double-strand breaks (Caron et al., 2019; Chaudhuri and Nussenzweig, 2017), other authors argue that PARP is not involved in the repair of double-strand breaks (Noël et al., 2003; Ashworth, 2008). Therefore, to safeguard our model, we argue that Model MC is the better model to study cell population dynamics in response to 0.1-1 $\mu$M ceralasertib doses and 0.2-1 $\mu$M olaparib doses.

#### S4 A consistency analysis is conducted

We perform a consistency analysis for the ABM to conclude that 100 simulation runs suffice to form the basis for our reported results. For this study, we perform a consistency analysis on the cell crowding limited ABM where cells are seeded on the lattice in single-cell clusters and no drug-resistant cells are included. We choose two output variables of interest, $X_1$ and $X_2$, regarding the cell confluency with $[\text{drug}_1] = 0.3$ $\mu$M and $[\text{drug}_2] = 0.2$ $\mu$M. $X_1$ is the cell confluency halfway through the simulations and $X_2$ is the cell confluency at the end of the simulations. We choose this combination treatment to ensure we have stochasticity from both drugs, without non-cycling cells having a significant effect on the cell confluency output. We seek the smallest number of in silico simulations, $d$, needed to be run per experiment that mitigates uncertainty originating from intrinsic model stochasticity, i.e., the smallest $d$ to produce a small statistical significance. In this study, we test values of $d$ = 1, 5, 50, 100, and 300.

The results of this analysis are summarised in Fig. S4 and an overview of the procedure is outlined in Section S4.1. The procedure follows that outlined by Hamis et al. (2021a) which builds on previous statistical work by Vargha et al. (2000) and Alden et al. (2013).

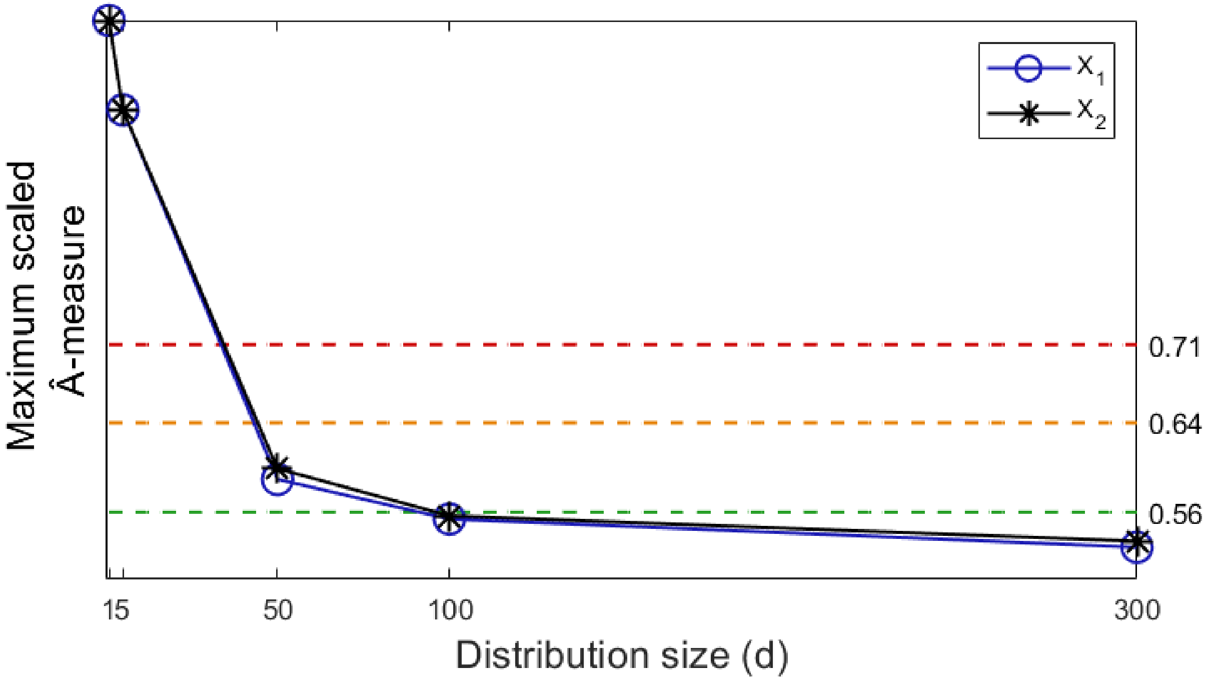

Fig. S4: **A consistency analysis is performed to decide how many in silico simulations are needed to make conclusions from the ABM to mitigate any model uncertainty arising from intrinsic-model stochasticity.** The figure shows the maximum scaled $\hat{A}$-measure (vertical axis) for two output variables $X_1$ and $X_2$ for different distribution sizes $d$ (horizontal axis). $X_1$ represents the cell confluency halfway through the simulation and $X_2$ represents the cell confluency at the end of the simulation both with $[\text{drug}_1] = 0.3$ $\mu$M and $[\text{drug}_2] = 0.2$ $\mu$M. We seek the smallest $d$ for which the maximum scaled $\hat{A}$-measure for both outputs $X_1$ and $X_2$ are below 0.56 (under the green line).

##### S4.1 Overview of the consistency analysis procedure

When conducting consistency analysis, one in silico simulation produces one data sample of the chosen variables $X_i$, $i = 1, 2$. We generate distributions of data samples with varying sizes, where a distribution with distribution-size $d$ has $d$ data samples. We produce distribution groups, $g_d$, each comprising 20 distributions of distribution-size $d$, i.e., a group, $g_1$, comprising 20 distributions of size $d = 1$, another group $g_5$ comprising 20 distributions of size $d = 5$, etc. Overall, we have to produce $20(1 + 5 + 50 + 100 + 300) = 9120$ data samples to perform the consistency analysis. In each distribution group, $g_d$, we compare the distributions 1 and $k'$, $k' = 2, 3, \ldots, 20$, for each variable $X_i$ to deduce how stochastically equal the two distributions are. This can be done by computing the $\hat{A}$-measure, denoted $\hat{A}^{g_d}_{1,k'}(X_i)$ (Eqs. 6 and 7) (Hamis et al., 2021a).

Let $B$ and $C$ be two distributions comprising $m_1$ and $m_2$ data samples respectively of some variable $X$ such that $B = \{b_1, b_2, \ldots b_{m_1}\}$ and $C = \{c_1, c_2, \ldots c_{m_2}\}$. The $\hat{A}$-measure is calculated as follows

$$\hat{A}_{BC}(X) = \frac{1}{m_1 m_2} \sum_{i=1}^{m_1} \sum_{j=1}^{m_2} H(b_i - c_j), \tag{6}$$

where

$$H(x) = \begin{cases} 1 & \text{if } x > 0, \\ \frac{1}{2} & \text{if } x = 0, \\ 0 & \text{if } x < 0. \end{cases} \tag{7}$$

In other words, we compare all values $b_i$, $i = 1, \ldots, m_1$ in distribution $B$ to values $c_j$, $j = 1, \ldots, m_2$ in distribution $C$ and give them a score of 0, 0.5, or 1. To deduce how stochastically equal distributions B and C are, we calculate the $\hat{A}$-measure, $\hat{A}_{BC}(X) \in [0, 1]$, by averaging the sum of the scores by dividing by $m_1 m_2$. If all values $b_i$ are bigger than all values $c_j$, the $\hat{A}$-measure value is 1. Similarly, if all values $b_i$ are smaller than all values $c_j$, the $\hat{A}$-measure value is 0. The closer the $\hat{A}$-measure value is to 0.5, i.e., when the values $b_i$ are bigger than $c_j$ half of the time, the more stochastically equal distributions $B$ and $C$ are with regard to the variable $X$. Here, we are only interested in how much the distributions are equal, not which distribution is stochastically greater. Therefore, to omit direction we calculate the scaled $\hat{A}$-measure, denoted $\underline{\hat{A}} \in [0.5, 1]$ (Hamis et al., 2021a), given by

$$\underline{\hat{A}}_{BC}(X) = \begin{cases} \hat{A}_{BC}(X) & \text{if } \hat{A}_{BC}(X) \geq 0.5, \\ 1 - \hat{A}_{BC}(X) & \text{if } \hat{A}_{BC}(X) < 0.5. \end{cases} \tag{8}$$

For each group, $g_d$, we calculate the maximal scaled $\hat{A}$-measure, $\underline{\hat{A}}^{g_d}_{\max}$, and compare it to threshold values (Eq. 9) to deduce whether the statistical significance between the two distributions is small, medium, or large (Hamis et al., 2021a; Cohen, 1988). The threshold values are given by

$$\text{Statistical Significance} = \begin{cases} \text{small} & \text{if } \underline{\hat{A}}^{g_d}_{\max}(X) \in [0.5, 0.56], \\ \text{medium} & \text{if } \underline{\hat{A}}^{g_d}_{\max}(X) \in (0.56, 0.64], \\ \text{large} & \text{if } \underline{\hat{A}}^{g_d}_{\max}(X) \in (0.64, 0.71]. \end{cases} \tag{9}$$

Hence, we want to find the smallest value of $d$ such that $\underline{\hat{A}}^{g_d}_{\max} \leq 0.56$, i.e., the smallest distribution-size $d$ yielding a small stochastic significance for all 19 $\hat{A}$-measures (Hamis et al., 2021a).

We conclude from Fig. S4 and Table S3 that $d = 100$ is the smallest tested value of $d$ that has a maximum scaled $\hat{A}$-measure value under the threshold value 0.56 for both outputs $X_1$ and $X_2$. Therefore, 100 simulation runs per in silico experiment will suffice to produce a small enough statistical significance to enable us to draw conclusions from the ABM. Note that Fig. S5 shows that smaller values of $d$ result in a more irregular pattern in the scaled and initial $\hat{A}$-measure values. The $\hat{A}$-measure values become more consistent with a small statistical significance for values $d = 100$ and 300.

| Distribution Size | $d = 1$ | $d = 5$ | $d = 50$ | $d = 100$ | $d = 300$ |
|---|---|---|---|---|---|
| Output $X_1$ (half time) | 1 | 0.92 | 0.5894 | 0.5539 | 0.5287 |
| Output $X_2$ (end time) | 1 | 0.92 | 0.5990 | 0.5569 | 0.5341 |

Table S3: **Scaled $\hat{A}$-measure calculate how stochastically equal two distributions are.** The table shows the scaled $\hat{A}$-measure values for distribution sizes $d$ = 1, 5, 50, 100, and 300 relating to two output variables. $X_1$ represents the cell confluency halfway through the simulation and $X_2$ represents the cell confluency at the end of the simulation with [drug$_1$] = 0.3 $\mu$M and [drug$_2$] = 0.2 $\mu$M.

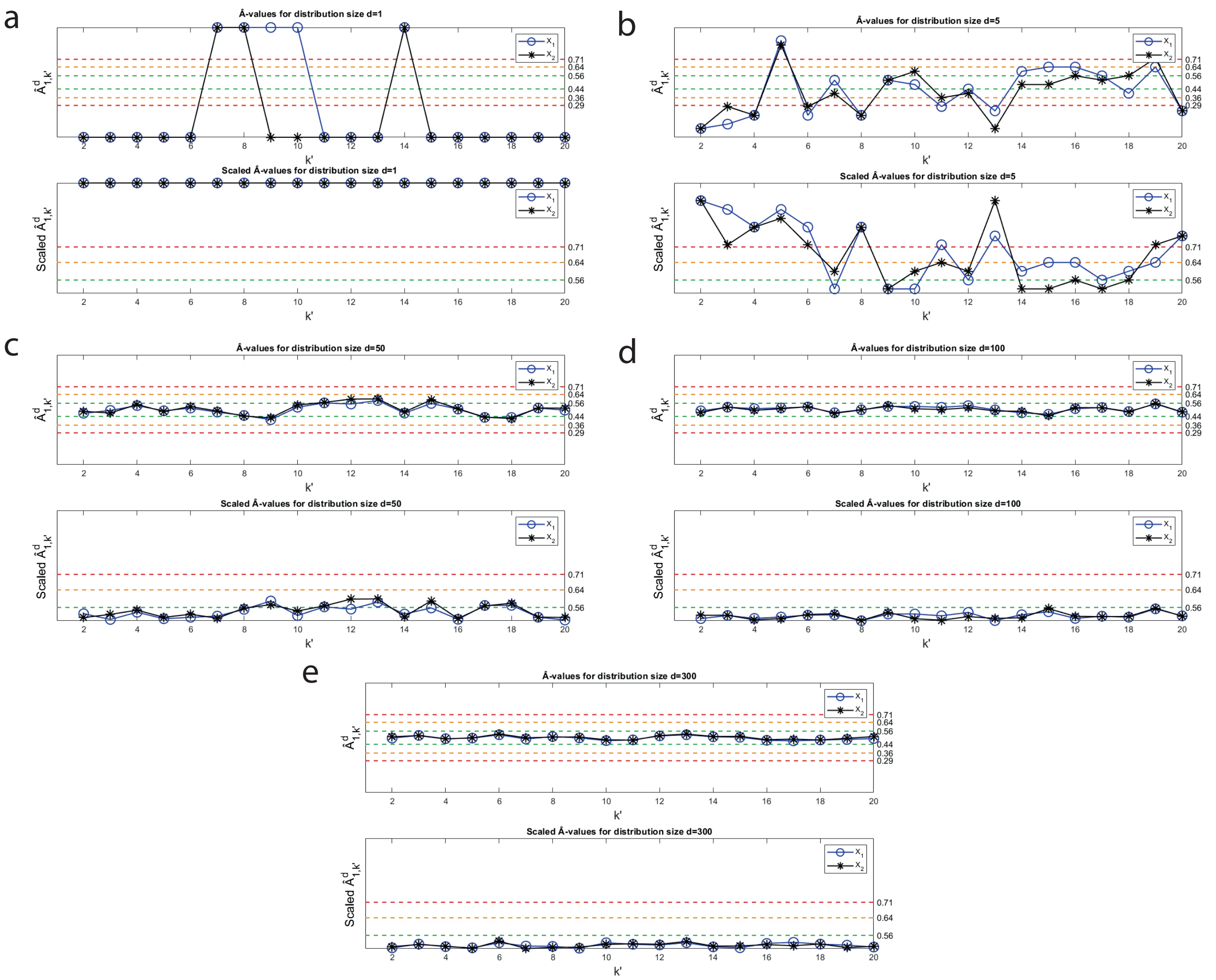

Fig. S5: **A consistency analysis is performed to decide how many in silico simulations are enough to draw conclusions from the ABM to mitigate any model uncertainty arising from intrinsic-model stochasticity.** Each figure displays the $\hat{A}$-measures (top) and the scaled $\hat{A}$-measures (bottom) for distribution sizes (**a**) $d = 1$, (**b**) $d = 5$, (**c**) $d = 50$, (**d**) $d = 100$, and (**e**) $d = 300$. $X_1$ represents the cell confluency halfway through the simulation and $X_2$ represents the cell confluency at the end of the simulation with [drug$_1$] = 0.3 $\mu$M and [drug$_2$] = 0.2 $\mu$M.

#### S5 End time cell configurations are visualised

In Fig. S6, we show how the spatial organisation of cells at the end of the simulation is impacted by different drug treatments and doses while initialised with the same spatial configuration.

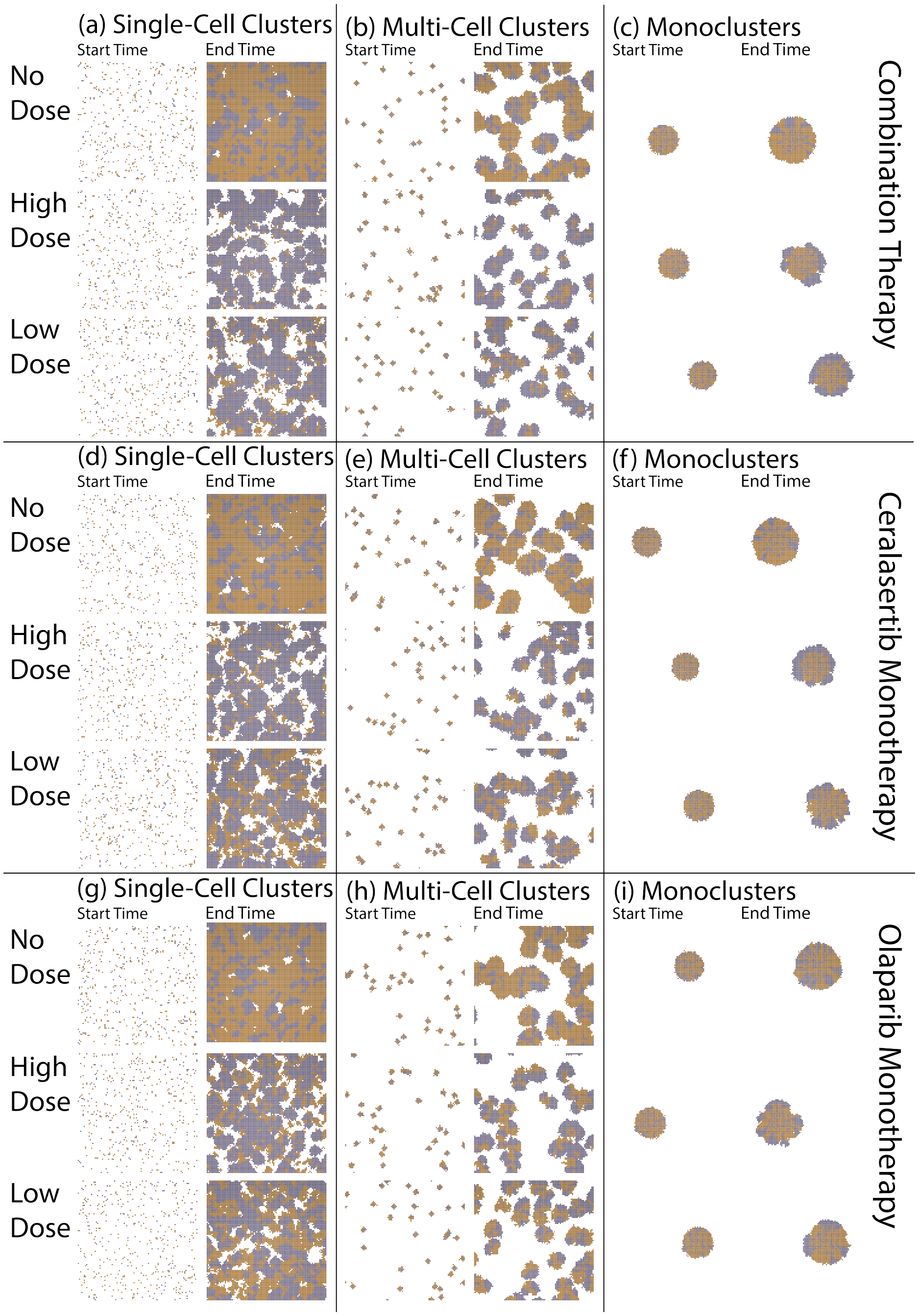

Fig. S6: **The spatial organisation of cells with and without treatment.** In (**a**-**i**), we seed a total number of $P_0$ cells on the lattice, where $0.7P_0$ cells are drug-sensitive (orange) and $0.3P_0$ cells are drug-resistant (purple). Cells are seeded in single-cell clusters (**a**,**d**,**g**), multi-cell clusters (**b**,**e**,**h**), or monoclusters (**c**,**f**,**i**). Simulations are performed with no drugs (top row in each panel **a-i**), combination therapy (**a**,**b**,**c**), ceralasertib monotherapy (**d**,**e**,**f**), and olaparib monotherapy (**g**,**h**,**i**) for high (middle row in each panel **a-i**) and low (bottom row in each panel **a-i**) doses. Examples of initial and final simulation snapshots are shown in each panel.

#### S6 Experiments are repeated where NC cells are removed from the lattice and G0 cells can re-enter the cell cycle

The experiments performed in Section 3 of the main manuscript (Figs. 3-7) are repeated for the case where: (1) NC cells are removed from the lattice after one cell cycle and (2) cells in state G0 can re-enter the cell cycle, initiated in state G1, if a free lattice site becomes available in its first order-neighbourhood. Without cell removal, clusters develop a quiescent inner core as only cells on the periphery can proliferate. However, the quiescent core is not as apparent with NC cell removal because cell removal frees up lattice sites for cells in state G0 to re-enter the cell cycle (see Fig. S7). Therefore, incorporating the new modelling rules significantly impacts cell population dynamics, particularly in larger-sized clusters (see Figs. S8-S12). These additional rules lead to a significantly higher fraction of drug-resistant cells in both multi-cell clusters and monoclusters compared to the results obtained without them.

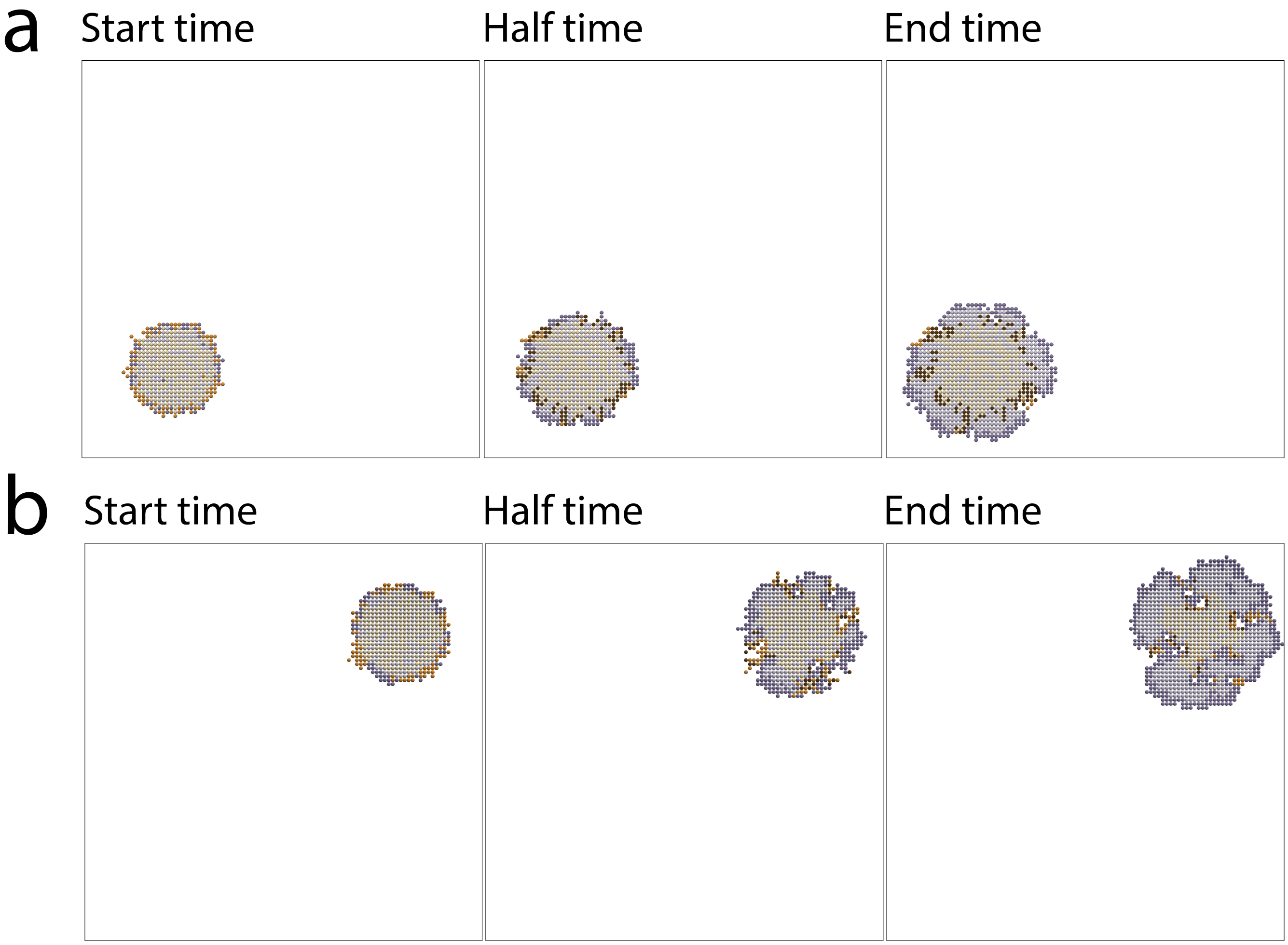

Fig. S7: **The cell crowding limited ABM is compared to an updated version in which NC cells are removed and quiescent cells can re-enter the cell cycle.** We seed a total number of $P_0$ cells on the lattice, where $0.7P_0$ cells are drug-sensitive (orange) and $0.3P_0$ cells are drug-resistant (purple). Examples of initial (left), half time (middle), and final (right) simulation snapshots are shown for (**a**) the original cell crowding limited ABM and (**b**) the cell crowding limited ABM with the updated rules. Darker and lighter colours represent cells in state NC and state G0 cells respectively.

#### S6.1 Spatial cell structures impact dynamic treatment responses

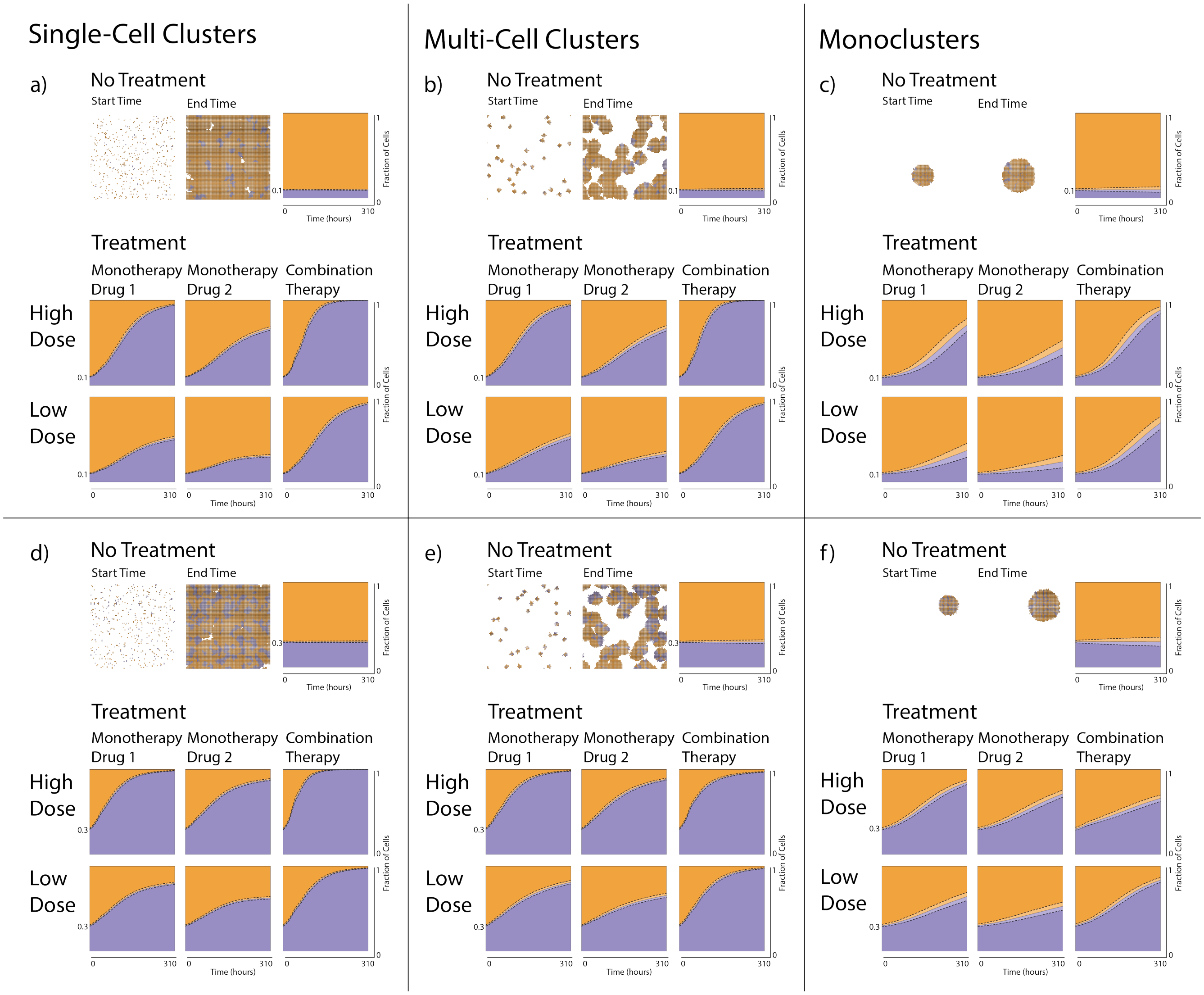

Fig. S8: **Cells that are resistant to both an ATRi (drug 1) and a PARPi (drug 2) compete for spatial resources with drug-sensitive cells when cells in state NC are removed after one cell cycle and cells in state G0 can re-enter the cell cycle.** A total number of $P_0$ cells are randomly seeded on the lattice in single-cell clusters (**a**,**d**), multi-cell clusters (**b**,**e**), or monoclusters (**c**,**f**), with either $0.1P_0$ (**a**,**b**,**c**) or $0.3P_0$ (**d**,**e**,**f**) drug-resistant cells. In each panel **a-f**, results are shown from simulations with no drugs (top row), drug 1 monotherapy (left column in each treatment panel), drug 2 monotherapy (middle column in each treatment panel), and combination therapy (right column in each treatment panel) for high (middle row) and low (bottom row) doses. Examples of initial and final simulation snapshots are shown in each panel. Drug-sensitive and drug-resistant cells are, respectively, orange and purple. The panels also include plots of the dynamic mean fraction of drug-sensitive (orange) and drug-resistant (purple) cells, and standard deviations (dashed lines) from 100 simulation runs.

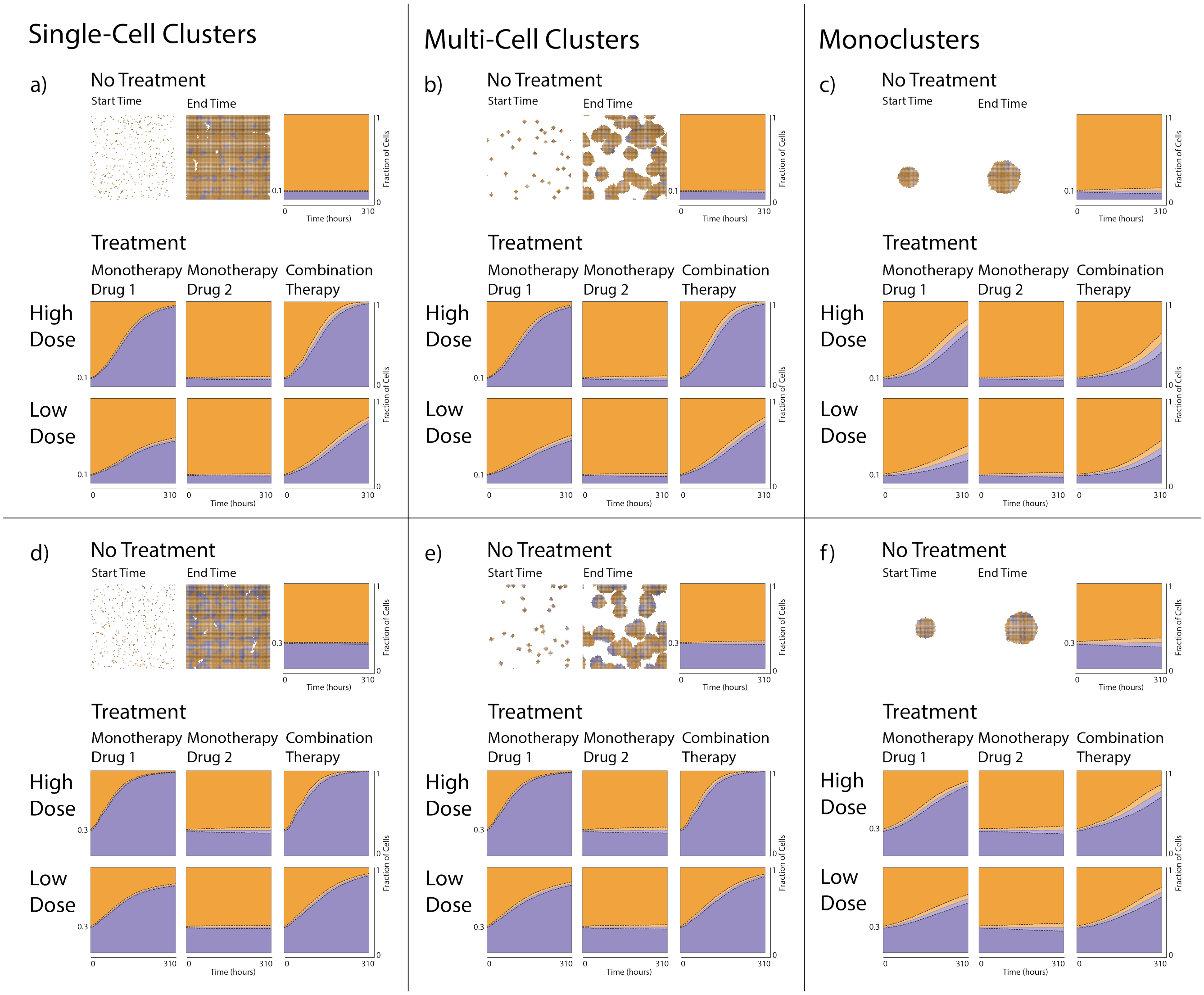

Fig. S9: **Cells that are resistant to an ATRi (drug 1) compete for spatial resources with drug-sensitive cells when cells in state NC are removed after one cell cycle and cells in state G0 can re-enter the cell cycle.** A total number of $P_0$ cells are randomly seeded on the lattice in single-cell clusters (**a**,**d**), multi-cell clusters (**b**,**e**), or monoclusters (**c**,**f**), with either $0.1P_0$ (**a**,**b**,**c**) or $0.3P_0$ (**d**,**e**,**f**) drug-resistant cells. In each panel **a-f**, results are shown from simulations with no drugs (top row), drug 1 monotherapy (left column in each treatment panel), drug 2 monotherapy (middle column in each treatment panel), and combination therapy (right column in each treatment panel) for high (middle row) and low (bottom row) doses. Examples of initial and final simulation snapshots are shown in each panel. Drug-sensitive and drug-resistant cells are, respectively, orange and purple. The panels also include plots of the dynamic mean fraction of drug-sensitive (orange) and drug-resistant (purple) cells, and standard deviations (dashed lines) from 100 simulation runs.

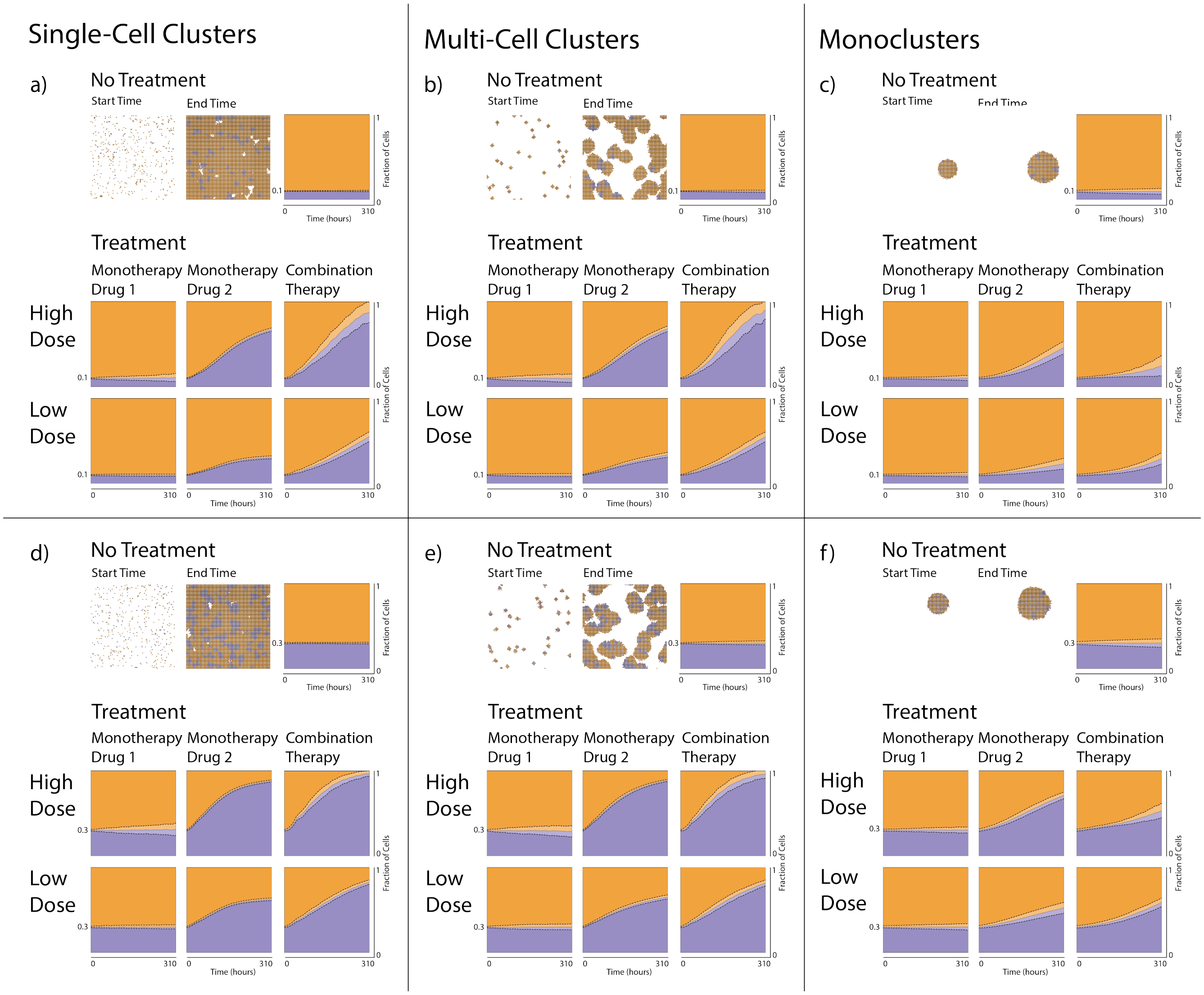

Fig. S10: **Cells that are resistant to a PARPi (drug 2) compete for spatial resources with drug-sensitive cells when cells in state NC are removed after one cell cycle and cells in state G0 can re-enter the cell cycle.** A total number of $P_0$ cells are randomly seeded on the lattice in single-cell clusters (**a**,**d**), multi-cell clusters (**b**,**e**), or monoclusters (**c**,**f**), with either $0.1P_0$ (**a**,**b**,**c**) or $0.3P_0$ (**d**,**e**,**f**) drug-resistant cells. In each panel **a-f**, results are shown from simulations with no drugs (top row), drug 1 monotherapy (left column in each treatment panel), drug 2 monotherapy (middle column in each treatment panel), and combination therapy (right column in each treatment panel) for high (middle row) and low (bottom row) doses. Examples of initial and final simulation snapshots are shown in each panel. Drug-sensitive and drug-resistant cells are, respectively, orange and purple. The panels also include plots of the dynamic mean fraction of drug-sensitive (orange) and drug-resistant (purple) cells, and standard deviations (dashed lines) from 100 simulation runs.

#### S6.2 Spatial cell structures impact the feasibility of treatment objectives

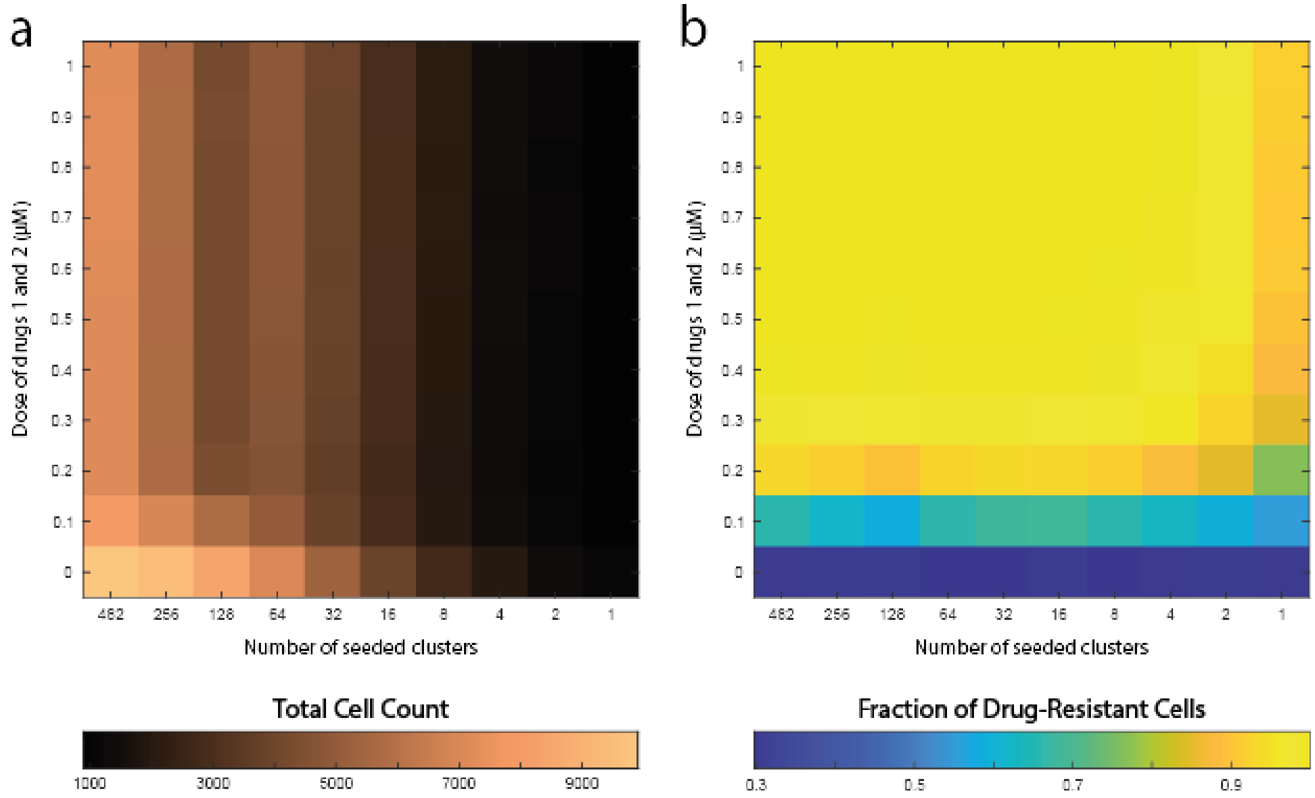

Fig. S11: **Drug doses and spatial cell configurations impact total cell counts and the composition between drug-sensitive and drug-resistant cells when cells in state NC are removed after one cell cycle and cells in state G0 can re-enter the cell cycle.** The heatmaps show results from in silico experiments in which drug-sensitive and drug-resistant cells coexist. At the start of the experiments, a total number of $P_0$ cells are seeded on the lattice, where $0.3P_0$ cells are resistant to both drugs 1 and 2. Two inputs are varied in the simulations: (1) the number of clusters in which cells are seeded (indicated by the horizontal heatmap axes), and (2) the combination treatment drug doses (indicated by the vertical heatmap axes). The results show (**a**) the total cell count and (**b**) the fraction of drug-resistant cells at the end of the simulations. Each heatmap bin shows the mean value of 100 simulation runs.

#### S6.3 Spatial cell structures impact doubling time associated treatment responses

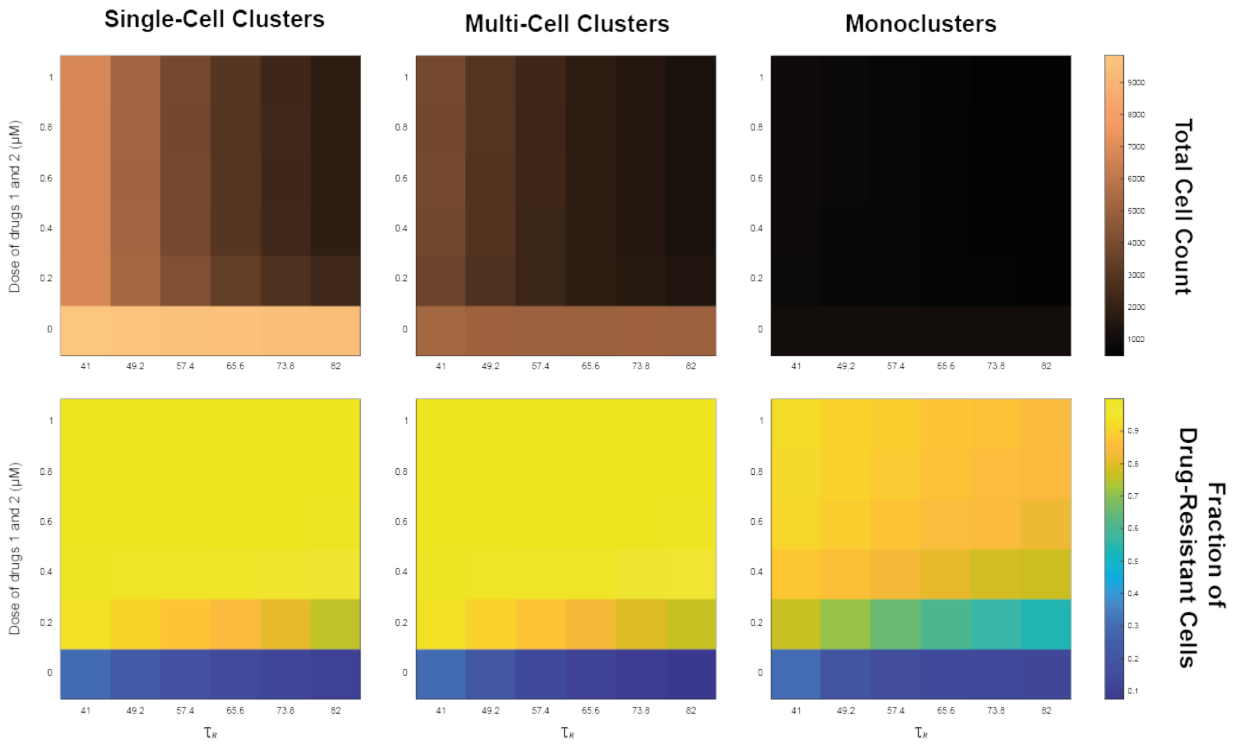

Fig. S12: **Spatial cell configurations, drug doses, and cell doubling times impact total cell counts and the composition between drug-sensitive and drug-resistant cells when cells in state NC are removed after one cell cycle and cells in state G0 can re-enter the cell cycle.** At the start of the experiments, a total number of $P_0$ cells are seeded on the lattice, where $0.3P_0$ cells are resistant to both drugs 1 and 2. Three inputs are varied in the simulations: (1) the seeded cluster configurations (left, middle, right panel), (2) the combination treatment drug doses (indicated by the vertical heatmap axes), and (3) the mean doubling time of the drug-resistant cells (indicated by the horizontal heatmap axes). The heatmaps show the total cell count (top panel) and the fraction of drug-resistant cells (bottom panel) at the end of the simulation. Each heatmap bin shows the mean value of 100 simulation runs.

### S7 Experiments are repeated where daughter cells can be placed in higher order neighbourhoods

#### S7.1 Daughter cells can be placed in 1st- and 2nd-order neighbourhoods

In Figs. S13-S17, the experiments performed in Section 3 of the main manuscript (Figs. 3-7) are repeated for the case where daughter cells may be placed in their 1st- and 2nd-order neighbourhoods ($V_c = 2$).

##### S7.1.1 Spatial cell structures impact dynamic treatment responses

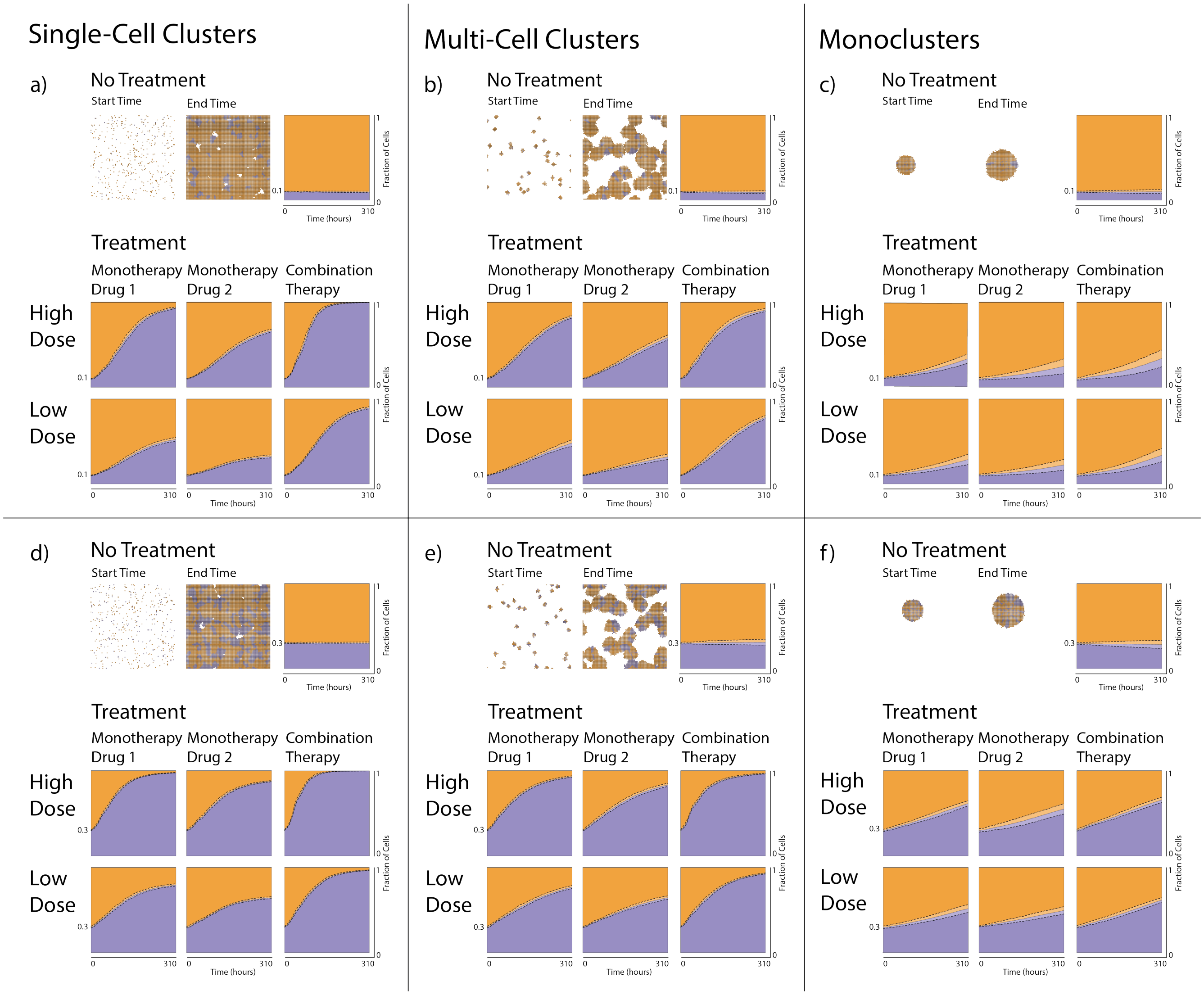

Fig. S13: **Cells that are resistant to both an ATRi (drug 1) and a PARPi (drug 2) compete for spatial resources with drug-sensitive cells when daughter cells can be placed in 1st- and 2nd-order neighbourhoods.** A total number of $P_0$ cells are randomly seeded on the lattice in single-cell clusters (**a**,**d**), multi-cell clusters (**b**,**e**), or monoclusters (**c**,**f**), with either $0.1P_0$ (**a**,**b**,**c**) or $0.3P_0$ (**d**,**e**,**f**) drug-resistant cells. In each panel **a-f**, results are shown from simulations with no drugs (top row), drug 1 monotherapy (left column in each treatment panel), drug 2 monotherapy (middle column in each treatment panel), and combination therapy (right column in each treatment panel) for high (middle row) and low (bottom row) doses. Examples of initial and final simulation snapshots are shown in each panel. Drug-sensitive and drug-resistant cells are, respectively, orange and purple. The panels also include plots of the dynamic mean fraction of drug-sensitive (orange) and drug-resistant (purple) cells, and standard deviations (dashed lines) from 10 simulation runs.

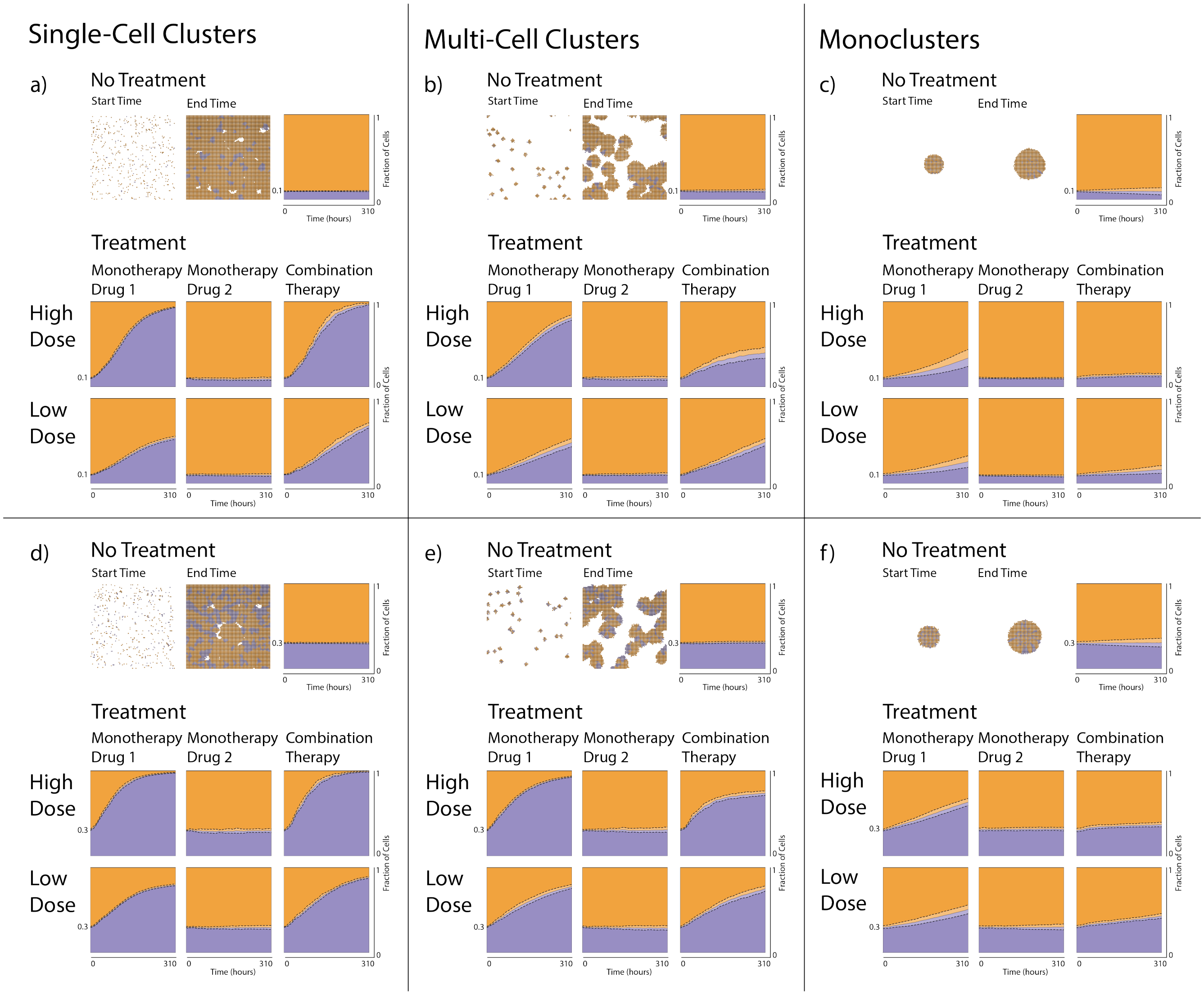

Fig. S14: **Cells that are resistant to an ATRi (drug 1) compete for spatial resources with drug-sensitive cells when daughter cells can be placed in 1st- and 2nd-order neighbourhoods.** A total number of $P_0$ cells are randomly seeded on the lattice in single-cell clusters (**a**,**d**), multi-cell clusters (**b**,**e**), or monoclusters (**c**,**f**), with either $0.1P_0$ (**a**,**b**,**c**) or $0.3P_0$ (**d**,**e**,**f**) drug-resistant cells. In each panel **a-f**, results are shown from simulations with no drugs (top row), drug 1 monotherapy (left column in each treatment panel), drug 2 monotherapy (middle column in each treatment panel), and combination therapy (right column in each treatment panel) for high (middle row) and low (bottom row) doses. Examples of initial and final simulation snapshots are shown in each panel. Drug-sensitive and drug-resistant cells are, respectively, orange and purple. The panels also include plots of the dynamic mean fraction of drug-sensitive (orange) and drug-resistant (purple) cells, and standard deviations (dashed lines) from 10 simulation runs.

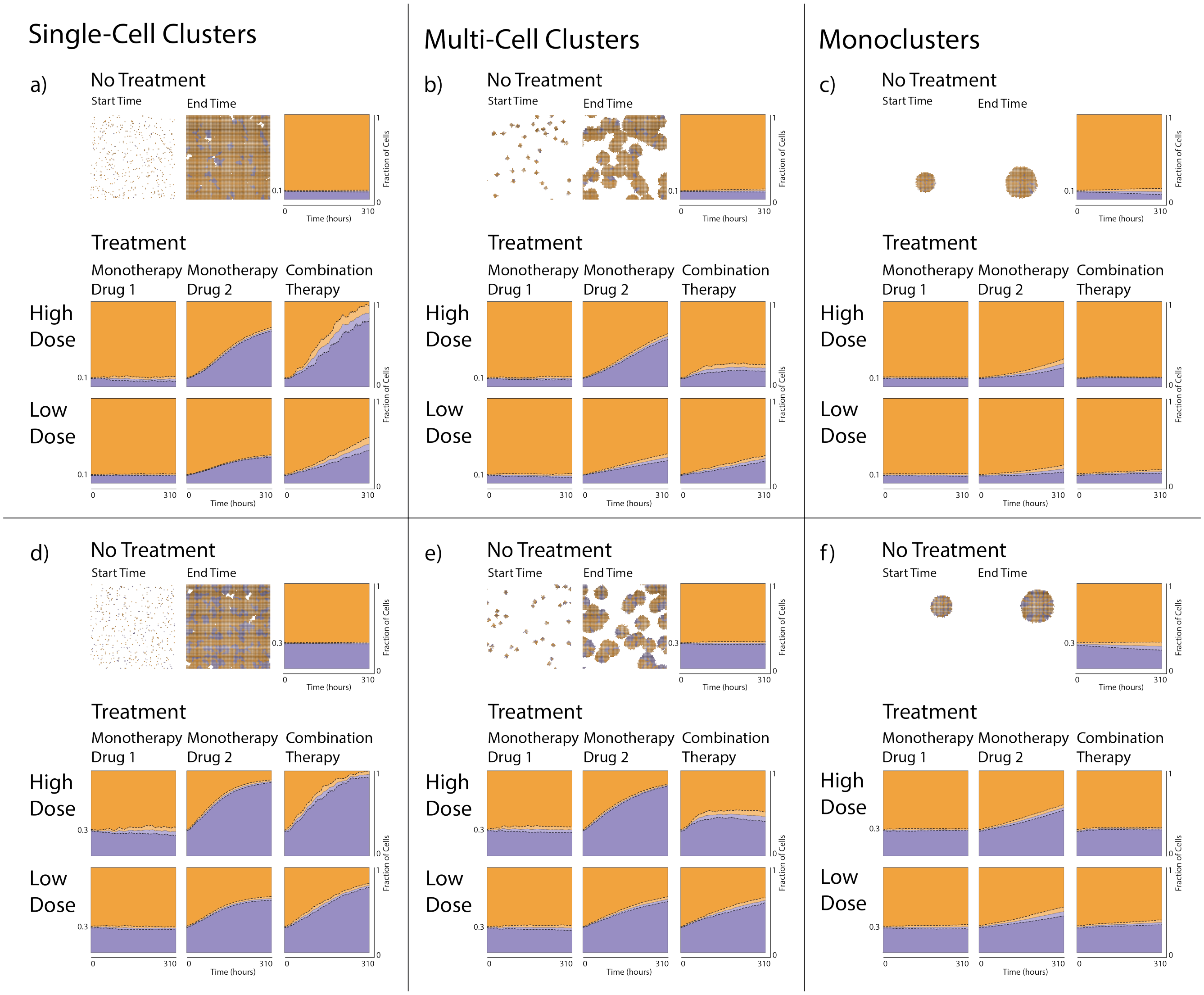

Fig. S15: **Cells that are resistant to a PARPi (drug 2) compete for spatial resources with drug-sensitive cells when daughter cells can be placed in 1st- and 2nd-order neighbourhoods.** A total number of $P_0$ cells are randomly seeded on the lattice in single-cell clusters (**a**,**d**), multi-cell clusters (**b**,**e**), or monoclusters (**c**,**f**), with either $0.1P_0$ (**a**,**b**,**c**) or $0.3P_0$ (**d**,**e**,**f**) drug-resistant cells. In each panel **a-f**, results are shown from simulations with no drugs (top row), drug 1 monotherapy (left column in each treatment panel), drug 2 monotherapy (middle column in each treatment panel), and combination therapy (right column in each treatment panel) for high (middle row) and low (bottom row) doses. Examples of initial and final simulation snapshots are shown in each panel. Drug-sensitive and drug-resistant cells are, respectively, orange and purple. The panels also include plots of the dynamic mean fraction of drug-sensitive (orange) and drug-resistant (purple) cells, and standard deviations (dashed lines) from 10 simulation runs.

###### S7.1.2 Spatial cell structures impact the feasibility of treatment objectives

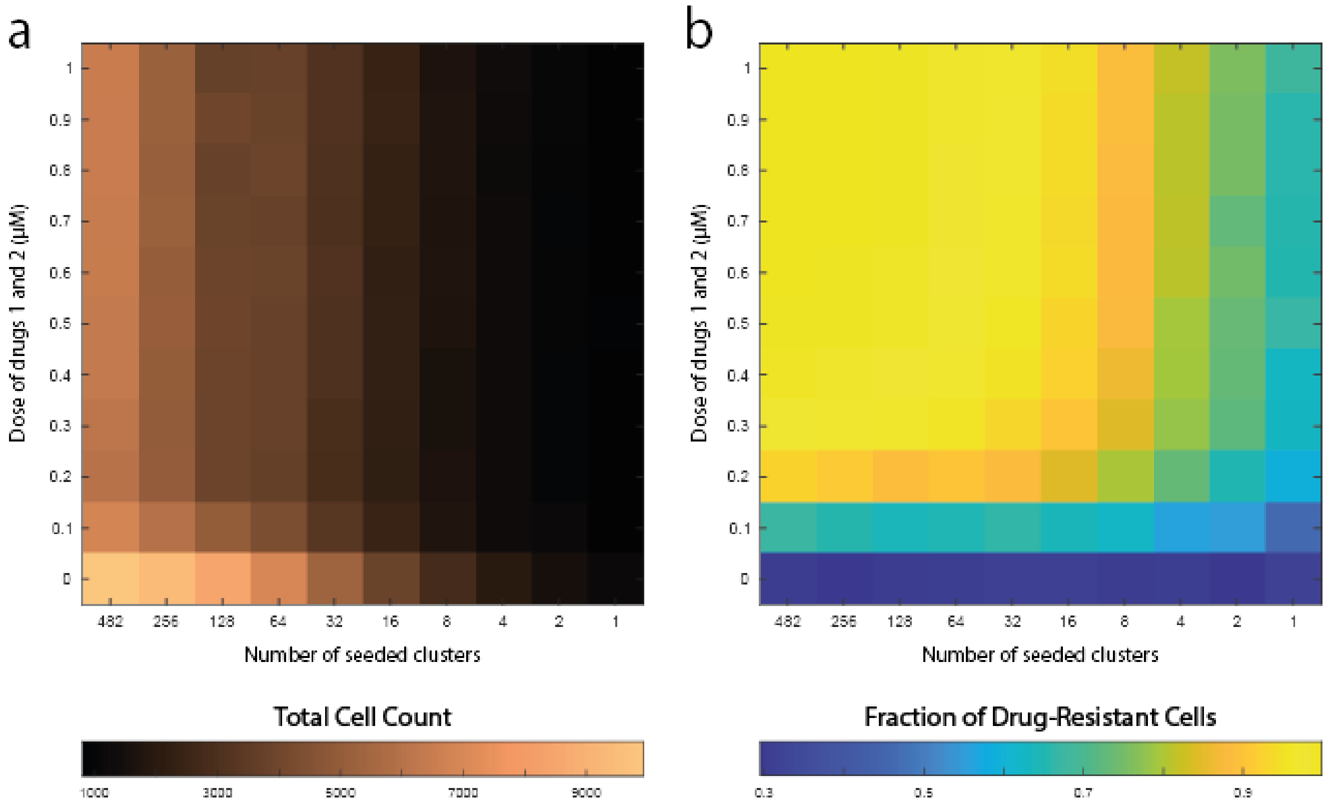

Fig. S16: **Drug doses and spatial cell configurations impact total cell counts and the composition between drug-sensitive and drug-resistant cells when daughter cells can be placed in 1st- and 2nd-order neighbourhoods.** The heatmaps show results from in silico experiments in which drug-sensitive and drug-resistant cells coexist. At the start of the experiments, a total number of $P_0$ cells are seeded on the lattice, where $0.3P_0$ cells are resistant to both drugs 1 and 2. Two inputs are varied in the simulations: (1) the number of clusters in which cells are seeded (indicated by the horizontal heatmap axes), and (2) the combination treatment drug doses (indicated by the vertical heatmap axes). The results show (**a**) the total cell count and (**b**) the fraction of drug-resistant cells at the end of the simulations. Each heatmap bin shows the mean value of 10 simulation runs.

##### S7.1.3 Spatial cell structures impact doubling time associated treatment responses

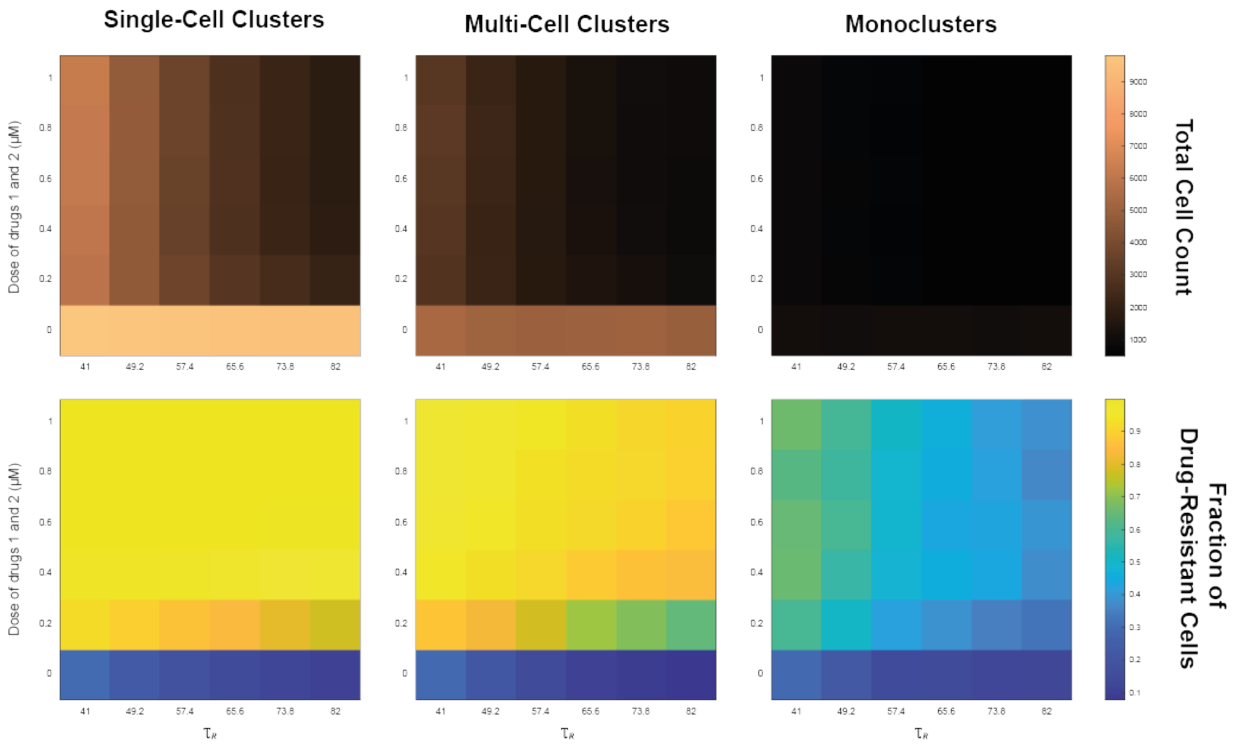

Fig. S17: **Spatial cell configurations, drug doses, and cell doubling times impact total cell counts and the composition between drug-sensitive and drug-resistant cells when daughter cells can be placed in 1st- and 2nd-order neighbourhoods.** At the start of the experiments, a total number of $P_0$ cells are seeded on the lattice, where $0.3P_0$ cells are resistant to both drugs 1 and 2. Three inputs are varied in the simulations: (1) the seeded cluster configurations (left, middle, right panel), (2) the combination treatment drug doses (indicated by the vertical heatmap axes), and (3) the mean doubling time of the drug-resistant cells (indicated by the horizontal heatmap axes). The heatmaps show the total cell count (top panel) and the fraction of drug-resistant cells (bottom panel) at the end of the simulation. Each heatmap bin shows the mean value of 10 simulation runs.

#### S7.2 Daughter cells can be placed in 1st-, 2nd- and 3rd-order neighbourhoods

In Figs. S18-S22, the experiments performed in Section 3 of the main manuscript (Figs. 3-7) are repeated for the case where daughter cells may be placed in their 1st-, 2nd-, and 3rd-order neighbourhoods ($V_c = 3$).

##### S7.2.1 Spatial cell structures impact dynamic treatment responses

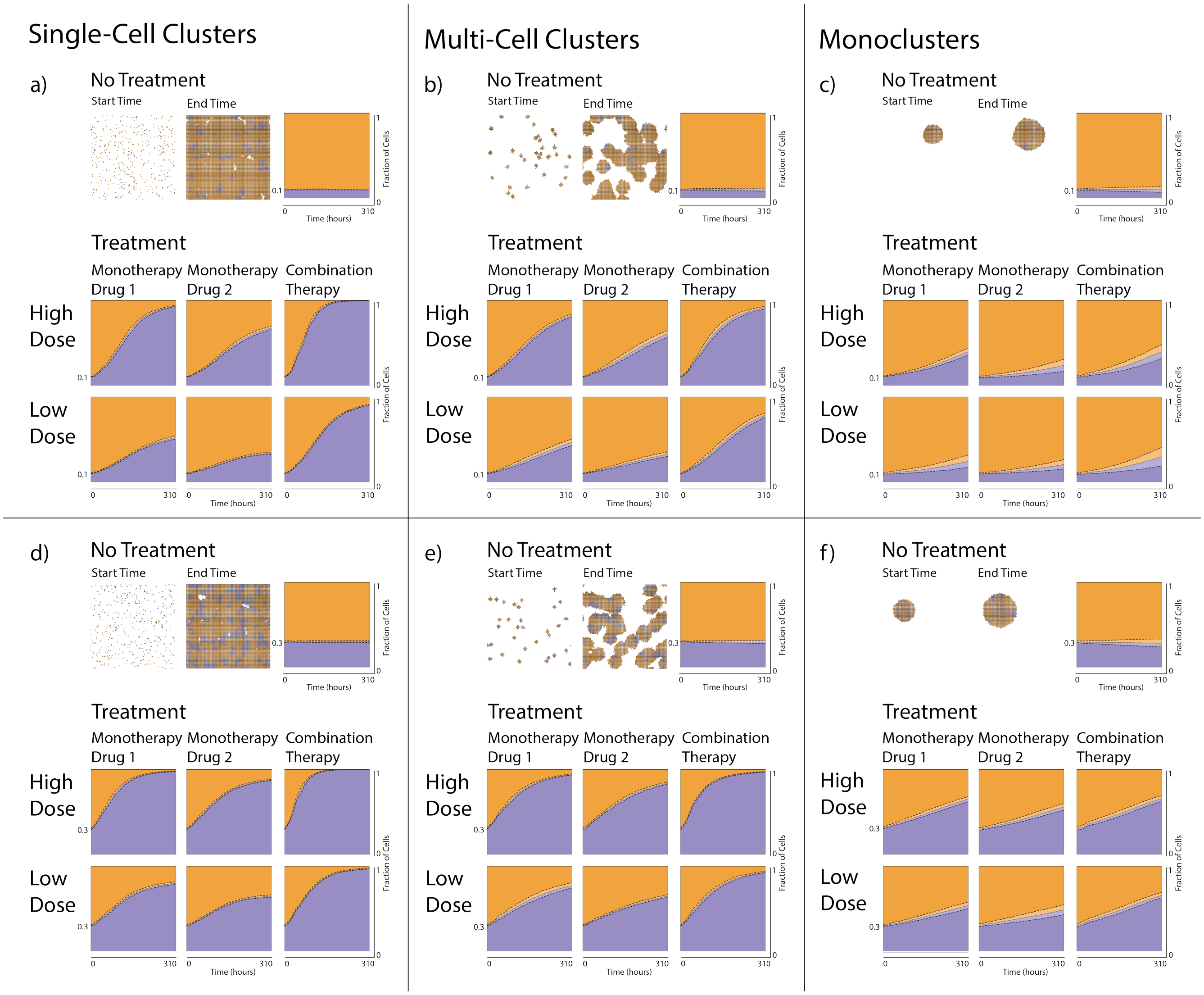

Fig. S18: **Cells that are resistant to both an ATRi (drug 1) and a PARPi (drug 2) compete for spatial resources with drug-sensitive cells when daughter cells can be placed in 1st-, 2nd-, and 3rd-order neighbourhoods.** A total number of $P_0$ cells are randomly seeded on the lattice in single-cell clusters (**a**,**d**), multi-cell clusters (**b**,**e**), or monoclusters (**c**,**f**), with either $0.1P_0$ (**a**,**b**,**c**) or $0.3P_0$ (**d**,**e**,**f**) drug-resistant cells. In each panel **a-f**, results are shown from simulations with no drugs (top row), drug 1 monotherapy (left column in each treatment panel), drug 2 monotherapy (middle column in each treatment panel), and combination therapy (right column in each treatment panel) for high (middle row) and low (bottom row) doses. Examples of initial and final simulation snapshots are shown in each panel. Drug-sensitive and drug-resistant cells are, respectively, orange and purple. The panels also include plots of the dynamic mean fraction of drug-sensitive (orange) and drug-resistant (purple) cells, and standard deviations (dashed lines) from 10 simulation runs.

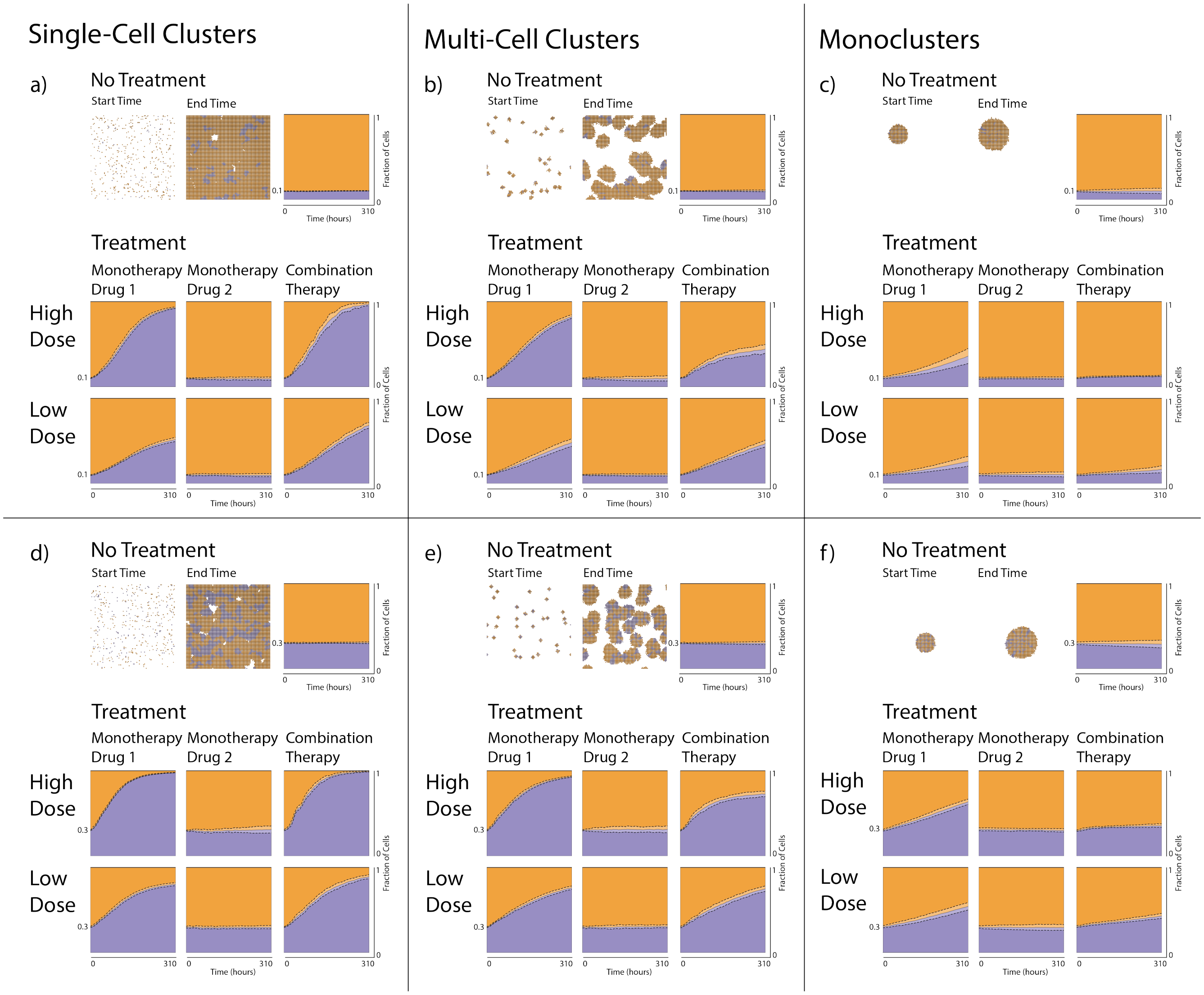

Fig. S19: **Cells that are resistant to an ATRi (drug 1) compete for spatial resources with drug-sensitive cells when daughter cells can be placed in 1st-, 2nd-, and 3rd-order neighbourhoods.** A total number of $P_0$ cells are randomly seeded on the lattice in single-cell clusters (**a**,**d**), multi-cell clusters (**b**,**e**), or monoclusters (**c**,**f**), with either $0.1P_0$ (**a**,**b**,**c**) or $0.3P_0$ (**d**,**e**,**f**) drug-resistant cells. In each panel **a**-**f**, results are shown from simulations with no drugs (top row), drug 1 monotherapy (left column in each treatment panel), drug 2 monotherapy (middle column in each treatment panel), and combination therapy (right column in each treatment panel) for high (middle row) and low (bottom row) doses. Examples of initial and final simulation snapshots are shown in each panel. Drug-sensitive and drug-resistant cells are, respectively, orange and purple. The panels also include plots of the dynamic mean fraction of drug-sensitive (orange) and drug-resistant (purple) cells, and standard deviations (dashed lines) from 10 simulation runs.

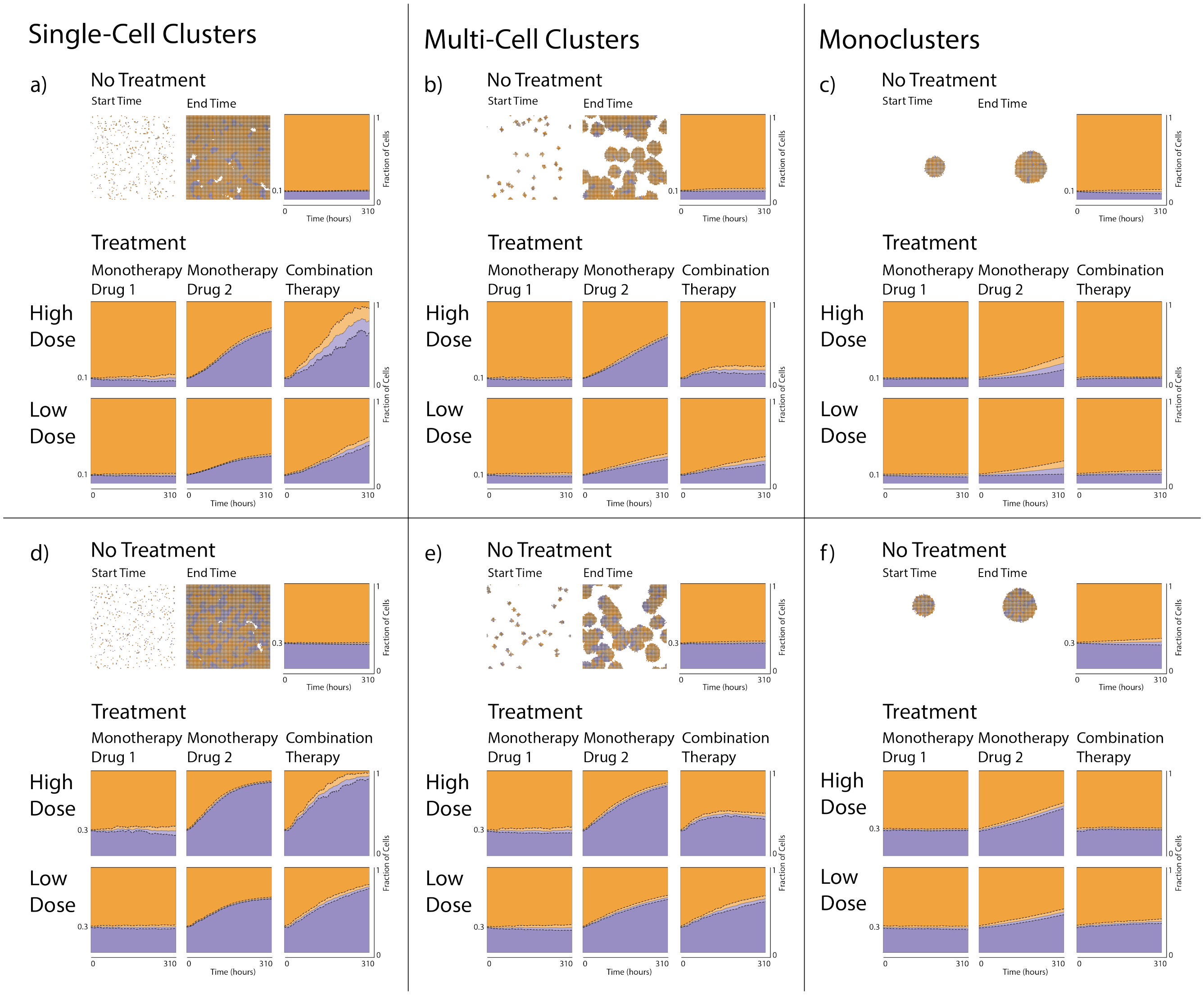

Fig. S20: **Cells that are resistant to a PARPi (drug 2) compete for spatial resources with drug-sensitive cells when daughter cells can be placed in 1st-, 2nd-, and 3rd-order neighbourhoods.** A total number of $P_0$ cells are randomly seeded on the lattice in single-cell clusters (**a**,**d**), multi-cell clusters (**b**,**e**), or monoclusters (**c**,**f**), with either $0.1P_0$ (**a**,**b**,**c**) or $0.3P_0$ (**d**,**e**,**f**) drug-resistant cells. In each panel **a**-**f**, results are shown from simulations with no drugs (top row), drug 1 monotherapy (left column in each treatment panel), drug 2 monotherapy (middle column in each treatment panel), and combination therapy (right column in each treatment panel) for high (middle row) and low (bottom row) doses. Examples of initial and final simulation snapshots are shown in each panel. Drug-sensitive and drug-resistant cells are, respectively, orange and purple. The panels also include plots of the dynamic mean fraction of drug-sensitive (orange) and drug-resistant (purple) cells, and standard deviations (dashed lines) from 10 simulation runs.

##### S7.2.2 Spatial cell structures impact the feasibility of treatment objectives

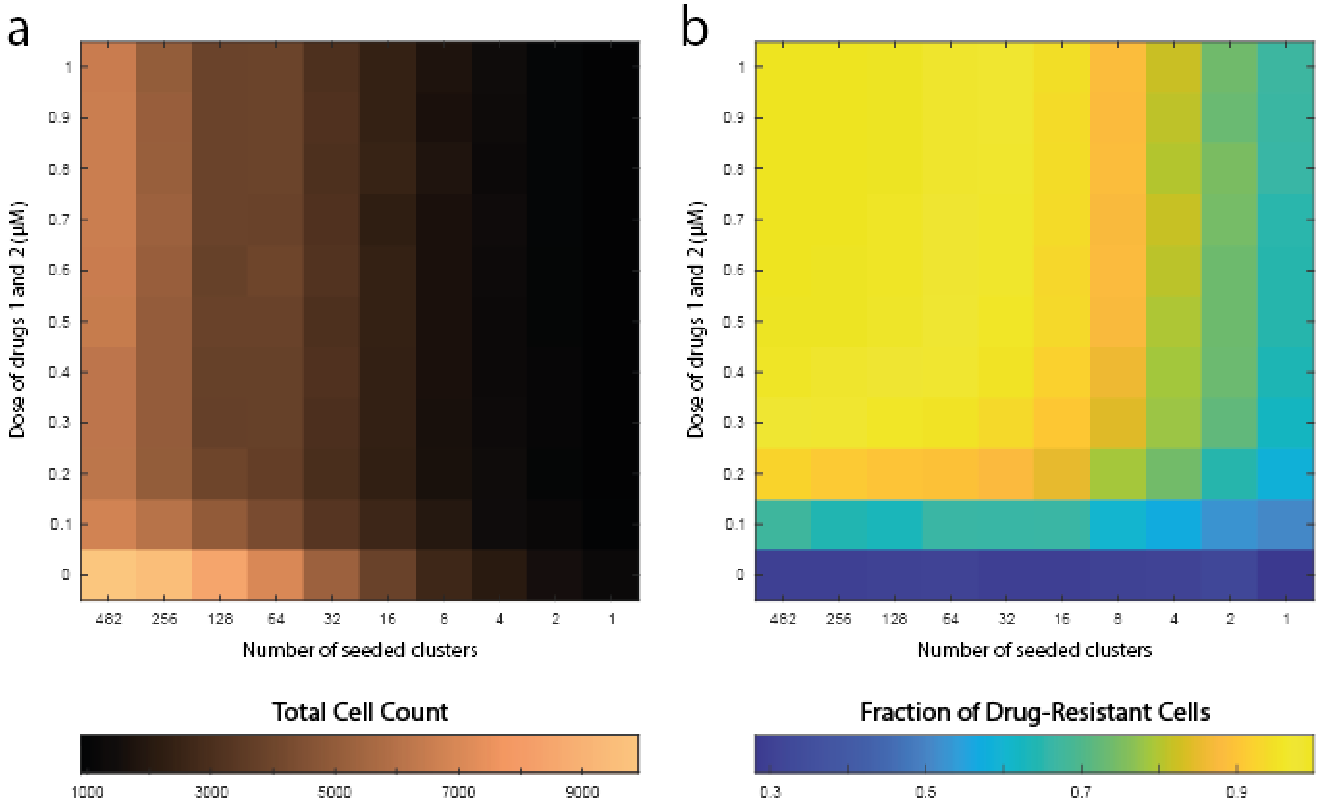

Fig. S21: **Drug doses and spatial cell configurations impact total cell counts and the composition between drug-sensitive and drug-resistant cells when daughter cells can be placed in 1st-, 2nd-, and 3rd-order neighbourhoods**. The heatmaps show results from in silico experiments in which drug-sensitive and drug-resistant cells coexist. At the start of the experiments, a total number of $P_0$ cells are seeded on the lattice, where $0.3P_0$ cells are resistant to both drugs 1 and 2. Two inputs are varied in the simulations: (1) the number of clusters in which cells are seeded (indicated by the horizontal heatmap axes), and (2) the combination treatment drug doses (indicated by the vertical heatmap axes). The results show (**a**) the total cell count and (**b**) the fraction of drug-resistant cells at the end of the simulations. Each heatmap bin shows the mean value of 10 simulation runs.

##### S7.2.3 Spatial cell structures impact doubling time associated treatment responses

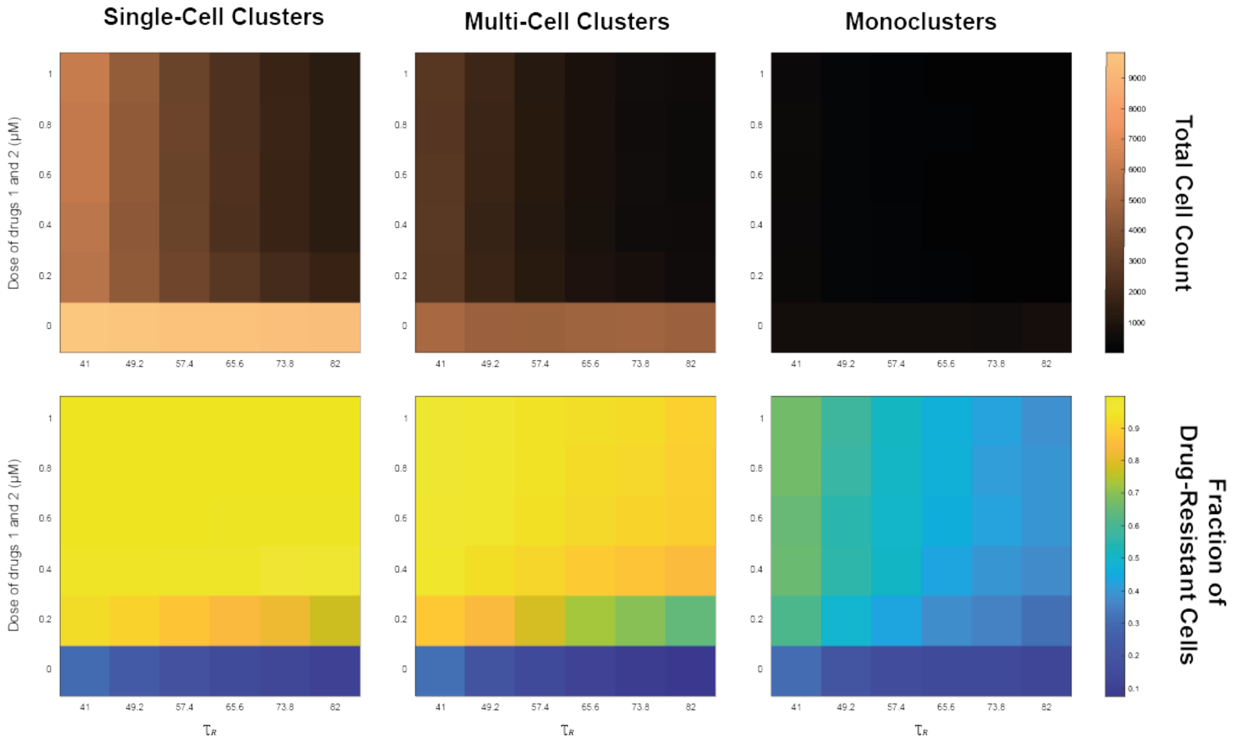

Fig. S22: **Spatial cell configurations, drug doses, and cell doubling times impact total cell counts and the composition between drug-sensitive and drug-resistant cells when daughter cells can be placed in 1st-, 2nd-, and 3rd-order neighbourhoods.** At the start of the experiments, a total number of $P_0$ cells are seeded on the lattice, where $0.3P_0$ cells are resistant to both drugs 1 and 2. Three inputs are varied in the simulations: (1) the seeded cluster configurations (left, middle, right panel), (2) the combination treatment drug doses (indicated by the vertical heatmap axes), and (3) the mean doubling time of the drug-resistant cells (indicated by the horizontal heatmap axes). The heatmaps show the total cell count (top panel) and the fraction of drug-resistant cells (bottom panel) at the end of the simulation. Each heatmap bin shows the mean value of 10 simulation runs.

#### S8 Low versus high combination treatments are zoomed in on

In the main manuscript we found that when cells are seeded in multi-cell clusters or monoclusters and populations include cells resistant to only one drug, low-dose combination treatments favour the drug-resistant population compared to high-dose combination treatments at the end of the simulations. In Fig. S23, we zoom in on Figs. 4 and 5 from the main manuscript to investigate the difference between low- and high-dose combination treatments in more detail. In Fig. S23, low-dose combination treatments (blue curves) grow a lot steeper compared to high-dose combination treatments (red curves). This is particularly noticeable when cells are resistant to the PARPi and are seeded in multi-cell clusters (Fig. S23**c**), where the low-dose combination treatments result in a significantly higher drug-resistant fraction compared to the high-dose combination treatments.

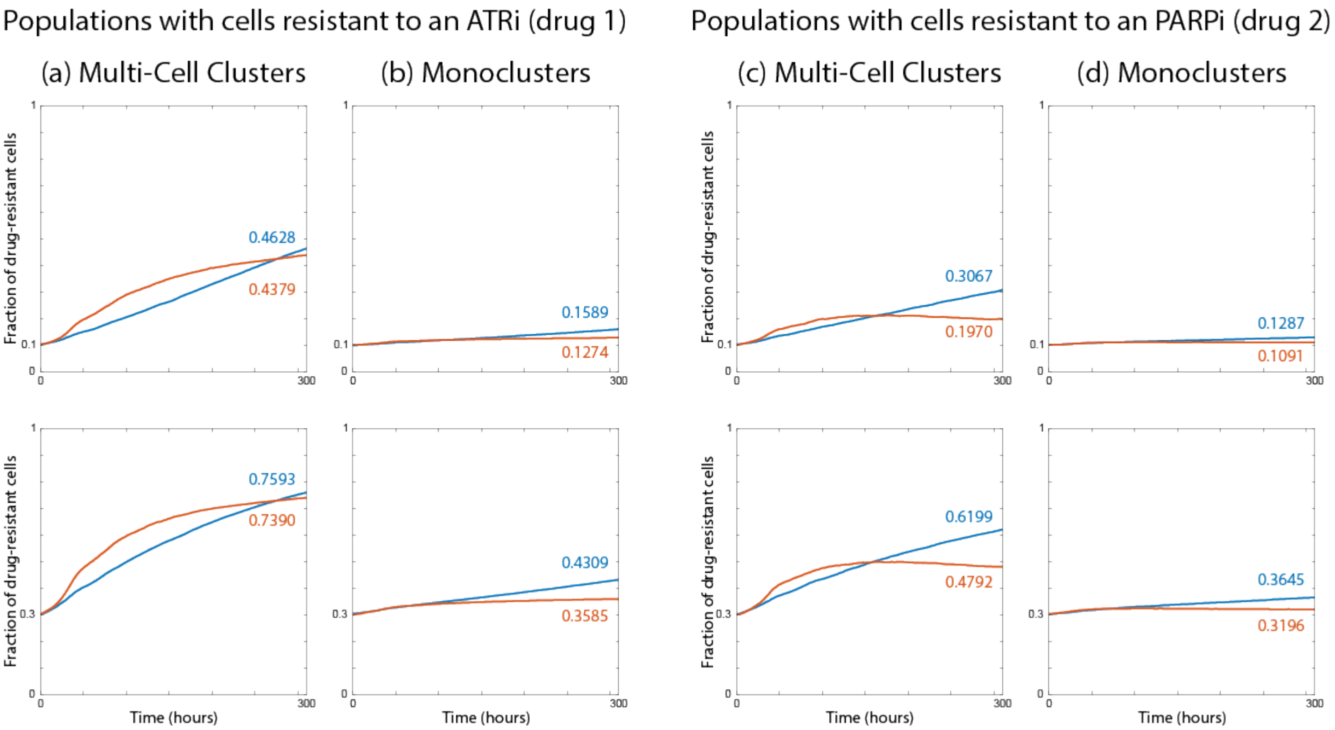

Fig. S23: **Low-dose combination therapy favours the drug-resistant population compared to high-dose combination treatments.** At the start of the experiments, a total number of $P_0$ cells are seeded on the lattice, where cells are resistant to either drug 1 (**a** and **b**) or drug 2 (**c** and **d**). The cells are seeded in multi-cell clusters (**a** and **c**) or monoclusters (**b** and **d**). The panels include plots of the dynamic mean fraction of drug-resistant cells from 100 simulation runs. The blue and red curves are low-dose and high-dose combination treatments respectively. In each panel (**a-d**) either $0.1P_0$ (top row) or $0.3P_0$ (bottom row) cells are drug-resistant. The fraction of cells at the end time of the simulations is written on each plot for low-dose (blue) and high-dose (red) combination treatment.

#### References

K. Alden, M. Read, J. Timmis, P. S. Andrews, H. Veiga-Fernandes, and M. Coles. Spartan: A comprehensive tool for understanding uncertainty in simulations of biological systems. *PLoS Comput Biol*, 9(2):e1002916, 2013. doi:10.1371/journal.pcbi.1002916.

A. Ashworth. A synthetic lethal therapeutic approach: Poly(ADP) ribose polymerase inhibitors for the treatment of cancers deficient in DNA double-strand break repair. *J Clin Oncol*, 26(22):3785–3790, 2008. doi:10.1200/JCO.2008.16.0812.

M. C. Caron, A. K. Sharma, J. O'Sullivan, L. R. Myler, M. T. Ferreira, A. Rodrigue, Y. Coulombe, C. Ethier, J. Gagné, M. Langelier, J. M. Pascal, I. J. Finkelstein, M. J. Hendzel, G. G. Poirier, and J. Masson. Poly(ADP-ribose) polymerase-1 antagonizes DNA resection at double-strand breaks. *Nat Commun*, 10(1):2954, 2019. doi:10.1038/s41467-019-10741-9.

A. R. Chaudhuri and A. Nussenzweig. The multifaceted roles of PARP1 in DNA repair and chromatin remodelling. *Nat Rev Mol Cell Biol*, 18(10):610–621, 2017. doi:10.1038/nrm.2017.53.

A. Chen. PARP inhibitors: Its role in treatment of cancer. *Chin J Cancer*, 30(7):463–471, 2011. doi:10.5732/cjc.011.10111.

J. Cohen. *Statistical power analysis for the behavioral sciences (2nd ed.)*. Routledge, 1988. doi:10.4324/9780203771587.

S. Hamis, S. Stratiev, and G. G. Powathil. Uncertainty and sensitivity analyses methods for agent-based mathematical models: An introductory review. In *The Physics of Cancer*, chapter 1, pages 1–37. World Scientific, 2021a. doi:10.1142/9789811223495_0001.

S. Hamis, J. Yates, M. A. J. Chaplain, and G. G. Powathil. Targeting cellular DNA damage responses in cancer: An in vitro-calibrated agent-based model simulating monolayer and spheroid treatment responses to ATR-inhibiting drugs. *Bull Math Biol*, 83(10):103, 2021b. doi:10.1007/s11538-021-00935-y.

K. Levenberg. A method for the solution of certain non-linear problems in least squares. *Q Appl Math*, 2(2): 164–168, 1944. doi:10.1090/QAM/10666.

R. L. Lloyd, P. W. G. Wijnhoven, A. Ramos-Montoya, Z. Wilson, G. Illuzzi, K. Falenta, G. N. Jones, N. James, C. D. Chabbert, J. Stott, E. Dean, A. Lau, and L. A. Young. Combined PARP and ATR inhibition potentiates genome instability and cell death in ATM-deficient cancer cells. *Oncogene*, 39(25):4869–4883, 2020. doi:10.1038/s41388-020-1328-y.

E. D. Lobo and J. P. Balthasar. Pharmacodynamic modeling of chemotherapeutic effects: Application of a transit compartment model to characterize methotrexate effects in vitro. *AAPS PharmSci*, 4(4):E42, 2002. doi:10.1208/ps040442.

X. Miao, G. Koch, S. Ait-Oudhia, R. M. Straubinger, and William J. Jusko. Pharmacodynamic modeling of cell cycle effects for gemcitabine and trabectedin combinations in pancreatic cancer cells. *Front Pharmacol*, 7:421, 2016a. doi:10.3389/fphar.2016.00421.

X. Miao, G. Koch, R. M. Straubinger, and W. J. Jusko. Pharmacodynamic modeling of combined chemotherapeutic effects predicts synergistic activity of gemcitabine and trabectedin in pancreatic cancer cells. *Cancer Chemother Pharmacol*, 77(1):181–193, 2016b. doi:10.1007/s00280-015-2907-4.

G. Noël, N. Giocanti, M. Fernet, F. Mégnin-Chanet, and V. Favaudon. Poly(adp-ribose) polymerase (PARP-1) is not involved in DNA double-strand break recovery. *BMC Cell Biol*, 16:4, 2003. doi:10.1186/1471-2121-4-7.

M. J. O'Connor. Targeting the DNA damage response in cancer. *Mol Cell*, 60(4):547–560, 2015. doi:10.1016/j.molcel.2015.10.040.

K. Pugh, M. Davies, and G. Powathil. A mathematical model to investigate the effects of ceralasertib and olaparib in targeting the cellular DNA damage response pathway. *J Pharmacol Exp Ther*, 387(1):55–65, 2023. doi:10.1124/jpet.122.001558.

Z. Ugray, L. Lasdon, J. Plummer, F. Glover, J. Kelly, and R. Martí. Scatter search and local NLP solvers: A multistart framework for global optimization. *INFORMS J Comput*, 19(3):328–340, 2007. doi:10.1287/ijoc.1060.0175.

A. Vargha and H.D. Delaney. A critique and improvement of the "cl" common language effect size statistics of McGraw and Wong. *J Educ Behav Stat*, 25(2):101–132, 2000. doi:10.3102/10769986025002101.
